# Osteosarcoma tumors maintain intra-tumoral transcriptional heterogeneity during bone and lung colonization

**DOI:** 10.1101/2020.11.03.367342

**Authors:** Sanjana Rajan, Emily Franz, Camille A. McAloney, Tatyana A. Vetter, Maren Cam, Amy C. Gross, Cenny Taslim, Meng Wang, Matthew V. Cannon, Alexander Oles, Ryan D. Roberts

**Author notes:** Corresponding author, **Correspondence:** Ryan D. Roberts, Division of Pediatric Hematology, Oncology, and BMT, Nationwide Children’s Hospital, 700 Children’s Drive, Columbus, OH 43205. Co-first authors (Sanjana Rajan, Emily Franz). **Email addresses:** SR, EF, CAM, TAV, MC, ACG, CT, MW, MVC, AO.

## Abstract

**Background:** Tumors are complex tissues containing collections of phenotypically diverse malignant and nonmalignant cells. We know little of the mechanisms that govern heterogeneity of tumor cells nor of the role heterogeneity plays in overcoming stresses, such as adaptation to different microenvironments. Osteosarcoma is an ideal model for studying these mechanisms—it exhibits widespread inter- and intra-tumoral heterogeneity, predictable patterns of metastasis, and a lack of clear targetable driver mutations. Understanding the processes that facilitate adaptation to primary and metastatic microenvironments could inform the development of therapeutic targeting strategies.

**Results:** We investigated single-cell RNA-sequencing profiles of 47,977 cells obtained from cell line and patient-derived xenograft models as cells adapted to growth within primary bone and metastatic lung environments. Tumor cells maintained phenotypic heterogeneity as they responded to the selective pressures imposed during bone and lung colonization. Heterogenous subsets of cells defined by distinct transcriptional profiles were maintained within bone- and lung-colonizing tumors, despite high-level selection. One prominent heterogenous feature involving glucose metabolism was clearly validated using immunofluorescence staining. Finally, using concurrent lineage tracing and single-cell transcriptomics, we found that lung colonization enriches for multiple clones with distinct transcriptional profiles that are preserved across cellular generations.

**Conclusions:** Response to environmental stressors occurs through complex and dynamic phenotypic adaptations. Heterogeneity is maintained, even in conditions that enforce clonal selection. These findings likely reflect the influences of developmental processes promoting diversification of tumor cell subpopulations, which are retained, even in the face of selective pressures.

## Background

Osteosarcoma is the most common malignant tumor of bone in pediatric patients^1^. This aggressive disease occurs in the metaphyseal region of the long bones, coinciding in both anatomic location and developmental timing with periods of rapid linear bone growth^2^. Localized disease has a favorable five-year survival rate of over 70%^3^, but survival plummets to less than 20% in the event of metastasis^4^. Unfortunately, survival rates have not changed since the introduction of chemotherapy in the 1980s^5^. Staging and therapeutic assignment remain solely based on the presence or absence of metastasis^6^. One unique feature of osteosarcoma is its extreme tropism for the lung, which is the primary site of metastasis in nearly 90% of patients^7,8^. How these tumors adapt to survive and grow as they migrate from a bone to a lung environment remains poorly understood. While others have reported that osteosarcoma metastasis is a polyclonal process^9^, the molecular pathways that mediate these adaptations during tissue colonization—which may involve a coordination of behaviors between cells from distinct clones with distinct phenotypes—remain to be studied at a single-cell level. Such investigation may provide novel insights into tumor biology and reveal vulnerabilities not evident in studies that utilize bulk methods.

Most osteosarcoma patients have disseminated tumor cells at the time of diagnosis^6,10^. Thus, studies focused on the latter steps in the metastatic process, especially lung colonization, are those most likely to produce targets that will translate into impactful clinical interventions. A large body of literature suggests that tumor cells face enormous stresses upon dissemination to the lung—only the fittest cells survive the transition from dissemination to lung colonization, while most cells die^11^. Thus, the experiments shown here focus deliberately on understanding the contributions of clonal selection and transcriptional heterogeneity to the adaptive processes that allow tumors to traverse the tissue colonization bottleneck^11–15^, using reductionist systems that are agnostic of early steps in the metastatic cascade. For instance, whether the properties that mediate fitness within the lung arise from rare, but pre-existing clones that carry an intrinsic survival advantage or from some active adaptive response to the new environment remains poorly understood. Such fundamental questions have largely remained unanswered due to technical limitations. In this study, we overcome this gap by combining single-cell transcriptomics with lineage tracing to study transcriptional heterogeneity dynamics and clonal evolution using linked single cell datasets.

We analyzed single cell transcriptomic libraries of 47,977 osteosarcoma cells obtained from cell lines and low-passage patient-derived xenograft (PDX) models that were grown as both orthotopic primary tumors and lung metastases to identify changes in intra-tumoral transcriptional heterogeneity that occur during tibia and lung colonization. While we suspected that adaptation to growth within the tibia or lung would select for a rather narrowly-defined subset of cells with transcriptional phenotypes that endowed cells with increased fitness in these distinct environments, we instead found that tumors maintain a high degree of transcriptional heterogeneity while adapting to growth in both bone and lung environments. However, while the overall transcriptional heterogeneity was maintained, the process of adaptation to these distinct environments revealed reproducible patterns of biologic sub-specialization within subsets of cells that were characteristic for each tissue niche. In one intriguing example, populations of cells expressing genes suggestive of reliance on glycolytic and aerobic metabolism both emerged during lung colonization. Using immunofluorescence, we validated this metabolic heterogeneity in both primary and metastatic mouse tumors. Furthermore, we validated the heterogeneous activation of multiple pathways identified from our preclinical models in single cell RNA-seq datasets from patient tumors. Finally, we combined lineage tracing with single cell transcriptomics to study the contribution of clonal selection to the phenotypic shifts that we observed. Surprisingly, while several clones exhibited clear expansion during lung colonization, these clones emerged within clusters spread across the entire spectrum of transcriptional profiles, rather than a particular cluster representing a genome-wide transcriptional state that endowed tissue-specific fitness. However, isolation of cells from these expanding clones revealed activation of a more focused set of genes that again suggested the importance of metabolic flexibility during lung colonization.

Based on these investigations, we conclude that osteosarcoma cells undergo dynamic changes in their transcriptional phenotype as they colonize different microenvironmental conditions, while simultaneously maintaining an overall degree of transcriptional heterogeneity in the wake of clonal selection. Overall, these data highlight the importance of understanding intra-tumor heterogeneity in the context of tumor cell transcriptional changes or adaptations during tissue colonization.

## Results

### Transcriptional heterogeneity in cell line and PDX models of osteosarcoma

Recent work has shown that osteosarcoma tumors collected directly from patients exhibit significant transcriptional heterogeneity^16,17^. However, it is unclear if this heterogeneity is maintained within cell line and PDX models of disease^6^. We investigated phenotypic heterogeneity within two *in vitro* and *in vivo* models of osteosarcoma by performing single-cell RNA sequencing (scRNA-seq). We generated transcriptomic libraries from a cell line (OS-17) and a PDX (NCH-OS-7; see ***Table 1***, supplementary ***Table S1***, for full characteristics as per PDX-MI standards) grown in cell culture or as subcutaneous flank tumors (>2,500 cells per model). To distinguish tumor cells from murine host cells, we separated cells with reads that mapped to the human genome from those that mapped to mouse. Principal component analysis identified that the majority of the variation in gene expression of tumor cells was dominated by cell cycle related genes, a common finding in proliferative malignant tissues^18^ (see supplementary ***Figure S1***). Scaling out cell-cycle related genes unmasked underlying heterogeneity that did not correspond to variation arising solely from cells being in different states of cell cycle (see supplementary ***Figure S1*, S2**A, B, and ***Figure 1***A, B). Using genes that were differentially upregulated in each cluster relative to the remaining cells in each sample, we identified pathways enriched in these subsets of cells as well as those pathways differentially downregulated in each cluster (***Figure 1***C, see supplementary ***Figure S3***A). While subsets within each model shared some enriched gene sets--for example, OS-17 cells in cluster 0 and cluster 1 showed enrichment for epithelial to mesenchymal transition (EMT) genes--none of the subsets had completely identical profiles, suggesting that these subsets are indeed transcriptionally distinct. Interestingly, while EMT transcription factors play an important role in promotion of osteosarcoma cell invasion and metastasis^19^, expression of EMT-related genes was associated with certain, distinct subsets. Furthermore, individual subsets had differentially expressed genes that were associated with multiple gene sets. For example, OS-17 cells in cluster 2 were enriched for E2F targets, G2M checkpoint target genes, and MYC target genes; while NCH-OS-7 cells in cluster 0 were enriched for genes associated with EMT, glycolysis, hypoxia, and MYC target genes. This identification of gene sets exclusive to specific clusters of cells validated that these cell line and PDX models demonstrate intra-tumor phenotypic heterogeneity.

**Table 1.**
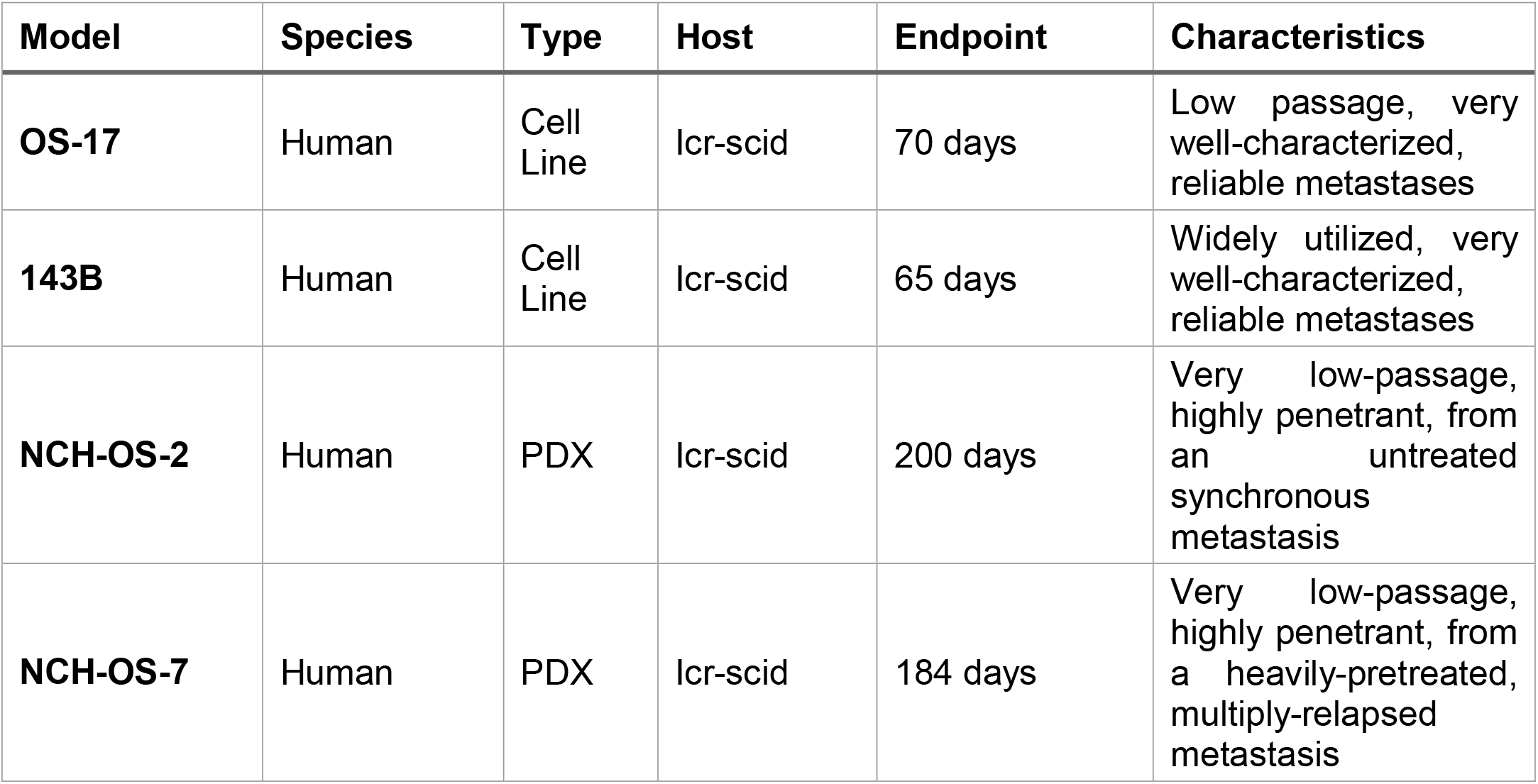
Characteristics of osteosarcoma models used in this study.

**Figure 1.**
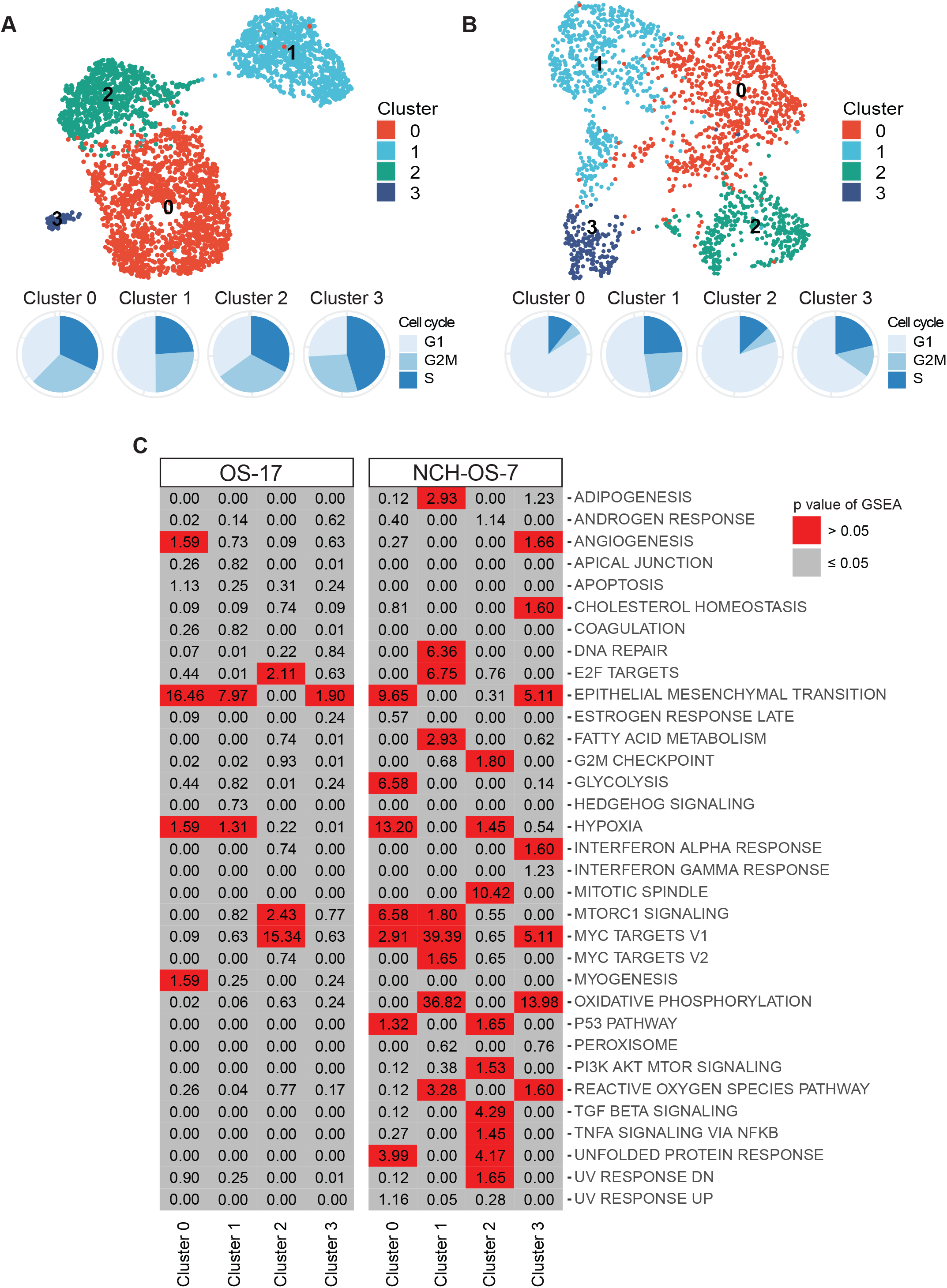
Cell line and PDX models of osteosarcoma display transcriptional heterogeneity. A) UMAP analysis of OS-17 cells (n = 3327) grown in cell culture. Cell cycle distribution of cells (n = 3327) is visualized as pie charts. B) UMAP analysis of NCH-OS-7 cells (n = 1998) grown as subcutaneous flank tumor. Cell cycle distribution of cells is visualized as pie charts. C) Pathway enrichment analysis for hallmark gene sets associated with genes upregulated in distinct clusters identified in the two osteosarcoma models. P values were adjusted for multiple comparisons. Boxes in grey identify non-significant enrichments, whereas boxes in red identify statistically significant enrichments. MTORC1: mechanistic target of rapamycin (mTOR) complex 1. PI3K: Phosphoinositide 3-kinase. AKT: Protein kinase B. TGF: Transforming growth-factor. TNF: tumor necrosis factor. NFKB: Nuclear Factor kappa-light-chain-enhancer of activated B cells.

### Transcriptional signatures associated with colonization of bone and lung microenvironments

Microenvironment cues play a critical role in shaping transcriptional plasticity^20^. We hypothesized that transcriptional signatures shared across models upon colonization of tibia or lung microenvironments will inform characteristics that are microenvironment-specific rather than tumor-specific. Single cell suspensions of two cell lines and two PDXs were injected intratibially or intra-venously to generate orthotopic models of bone-colonizing and lung-colonizing lesions (***Figure 2***A schematic). Both the PDXs included in this study (NCH-OS-2, NCH-OS-7) are early passage (passage 3, passage 5) and were obtained from patient lung metastases.

**Figure 2.**
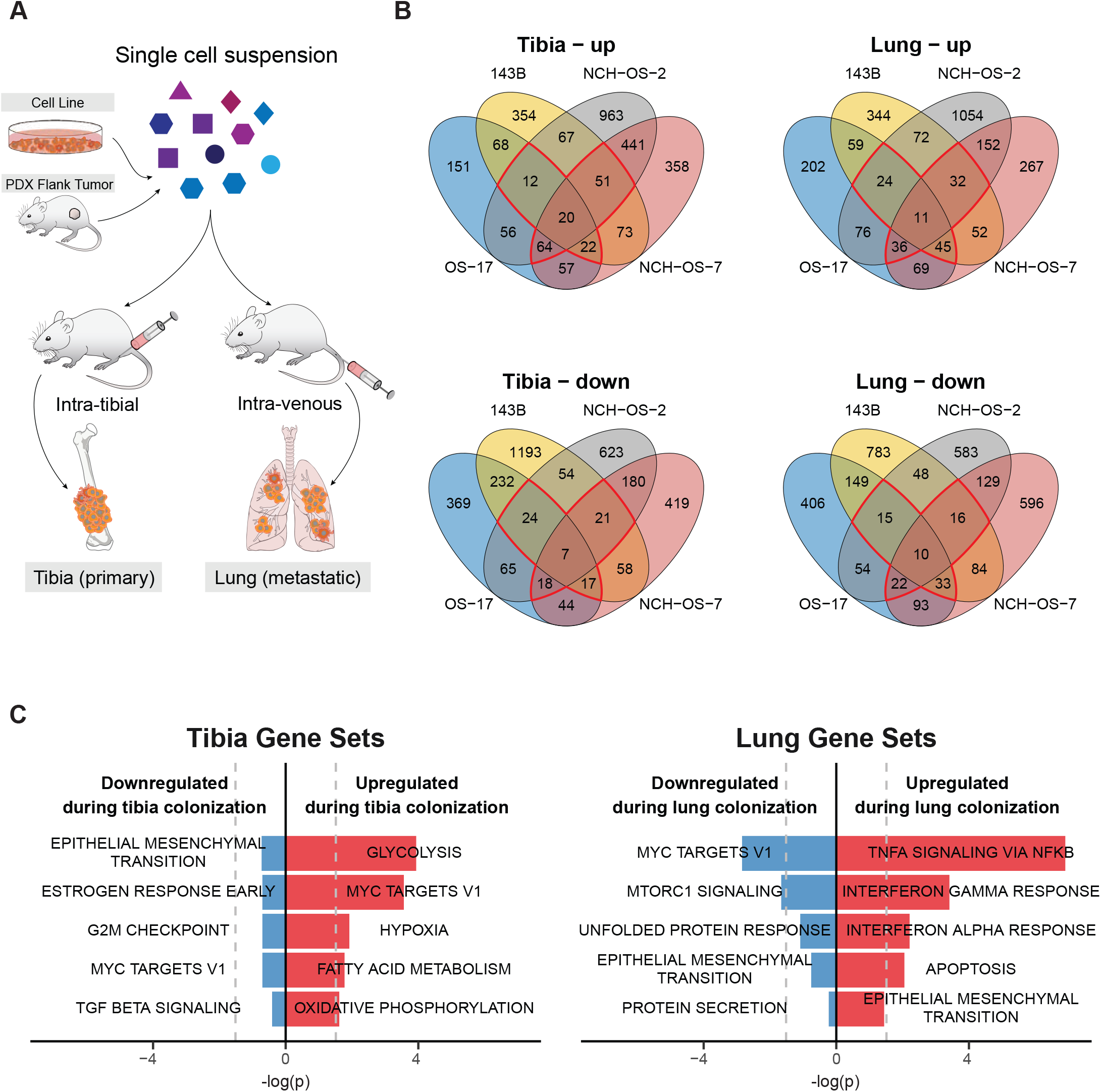
Osteosarcoma cells adopt distinct transcriptional profiles as they colonize tibia and lung microenvironments. A) Schematic of study design depicting generation of orthotopic tibia- and lung-colonizing tumors. Tumors were harvested for scRNA-seq when mice reached endpoint. B) Venn diagrams showing overlap of differentially expressed genes that are up- or down-regulated in tibia or lung lesions relative to corresponding starting population of cells (cell culture or PDX flank tumors). Regions outlined in red identify differentially expressed genes shared across at least three models. C) Pathway enrichment analysis with adjusted p values for hallmark gene sets associated with these shared genes.

To study the transcriptional profiles associated with tibia and lung colonization, we performed a differential expression analysis of the pseudo-bulked scRNA-seq data from bone-colonizing tumors or lung-colonizing lesions relative to their corresponding starting population (cell culture/flank tumors). Most of the differentially regulated genes were unique to each model, with only 0.3-0.8% of genes shared across all four models (***Figure 2***B). Using genes that were either up- or down-regulated in at least three of the four models (2.7-5.6% of genes), we performed pathway enrichment analysis to identify pathways associated with colonization of tibia or lung tissues (***Figure 2***C). We verified the shared pathways between these human osteosarcoma samples and mouse osteosarcoma models (K7M2, F420) to determine shared pathways between xenograft and syngeneic models (see supplementary ***Figure S4***). Genes that were significantly upregulated upon tibia colonization included those associated with glycolysis, hypoxia, MYC targets, fatty acid metabolism, and oxidative phosphorylation. None of the identified downregulated gene sets were statistically significant. Gene sets that were most significantly upregulated with lung colonization included those associated with TNFα signaling via NFκB and EMT, whereas those significantly downregulated included MYC targets and MTORC1 signaling. Overall, we identified common pathways that were differentially regulated across the four models upon tibia or lung colonization.

### Transcriptional heterogeneity of individual cells within a tumor

We analyzed single cell transcriptional profiles of the above datasets using our scRNA-seq bioinformatics workflow to study the role of intra-tumor heterogeneity in tissue colonization (***Figure 3***A). To estimate the degree of intra-tumor heterogeneity in our models, we computed intra-tumor heterogeneity scores (ITH scores) from the average Euclidian distance between each cell and every other cell in the same dataset^18^. In essence, this is a measure of how similar any particular cell is to the other cells within a sample. Since osteosarcoma can arise from transformed progenitor cells with osteoblastic differentiation and osteoid production^6^, we compared the heterogeneity scores in osteosarcoma cells relative to human primary osteoblast cells grown in cell culture. We found that most osteosarcoma samples exhibit somewhat higher ITH scores than do normal osteoblast cultures (p <0.001, ***Figure 3***B), though there remained a high degree of overlap in the distributions of phenotypic similarity (ITH scores) within the populations of all three cell cultures. Indeed, cells from osteoblast cultures exhibit several distinct phenotypes that readily cluster away from each other, consistent with previous reports^21^ (see supplementary ***Figure S5***) and similar to that observed in the osteosarcoma samples. We have included overlap scores^22^ in the figure, which gives a more appropriate statistical assessment of the proposed hypothesis than a p value, given the type of data^23^ (i.e., we are not evaluating whether two populations containing thousands of cells are identical, rather evaluating the degree of similarity vs difference in their ITH Score distribution).

**Figure 3.**
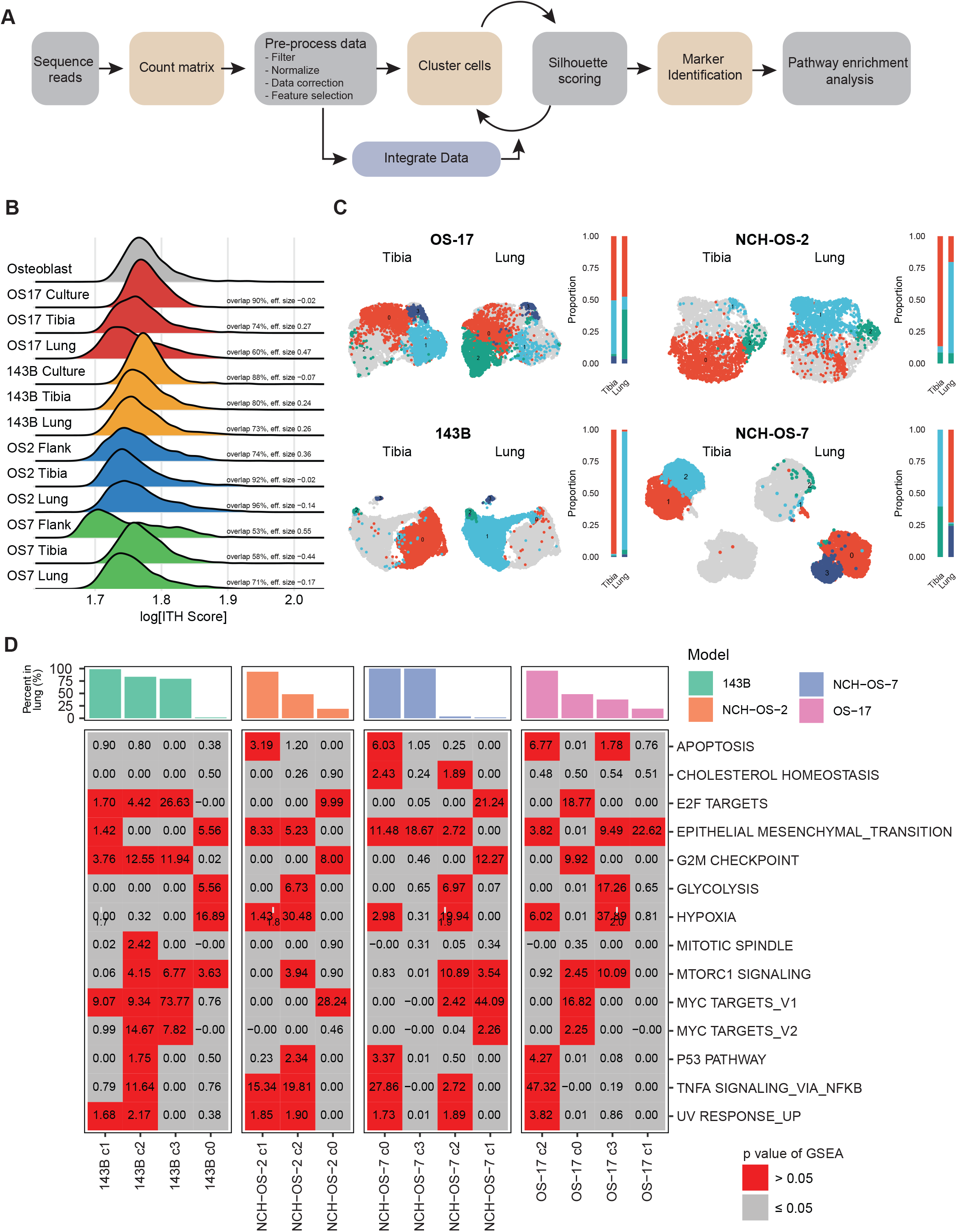
Osteosarcoma cells retain phenotypic heterogeneity despite adaptive changes in response to changing microenvironments. A) Schematic outlining scRNA-seq bioinformatics workflow. B) Osteosarcoma cells maintained overall heterogeneity with a high degree of overlap between conditions. The ridge plot shows ITH scores, which represent the gene expression “distance” between each tumor cell within a sample and all of the other tumor cells from that same sample. The overlap statistic describes the total percentage of overlap in the observed distributions between two samples. C) UMAP analysis for merged tibia- and lung-colonizing tumor samples in each of the four models. Cells in grey represent remaining cells in merged sample. Cluster enrichment analysis shows distribution of cells in each cluster in the two microenvironment conditions (tibia, lung). While some cells in the tibia and lung lesions adopted shared phenotypes, others adopted distinct phenotypes. D) Metabolic heterogeneity in glycolysis activation. We used a pathway enrichment analysis for hallmark gene sets using genes differentially upregulated in each cluster relative to every other cluster within the same model. P values were adjusted for multiple comparisons. Boxes in grey identify non-significant pathway enrichments, whereas boxes in red identify statistically significant enrichments. Bar plot shows percentage of cells identified per cluster in lung lesions. In B-D, samples subset to equal number of cells to allow inter-model comparison (n=1500 per condition).

We hypothesized that adaptation to distinct microenvironmental conditions would create selective pressures that would cause a narrowing of the transcriptional phenotypes of the tumor cells as the process would select for cells expressing genes that endow them with an environment-specific fitness phenotype. Surprisingly, we found that osteosarcoma cells grown in tibia and lung microenvironments showed no or little decrease in their ITH score distributions relative to those grown *in vitro* or as flank tumors (***Figure 3***B, see supplementary ***Figure S6***A), though they do show a broadening of the cell-cell similarity distribution, with small populations of cells emerging that have increasingly distinct phenotypes (higher ITH scores). These results suggest that osteosarcoma cells are at least as transcriptionally heterogeneous as non-malignant osteoblast cells and hint that adaptation to new tissues might drive the emergence of minor subpopulations. At a high level, these data support the concept that these cells retain the programs of the tissues they derive from, and that phenotypic diversity (as defined by transcriptional heterogeneity) may be a fundamental property that is retained as they colonize the tibia and lung microenvironments.

To determine if bone-colonizing tumors and lung-colonizing lesions display similar phenotypes, we attempted to use an integrative approach across models (see supplementary ***Figure S7***). However, cells predominantly clustered by tumor model of origin, a phenomenon reported in other cancers including small cell lung cancers^18^. This indicates a high degree of inter-tumor heterogeneity, a well-known characteristic of osteosarcoma tumors^24^. We then compared bone-colonizing and lung-colonizing lesions within the same model (OS-17) and characterized the transcriptional profile of each cluster (see supplementary ***Figure S8, S2***C-F). Using a cluster distribution analysis, we calculated the relative percentages of cells in each cluster to compare cells that colonized the tibia or the lung (see supplementary ***Table S2***). We identified a significant overlap in the transcriptional profiles of cells that colonized the two microenvironments, while certain cells demonstrated profiles specific to each microenvironment (***Figure 3***C, see supplementary ***Figure S6***B). For instance, both primary and metastatic OS-17 lesions had cells that exhibited the phenotype associated with cluster 0 (47% in tibia and 42% in lung), however, cells with the phenotype associated with cluster 2 were significantly enriched in metastatic lesions (1% in tibia and 40% in lung). Using the pathway enrichment analysis outlined in ***Figure 1***, we investigated the transcriptional profiles associated with each cluster (***Figure 3***D, see supplementary ***Figure S3***B, ***S6***C). This cluster enrichment-based strategy allowed us to overcome the challenges presented by inter-tumoral heterogeneity and instead focus on the characteristics of subsets that show differential fitness within the bone and lung microenvironments. Pathways enriched in each cluster were unique, with high heterogeneity in the activation of individual pathways across clusters within the same tumor (see supplementary ***Figure S9 - Figure S18***). For example, while OS-17 cells in cluster 0 and cluster 3 were associated with MTORC1 signaling, only cells in cluster 3 were enriched for glycolysis-related genes. Our results suggested that the maintenance of subpopulations with distinct phenotypes is reproducible and likely important for the biology of lung colonization. To ensure that these results were indicative of generalizable properties of osteosarcoma tumors (and not simply an artifact of our chosen model system), we validated the presence of these intratumoral subsets defined by glycolysis, hypoxia, epithelial-mesenchymal transition, and TNFa signaling via NFKB within scRNA-seq datasets generated directly from patient tumors (see supplementary ***Figure S19 - Figure S28***). Our results suggest that colonization of lung tissue requires contributions from tumor cell subsets exhibiting distinct metabolic behaviors.

### Metabolic heterogeneity in osteosarcoma tumors

To validate our observations suggesting changes in gene expression related to metabolic heterogeneity at the protein level, we performed immunofluorescence staining for a known activation marker of glycolysis, glucose transporter 1 (GLUT1, a primary component of the observed gene signature), in lung lesions from two models (OS-17 and 143B). Sections were co-stained for vimentin to distinguish osteosarcoma cells from lung parenchyma. We observed variable staining for GLUT1 protein within individual lung lesions, proving that the observed heterogeneity is relevant at the protein level and not an artifact resulting from collections of homogenous, spatially isolated lesions that stained differently from one another (***Figure 4***A, B). Interestingly, this heterogeneity was evident even in the smallest lesions, suggesting that this metabolic heterogeneity either represents a fundamental property intrinsic to subpopulations when exposed to the lung environment or provides some survival and/or growth advantage for metastatic lung lesions.

**Figure 4.**
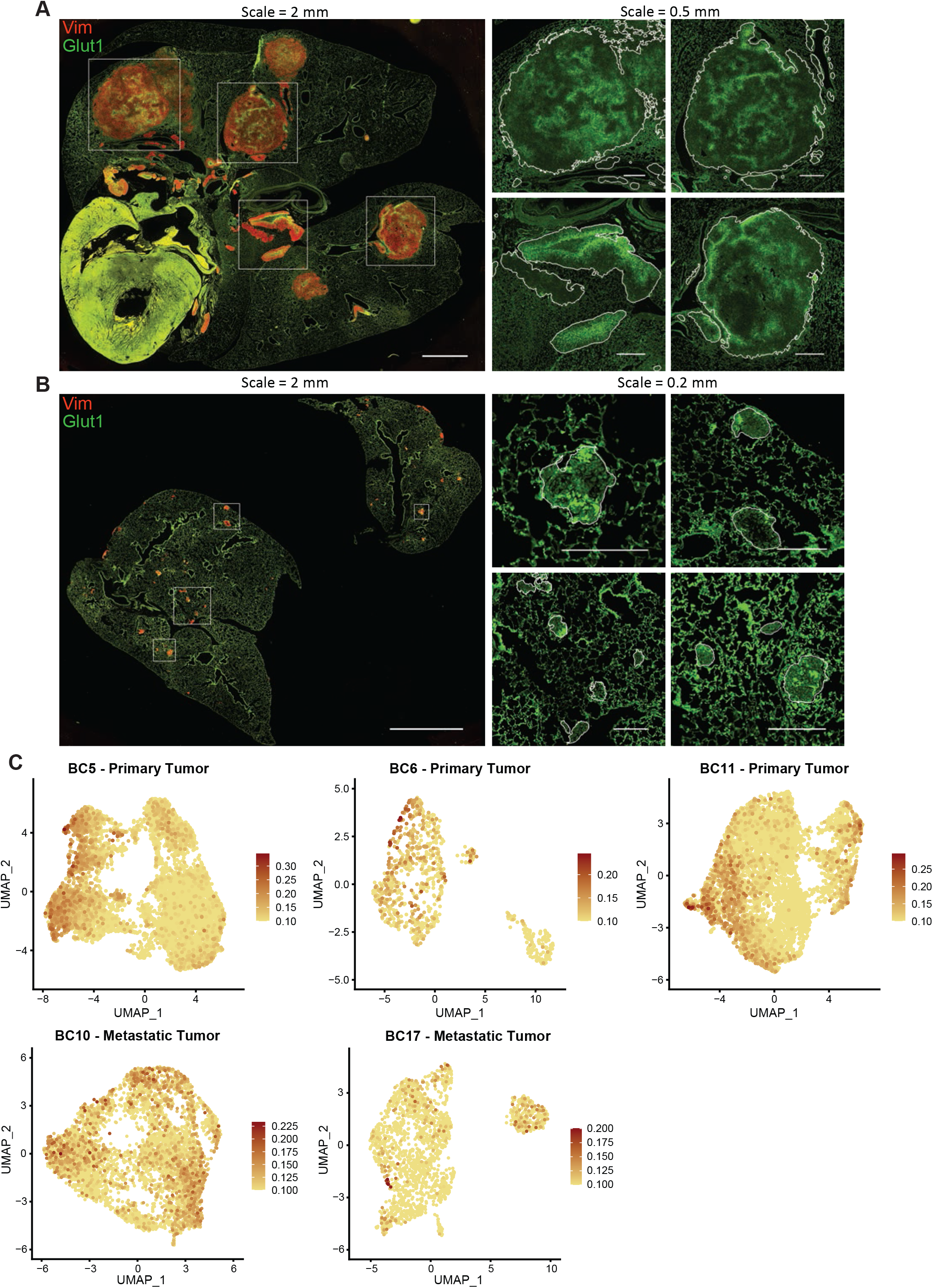
Heterogeneity in Glycolysis activation identified within lung-colonizing osteosarcoma lesions. A, B) Immunofluorescence staining of mouse lungs bearing OS-17 and 143B lung-colonizing tumors, respectively, for GLUT1 (green; a marker of glycolysis) and vimentin (red; marker to identify osteosarcoma cells). Magnified GLUT1 staining is shown for the boxed regions on the whole-section images. Lesion edges are indicated by white outlines in the magnified regions. Tumors showed a high degree of intra-tumor variation in GLUT1 staining intensity. C) FeaturePlots for the Glycolysis module score in patient primary tumor datasets. The module score for Glycolysis was calculated using the ‘AddModuleScore’ function in Seurat with msigdb HALLMARK_GLYCOLYSIS genes as input features for expression program.

To confirm that the heterogeneity observed within this representative pathway results from activation of gene expression modules within the cells and not simply from stochastic variations in gene expression, we stained for other members of the glycolysis gene set (hexokinase 2– HK2, carbonic anhydrase 9–CA9, monocarboxylate transporter 4–MCT4)^25^ and a known upstream regulator of glycolysis, MYC^26^ to evaluate the degree of concordance within individual cells. We measured the intensity of GLUT1 fluorescence of individual pixels, along with fluorescence of other markers, within areas positive for vimentin. We noted a high correlation in staining for GLUT1 and the other markers within 143B and OS-17 metastatic lesions (see supplementary ***Figure S29***). Immunofluorescence staining for markers of glycolysis in bone tumors from these models identified a high degree of correlation for high, low, and intermediate staining for all of these targets (see supplementary ***Figure S29A, Figure S30***). The same staining of these cells in culture indicated that heterogeneous expression is present in culture (see supplementary ***Figure S31***). This staining validated our sequencing-based finding of heterogeneity in glycolysis activation at the protein level and confirmed that this heterogeneity was not the result of stochastic gene expression changes.

### Tumor cell clones with diverse transcriptional phenotypes expand during lung colonization

To evaluate the contributions of clonal selection to phenotypic adaptation and heterogeneity as osteosarcoma cells colonize tibia and lung microenvironments, we combined expressed-tag lineage tracing with single cell transcriptomics (***A***, B) Immunofluorescence staining of mouse lungs bearing OS-17 and 143B lung-colonizing tumors, respectively, for GLUT1 (green; a marker of glycolysis) and vimentin (red; marker to identify osteosarcoma cells). Magnified GLUT1 staining is shown for the boxed regions on the whole-section images. Lesion edges are indicated by white outlines in the magnified regions. Tumors showed a high degree of intra-tumor variation in GLUT1 staining intensity. C) FeaturePlots for the Glycolysis module score in patient primary tumor datasets. The module score for Glycolysis was calculated using the ‘AddModuleScore’ function in Seurat with msigdb HALLMARK_GLYCOLYSIS genes as input features for expression program.

***Figure 5***A). Lentivirus infection introduced a very high diversity, heritable, and 3’ capture compatible lineage tag library in osteosarcoma cells in culture. After reserving a representative sample of the uniquely tagged cells, we generated bone and lung lesions. Single cell libraries of tibia tumors, lung tumors and cells in culture were processed bioinformatically to overlay phenotype and lineage information for each cell, and cluster transcriptional profiles were determined (see supplementary ***Figure S2***G). At the level of phenotype, we noted that cells in tibia and lung cluster away from cells in cell culture, suggesting tumor-level adaptation to tibia or lung microenvironments (***A***, B) Immunofluorescence staining of mouse lungs bearing OS-17 and 143B lung-colonizing tumors, respectively, for GLUT1 (green; a marker of glycolysis) and vimentin (red; marker to identify osteosarcoma cells). Magnified GLUT1 staining is shown for the boxed regions on the whole-section images. Lesion edges are indicated by white outlines in the magnified regions. Tumors showed a high degree of intra-tumor variation in GLUT1 staining intensity. C) FeaturePlots for the Glycolysis module score in patient primary tumor datasets. The module score for Glycolysis was calculated using the ‘AddModuleScore’ function in Seurat with msigdb HALLMARK_GLYCOLYSIS genes as input features for expression program.

**Figure 5.**
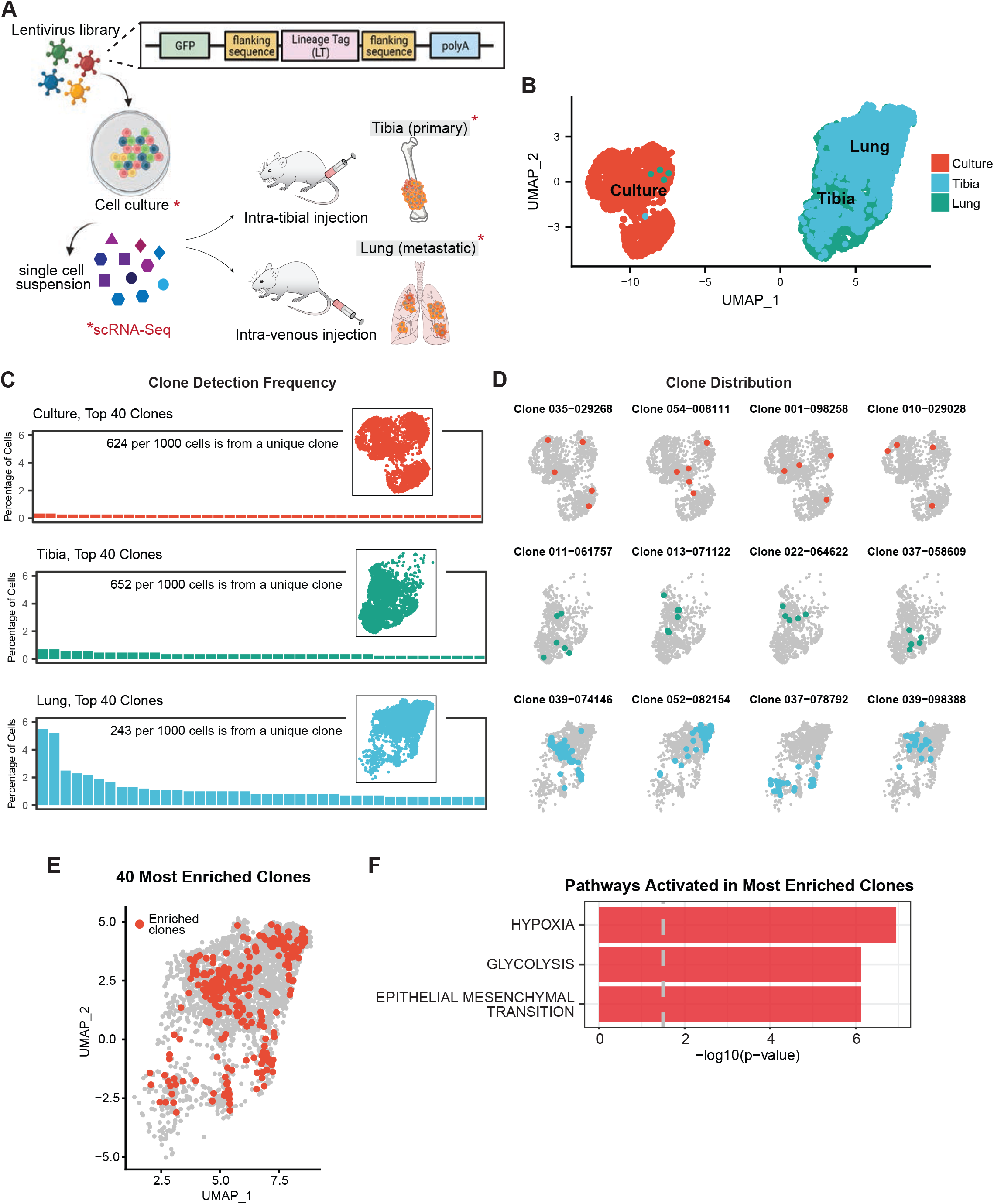
Lung colonization bottleneck enriches for clonal populations with a common transcriptional phenotype. A) Schematic of experimental set-up. B) UMAP analysis of lineage tagged OS-17 cells in culture, implanted into tibia, or inoculated to generate lung-colonizing tumors. These lineage tags are heritable, allowing us to track the transcriptional profiles of daughter cells originating from the same ancestor. Lung- and tibia-colonizing tumor samples share many cells with similar phenotypes, though cultured cells are dramatically different (n = 2800 per condition). C) Frequency distribution of clones identified in each of the conditions. A high level of clonal diversity is maintained in the tibia-colonizing tumors (78% unique clones in tibia-colonizing tumor compared to 86% unique clones in cell culture), while dominant clones emerge in the lung (only 33% unique clones). D) Overlay of cells sharing a common ancestor onto the dimensional reduction plots in the top four enriched clonal families. E) UMAP visualization of cells belonging to top ten enriched clonal families highlighted in red from the lung-colonizing tumor sample. F) Pathway enrichment analysis for hallmark gene sets comparing enriched clones relative to the remaining lung-colonizing tumor cells identifies hypoxia, glycolysis, and EMT signaling related genes to be significantly upregulated.

***Figure 5***B). Since the lineage tags are passed from parent to progeny as each cell divides, we used a frequency distribution analysis for lineage tags in the tibia and lung tumors to determine if tibia or lung colonization drove enrichment for specific clonal populations. Lineage barcode enrichment analysis for cells in cell culture identified 85% of cells to have a unique lineage tag, confirming that our starting population received lineage tags at high diversity. Surprisingly, cells from orthotopic primary tumors showed very little evidence of clonal selection, suggesting that bone represents a relatively permissive colonizing environment (***A***, B) Immunofluorescence staining of mouse lungs bearing OS-17 and 143B lung-colonizing tumors, respectively, for GLUT1 (green; a marker of glycolysis) and vimentin (red; marker to identify osteosarcoma cells). Magnified GLUT1 staining is shown for the boxed regions on the whole-section images. Lesion edges are indicated by white outlines in the magnified regions. Tumors showed a high degree of intra-tumor variation in GLUT1 staining intensity. C) FeaturePlots for the Glycolysis module score in patient primary tumor datasets. The module score for Glycolysis was calculated using the ‘AddModuleScore’ function in Seurat with msigdb HALLMARK_GLYCOLYSIS genes as input features for expression program.

***Figure 5***C). In contrast, cells from lung lesions demonstrated a higher level of enrichment for specific barcode, suggesting either an environment selecting for clones with specific properties or the activation of programs driving proliferation within a subset of tumor cells (***A***, B) Immunofluorescence staining of mouse lungs bearing OS-17 and 143B lung-colonizing tumors, respectively, for GLUT1 (green; a marker of glycolysis) and vimentin (red; marker to identify osteosarcoma cells). Magnified GLUT1 staining is shown for the boxed regions on the whole-section images. Lesion edges are indicated by white outlines in the magnified regions. Tumors showed a high degree of intra-tumor variation in GLUT1 staining intensity. C) FeaturePlots for the Glycolysis module score in patient primary tumor datasets. The module score for Glycolysis was calculated using the ‘AddModuleScore’ function in Seurat with msigdb HALLMARK_GLYCOLYSIS genes as input features for expression program.

***Figure 5***C). We confirmed that this pattern of clonal selection was reproducible in two independent replicates (see supplementary ***Figure S32****A*; supplementary ***Table S4*** *and* ***Table S5***, for lentivirus distribution in cell culture). Importantly, datasets from OS-17 cells from distinct biological replicates (different lot of cells, different lot of virus, different litter of mice) overlaid completely, even without batch correction, suggesting that intra-tumor transcriptional heterogeneity of these osteosarcoma cells is not random or stochastic and that our decision to avoid using batch correction techniques is biologically appropriate (see supplementary ***Figure S32****B*).

We then overlaid lineage information on phenotypic profiles to identify the transcriptional phenotypes most permissive to clonal expansion in the lung. Interestingly, while we found that cells within given enriched clonal families clustered together, suggesting that overarching transcriptional states were passed from parent to progeny, we also found that these expanding clones emerged from parents exhibiting phenotypes across the transcriptional spectrum (***A***, B) Immunofluorescence staining of mouse lungs bearing OS-17 and 143B lung-colonizing tumors, respectively, for GLUT1 (green; a marker of glycolysis) and vimentin (red; marker to identify osteosarcoma cells). Magnified GLUT1 staining is shown for the boxed regions on the whole-section images. Lesion edges are indicated by white outlines in the magnified regions. Tumors showed a high degree of intra-tumor variation in GLUT1 staining intensity. C) FeaturePlots for the Glycolysis module score in patient primary tumor datasets. The module score for Glycolysis was calculated using the ‘AddModuleScore’ function in Seurat with msigdb HALLMARK_GLYCOLYSIS genes as input features for expression program.

***Figure 5***D, see supplementary ***Figure S33***). This suggested that the process leading to clonal expansion might come from the triggering of specific gene programs rather than the selection of cells with any particularly fit phenotype. To identify gene programs that might be activated within these expanding clones, we performed pathway enrichment analysis using the genes differentially upregulated in the top 10 expanded clones. This analysis showed that genes associated with hypoxia, glycolysis, EMT and MTORC1 signaling were upregulated within the expanding clones, suggesting a potential mechanism for the metabolic heterogeneity noted earlier, whereas differentially downregulated genes were associated with interferon response pathways (***A***, B) Immunofluorescence staining of mouse lungs bearing OS-17 and 143B lung-colonizing tumors, respectively, for GLUT1 (green; a marker of glycolysis) and vimentin (red; marker to identify osteosarcoma cells). Magnified GLUT1 staining is shown for the boxed regions on the whole-section images. Lesion edges are indicated by white outlines in the magnified regions. Tumors showed a high degree of intra-tumor variation in GLUT1 staining intensity. C) FeaturePlots for the Glycolysis module score in patient primary tumor datasets. The module score for Glycolysis was calculated using the ‘AddModuleScore’ function in Seurat with msigdb HALLMARK_GLYCOLYSIS genes as input features for expression program.

***Figure 5***E-F, see ***Table S3***, for differentially upregulated genes, see supplementary ***Figure S3***C for downregulated pathways).

## Discussion

These studies provide insight into the roles that intra-tumor heterogeneity and clonal evolution play as osteosarcoma tumors adapt to bone and lung microenvironments. Using an orthotopic cell line and PDX models of osteosarcoma, we show that these tumor cells maintain transcriptional heterogeneity as they respond to changing microenvironment conditions while also plastically adapting their transcriptional profiles. We identified that tibia and lung tumors demonstrate phenotypic heterogeneity where subsets of cells upregulate genes associated with distinct pathways, including glycolysis, hypoxia, EMT, and TNFα signaling via NFκB. We validated this functional heterogeneity for the identified pathways in publicly available patient primary tumor datasets. Furthermore, we showed that intratumoral heterogeneity within at least one metabolically relevant pathway was reproducible using immunofluorescent staining for key proteins, suggesting heterogeneity in glycolytic metabolism within individual metastatic lesions, irrespective of lesion size. Finally, by combining lineage tracing and single-cell transcriptomic analysis, we showed that rapidly expanding clones emerge from cells across the transcriptional landscape, each of which adopts a growth-associated sub-phenotype, which is at least associated with the activation of pathways also associated with metabolism. Together, our studies suggest a tumor-intrinsic mechanism that allows for maintenance of phenotypic diversity, despite clonal selection, while undergoing adaptive transcriptional changes during lung colonization (***Figure 6***). These observations implicate several transcriptional programs in the process of lung metastasis, including several pathways linked to energy metabolism. Most interestingly, this data suggests potential cooperation between distinct tumor subpopulations, which could be a driver for the maintenance of heterogeneity and lung tropism. Further study will be necessary to determine the functional importance of these pathways.

**Figure 6.**
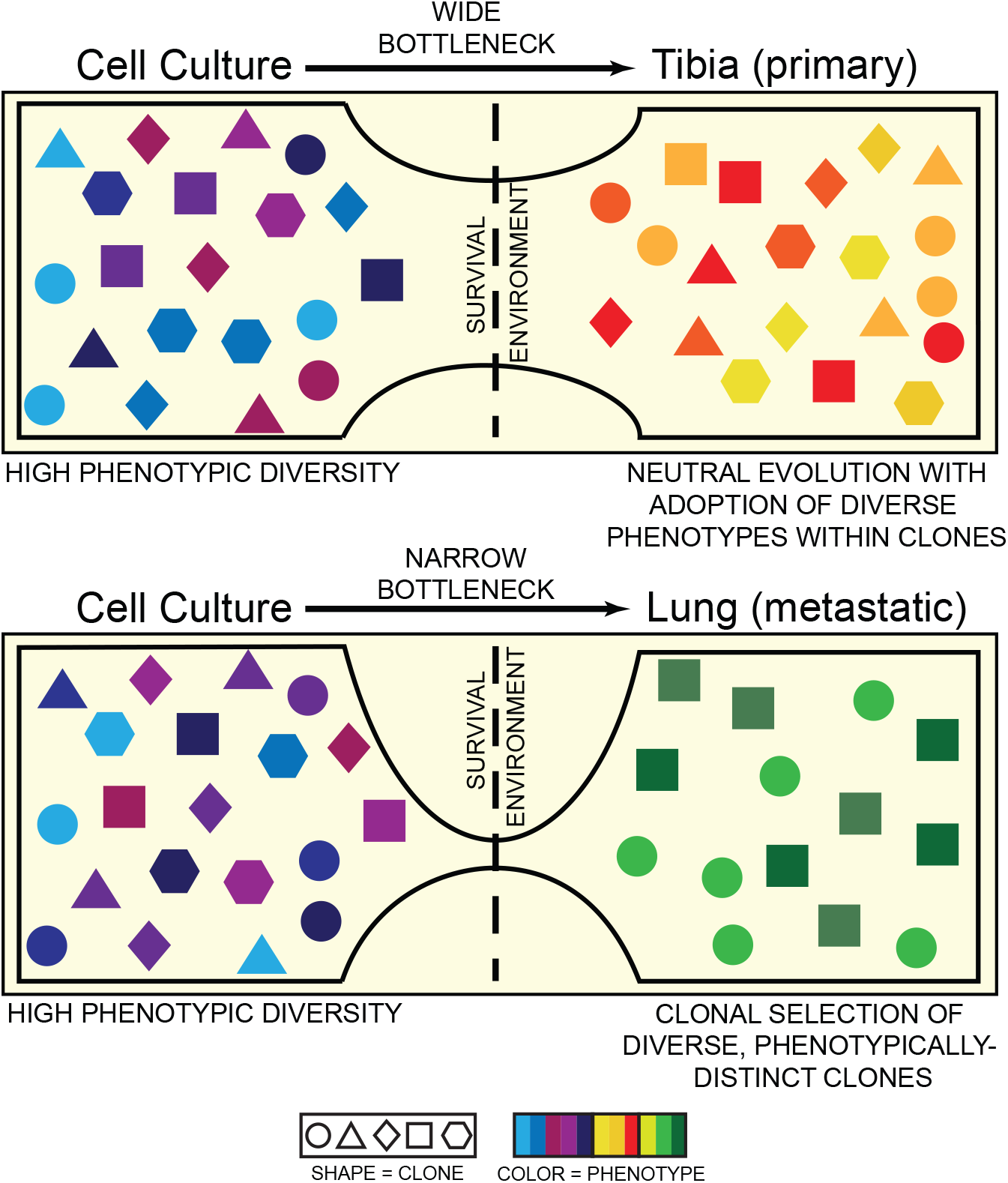
Schematic of tumor cell adaptation to changing microenvironments. Osteosarcoma cells demonstrate high transcriptional heterogeneity (colors) in cell culture, orthotopic tibia and lung microenvironments. Tumors that grow in the orthotopic tibia lesions follow dynamics of neutral evolution where different cancer clones (shapes) co-exist and propagate simultaneously. The selective pressure of lung colonization results in enrichment of clonal families with different phenotypes (colors), where cells belonging to the same family (shape) tend to have similar phenotypes (color).

Developmental processes in normal cells have been shown to require and maintain a fundamental level of phenotypic heterogeneity^27,28^. Congruently, osteoblast cells grown in cell culture demonstrated a baseline level of transcriptional heterogeneity. In our studies, osteosarcoma cells maintained an even higher degree of transcriptional heterogeneity, whether grown in mice or cell culture, and maintained that same degree of heterogeneity when exposed to selective pressures of distinct tissue environments. While more samples are needed to capture the true heterogeneity in osteosarcoma, one might speculate that the normal developmental programs that support normal bone growth are maintained (or even amplified) in tumor cells and that these cooperative mechanisms could produce a survival advantage in a transformed malignant state. Indeed, studies published using patient tumor tissues support this hypothesis^16,17^ and analysis of those published data sets identifies heterogeneity in the same pathways that we identified in our mouse models.

While pseudo-bulk analyses identified shared pathways associated with tibia and lung colonization across the models, single cell analysis identified that each tumor was composed of phenotypically distinct subsets associated with unique combinations of activated pathways. In this study, we validated findings of heterogeneity in glycolytic activation from our sequencing experiments by identifying correlated staining patterns for four distinct markers of glycolysis activation, namely GLUT1, HK2, CA9, MCT4^25^, along with a known upstream regulator of GLUT1, MYC^26^. We showed that this heterogeneity in glycolysis activation is intrinsic to individual metastatic lesions and not an artifact created by spatially isolated, metabolically distinct lesions. Strong correlation between stains for each of these five markers confirmed that the observed heterogeneity did not arise from stochastic gene expression changes. These findings also suggest that a transition to glycolytic metabolism occurs within subsets within a tumor. Reliance on glycolytic metabolism is a deeply-studied phenomenon in cancer cells, where rapid energy production can support the abnormally high needs of malignant transformation^29,30^. Our findings align with those reported in recent single cell sequencing studies in other cancers, which also showed heterogeneous activation of glycolytic pathways^31^. This may suggest that tumor cells with distinct phenotypes leverage some type of metabolic cooperation, with each cell specializing to perform a specific set of necessary tasks while relying on the activity of cells with different phenotypes for needs that they cannot meet alone. Indeed, many examples of metabolic cooperation between tumor cells and the surrounding stroma have been documented. Evidence within the remote literature hints that similar processes may be active within cultures of tumor cells^32,33^. Whatever the biologic advantage provided by this heterogeneity, these tumors clearly resemble something more like a developing tissue than a simple clonal proliferation of malignant cells. While tumors have long been described as complex organs, based on the myriad essential interactions between tumor and host cells within the microenvironment^34^, studies presented here lend support for the idea that interactions between tumor cells with distinct phenotypes may also be critical for tumor development.

Combining the power of scRNA-seq with lineage tracing allowed us to make inferences about the roles of clonal selection and transcriptional plasticity in the emergence of traits that facilitate lung colonization. We showed that lung colonization enriches for multiple clonal families, wherein clonally related cells tend to have similar transcriptional profiles. In contrast, tumor cells retained clonal diversity on tibia colonization, suggesting that this environment is relatively permissive—at least to cells previously growing in culture.

It should be noted that PDXs and patient derived cell lines may not fully recapitulate the entire spectrum of heterogeneity within a patient tumor. However, studies evaluating the rate of evolution within PDX models over time suggest that the low passage PDXs utilized in this study should retain a high degree of fidelity to the original tissue used to generate the models^35,36^.

We acknowledge that the experimental metastasis model chosen for these studies produces results that are agnostic of the multi-step process required to effect metastasis in a patient. This choice was deliberate and designed to very simply ask whether the capacity to colonize these different tissues is contained within a few rare clones with a pre-existing phenotype or whether diverse groups of cells have the capacity to adapt to stresses imposed by the tibia or lung environment. Our results suggest that both answers have some truth. The drastic shift in phenotype from cell culture or flank tumors to bone or lung tumors endorses the strong role that microenvironmental cues play in shaping tumor cell behavior.

However, lung colonization enriched for specific clonal populations that all demonstrated a similar growth-associated phenotype. Genes differentially upregulated in the top ten enriched clonal phylogenies overlapped mostly with genes related to hypoxia and glycolysis, a finding that may inform the interpretation of the staining patterns seen in our gene signature validation studies. Indeed, activation of pathways associated with hypoxia and glycolysis is closely linked and may be important in generating the energy and building blocks needed to support the rapid proliferation of these tumor cells. While our results cannot definitively show whether the enriched clones composed individual lung-colonized lesions, or whether the same clones colonized multiple lesions independently, the immunofluorescence stains for glycolysis-related gene performed earlier suggest that this metabolic heterogeneity is a phenomenon intrinsic to this process that likely occurs diffusely. Whether this transition represents a developmental step that occurs within every lesion as it begins to expand rapidly, what changes (likely epigenetic) constitute the trigger that facilitates clonal expansion, and whether the conservation of transcriptional profiles in cells that originated from a common ancestor has an underlying genetic or epigenetic basis all remain points of active study.

While the overall differential expression analysis identified genes associated with TNFα signaling via NFκB and EMT to be most significantly upregulated upon lung colonization (***Figure 2***C), the comparison between the most enriched clones and non-expanded clones identified a distinct transcriptional program associated with activation of glycolysis and hypoxia related genes. These data may suggest that lung colonization follows a two-step process—the first characterized by survival and niche establishment, with a second occurring through rapid expansion of a subset of cells that adopt a growth-associated phenotype distinct from the remaining bulk of the lung-colonizing tumor cells.

## Conclusions

Overall, we show that osteosarcoma tumors maintain a high level of intra-tumoral transcriptional heterogeneity, even while experiencing broad transcriptional changes as tumors adapt to tibia and lung microenvironments. Distinct tumors maintain subpopulations of cells characterized by activation of gene expression pathways that demonstrate similarities across models within each environment. The process of lung colonization enriches for distinct clonal populations. Interestingly, while a particular transcriptional signature appears common to rapidly expanding clonal phylogenies, these arise from groups of cells with diverse genome-wide transcriptional profiles that are maintained within each phylogeny. Additionally, maintenance of the range/degree of intra-tumor heterogeneity endorses the role of cell-autonomous mechanisms in holding up a basal evolutionary advantage associated with maintaining diverse transcriptional states^37^. How this transcriptional heterogeneity is regulated in osteosarcoma cells, and whether underlying genomic heterogeneity engenders transcriptional heterogeneity remain unanswered questions.

## Methods

### Experimental model - PDXs, Cell lines and murine studies

#### Patient derived xenografts (PDXs)

OS-17 PDX tissue was obtained from a primary femur biopsy performed at St. Jude’s Children’s Research Hospital in Memphis and was a gift from Peter Houghton^38^. Patient-derived samples, NCH-OS-2 and NCH-OS-7, were obtained from patients consented under an Institutional Review Board (IRB)-approved protocol IRB11-00478 at Nationwide Children’s Hospital (Human Subject Assurance Number 00002860).

#### Cell lines

OS-17 cells were derived from an early passage of the OS-17 PDX (see above). These cells were grown in RPMI (Corning, 10-040-CV) supplemented with 10% fetal bovine serum (FBS). 143B cells were obtained from the American Type Culture Collection (ATCC, CRL-8303) and grown in DMEM (Corning, 10-013-CV) supplemented with 10% FBS. Human osteoblast cells (hFOB 1.19) were obtained from the American Type Culture Collection (ATCC, CRL-11372) and grown in DMEM/F12 1:1 (Gibco, 21041-025) supplemented with 0.3 mg/ml G418 and 10% FBS at 34°C. All cell lines were tested annually for both STR genotyping and mycoplasma contamination using testing services provided by Genetica.

#### Murine Studies

All animal studies were approved by Nationwide Children’s Hospital Institutional Animal Care and Use Committee (IACUC protocols AR15-00022 and AR14-00045). *Flank tumors*. Cryopreserved viable tissue pieces from OS-17, 143B, NCH-OS-2 and NCH-OS-7 PDX tumors were placed in the right flank of recipient C.B-17/IcrHsd-Prkdc^scid^ mice (Envigo, Frederick, MD). These subcutaneous tumors were allowed to grow to 300 mm^3^ before excision. These were then prepped for single-cell RNA-seq. *Orthotopic primary tumors*. Single cell suspensions of 5×10^5^ cells of OS-17, 143B, NCH-OS-2 and NCH-OS-7 were injected intratibially in C.B-17/IcrHsd-Prkdc^scid^ mice as per IACUC guidelines. Primary tumors were excised once they grew to 800 mm^3^ and prepped for single-cell RNA-seq. *Experimental metastasis*. Single cell suspensions of 1×10^6^ cells of OS-17, 143B, NCH-OS-2 and NCH-OS-7 were injected intra-venously in C.B-17/IcrHsd-Prkdc^scid^ mice. Lungs were harvested once these mice reached endpoint. Endpoint criteria for euthanasia was defined as weight loss of >10% or a body condition score (BCS) of <10. For fluorescence immunohistochemistry studies, tumors were placed in 10% formalin for 24 hours at 4°C, then moved to (phosphate-buffered saline) PBS for at least one hour at 4°C. Tissues were then placed in tissue cassettes and placed in fresh PBS. Tissues were embedded in paraffin.

### Fluorescent immunohistochemistry

Primary antibodies against glucose transporter 1 (Abcam, ab15309, 1:200), hexokinase 2 (Abcam, [3D3] ab104836, 1:10), carbonic anhydrase 9 (Abcam, [2D3] ab107257, 1:100), c-myc (Invitrogen, [9E10] MA1-980, 1:100), monocarboxylate transporter 4 (Santa Cruz Biotechnology, [D-1] sc-376140, 1:50) and vimentin (Cell Signaling Technology, [D21H3] 5741, 1:100 and Abnova, [SRL33] MAB9596, 1:200), were used. Secondary antibodies and counterstain used were donkey anti-rabbit Alexa Fluor 488 (Invitrogen, A21206, 1:500), donkey anti-mouse Alexa Fluor 488 (Invitrogen, A21202, 1:500), donkey anti-rabbit Alexa Fluor 568 (Invitrogen, A10042, 1:500), donkey anti-mouse Alexa Fluor 568 (Invitrogen, A10037, 1:500), and DAPI (4, 6-diamidino-2-phenylindole dihydrochloride; Invitrogen, D1306, 1:500).

Paraffin embedded tissues were cut into 4uM sections and placed on glass slides. The sections were deparaffinized with xylene and rehydrated. Sections were submerged into a Tris-EDTA solution (pH 9.0) or a citrate solution (pH 6.0) and heated for antigen retrieval. Sections were blocked and permeabilized with a solution of PBS + 0.2% triton + 2% bovine serum albumin for one hour at room temperature. Primary antibodies were diluted in PBS + 0.2% triton + 2% bovine serum albumin. The blocking solution was removed from slides and primary antibodies were applied to the sections overnight at 4°C. Sections were washed three times with PBS + 0.2% triton. All secondary antibodies and DAPI were diluted in PBS + 0.2% triton + 2% bovine serum albumin and added to the samples for one hour at room temperature. Sections were washed three times with PBS + 0.2% triton, and once a final time with reverse osmosis water. Samples were mounted in an aqueous mountant (Invitrogen, 00-4958-02). Each tissue section was co-stained for vimentin and one of the other markers of interest.

Whole-section multichannel images were captured using a Nikon Ti2-E motorized microscope with a Lumencor SOLA LED light engine (at 50% power), a Hamamatsu ORCA Fusion camera, and Nikon Plan Apochromat Lambda objectives using Nikon NIS-Elements AR version 5.30 software. Each sample was imaged at 10x magnification with a final 16-bit image resolution of 0.64 µm/pixel. Identical imaging settings were used for all samples within the same staining and imaging set.

Images were analyzed using NikoNIS-Elements AR software version 5.30 with the General Analysis 3 module. Semi-automated tumor segmentation was performed based on vimentin staining, and the segmented regions were manually modified by an unbiased operator as needed to ensure specific and complete tumor selection. Images were then rescaled to 1.28 µm/pixel and the intensities of all pixels representing tumor tissue were recorded in a frequency table for each sample. For assessing correlation between non-vimentin markers, image channels representing each marker of interest from different tissue sections were aligned using the multimodal image registration tool. Following alignment, an intensity profile line of 2-3 mm in length was plotted through tumor regions showing diverse signal landscapes, and the intensities of pixels along the profile line were recorded for each marker. All marker intensities were plotted by distance along the profile line in GraphPad Prism software version 9.0.0, and the curves were smoothed using a 2^nd^ order polynomial based on 50 neighboring points (25 on each side) to account for minor mismatches in image channels from different tissue sections. The Pearson correlation of each pair of markers was then calculated from the smoothed XY curves.

### Single-cell RNA-Seq

Tumors and lungs harvested from mice were processed using the human tumor dissociation kit (Miltenyi Biotec, 130-095-929) with a GentleMacs Octo Dissociator with Heaters (Miltenyi Biotec, 130-096-427). Single cell suspensions in 0.04% BSA-PBS of cell lines, dissociated tumor and lung tissues were generated and run on the Chromium Single Cell 3′RNA-sequencing system (10x Genomics) with the Reagent Kit v3.1 (10XGenomics, PN-1000121) according to the manufacturer’s instructions. Briefly, cells were loaded into Chromium Next GEM Chip G Single Cell Kit (10x Genomics, PN-1000120) with a targeted cell recovery of 5,000 cells per sample. After performing cDNA purification, amplification, and library construction as instructed, we sequenced sample libraries on a half lane of HS4000 (Illumina) to yield (after quality control) about 65,000 paired-end reads per cell. For samples that contained lineage tracing barcodes 0.9 uL (100uM) of Cellecta FSeqRNA-BC14 primer was added into the sample index pcr reaction.

### Cellular Barcoding

Barcoded lentivirus libraries were synthesized using CloneTracker XP™ 10M Barcode-3’ Library with Venus-Puro (plasmid) (Cellecta, BCXP10M3VP-P) as described in Wang and McManus 2009 ^39^ with polyethylenimine (PEI), (Alfa Aesar, 43896) as the transfection reagent. OS-17 cells were infected with in-house prepared virus library in the presence of Polybrene (8 μg/ml) (MilliporeSigma, TR1003G). The barcoded plasmids contain ∼10M 38-bp semi-random oligonucleotide sequence that is captured on the Chromium Single Cell 3′RNA-sequencing system. 48 hours post-infection, OS-17 cells were used to generate orthotopic primary and metastatic tumor models as described above.

### Cellular Barcoding Computational Analysis

Raw sequencing data was pre-processed to extract the lineage tag (LT) using known flanking sequences, and the matching cell ID (CID). Lines without high-confidence CID assignment were removed and redundant reads were removed. The extracted LT barcode reads were matched against the known Cellecta barcode library. Reads with barcodes that didn’t match the Cellecta library were eliminated. The LT barcode for each matching CID was integrated into the SEURAT object metadata to allow for further analysis (see below).

### Single-cell RNA-Seq analysis

Cell Ranger version 3.0.2 (10x Genomics) was used to convert Illumina BCL files to FASTQ files. These FASTQ files were then de-multiplexed and aligned to the hg19 human reference genome provided by 10X Genomics, to generate gene-cell matrices. We used the SEURAT R package^40–42^ for quality control, filtering, and analysis of the data. Cells were filtered to remove doublets (outliers with high count and high genes per cell), low quality cells (outliers with low count and low genes per cell), and cells with high mitochondrial genes (indicative of cells with broken membrane). Cells with fewer than 800 expressed genes and genes expressed in fewer than 5 cells were filtered out. The total numbers of cells in each model after filtering were as follows: Osteoblasts (cell culture): 5735; OS-17 (cell culture): 3,327; OS-17 (tibia): 4,574; OS-17 (lung): 2,849; 143B (cell culture): 3,354; 143B (tibia): 6,187; 143B (lung): 5,134; NCH-OS-2 (flank): 6,749; NCH-OS-2 (tibia): 4,285; NCH-OS-2 (lung): 1,645; NCH-OS-7 (flank): 1,998; NCH-OS-7 (tibia): 2,891; NCH-OS-7 (lung): 5,133. In analyses that required comparison across conditions or models, samples were subset to contain equal numbers of cells and this number is specified in the figure legends. We transformed and normalized unique molecular identifier (UMI) counts using the ‘NormalizeData’ function in SEURAT package version 4.0.2^43^ with default parameters. We mitigated the effects of cell cycle heterogeneity by regressing out canonical G2/M- and S-Phase genes using the ‘ScaleData’ function in SEURAT. Principle component analysis (PCA) was performed using the top 2,000 highly variable genes identified by the SEURAT function ‘FindVariableFeatures’ with default parameters. For each dataset, the first twenty principal components were selected based on the elbow plot for percentage explained variances, representing ∼55.5–60.3% of total variances. The UMAP transformation^44^ was performed on selected principal components using the ‘RunUMAP’ function. We applied shared nearest neighbor (SNN) modularity optimization^45^ for clustering. We calculated silhouette scores for different number of clusters (from two clusters to ten clusters) to measure how similar one cell was to other cells in its own cluster compared with other clusters using the silhouette function in the R package cluster. We determined the optimum number of clusters for each sample by maximizing the Silhouette score. Datasets were integrated using four different approaches (FastMNN^46^, SCTransform^47^, Harmony^48^, merge Seurat objects^49^) to determine the optimal method. PCA using the expression of cell cycle genes identified that integrating datasets with ‘merge’ function in SEURAT or application of Harmony gave consistent results where cells did not cluster by cell cycle. Marker genes were identified for each cluster relative to other clusters using the ‘FindMarkers’ function, returning only positive markers, with ‘test.use’ set to DESeq2^50^ which uses a negative binomial distribution. Pathway enrichment analysis was performed on these marker genes using the enricher function in the R package clusterProfiler^51^ with default parameters with msigdb Hallmark gene sets^52^. Pseudo-bulk analysis was performed by setting identities of each cell to the sample. We identified gene differentially regulated using the ‘FindMarkers’ function for both positive and negative markers upon tibia or lung colonization. Pathway enrichment analysis was performed on these markers as described above. The ITH score was calculated as described in Stewart et al.^18^ Briefly, it was defined as the average Euclidean distance between each cell to every other cell in each sample, in terms of the selected principal components.

We downloaded validation datasets from Gene Expression Omnibus (GEO) for seven patient primary tumor tissues published in a recent study (GEO accession number-GSE152048)^16^. We used the Seurat R package^40–42^ for quality control, filtering, and analysis of these gene expression matrices. Cells were filtered to remove doublets, low quality cells, and cells with high mitochondrial genes as described above. Cells with fewer than 800 expressed genes and genes expressed in fewer than 5 cells were filtered out. We transformed and normalized UMI counts using the ‘NormalizeData’ function in SEURAT package version 4.0.2^43^ with default parameters. Principle component analysis (PCA) was performed using the top 2,000 highly variable genes identified by the SEURAT function ‘FindVariableFeatures’ with default parameters. Using the marker gene sets used in the paper^16^, we separated tumor cells from patient host cells. For each sample, the first twenty principal components were selected based on the elbow plot for percentage explained variances, representing ∼51.8–61.5% of total variances. The UMAP transformation^44^ was performed on selected principal components using the ‘RunUMAP’ function. Module scores for Glycolysis, Hypoxia, EMT, TNFα signaling via NFκB were calculated using the ‘AddModuleScore’ function in SEURAT with corresponding Hallmark msigdb gene sets as input features for the expression program.

### Statistical analysis

Statistical analyses were performed using R software environment for statistical computing^53^ and Prism 9 (GraphPad Software, Inc.). The packages used in R software are mentioned through the text in the methods section. For differential gene expression analysis, a negative binomial model (DESeq2)^50^ was used. For multiple testing, p values were adjusted using Benjamini Hochberg (BH) correction. For ITH score comparisons, data were subjected to one-way analysis of variance (ANOVA) followed by Šidák multiple comparisons test.

## Abbreviations

PDX: patient-derived xenograft
ITH: intra-tumor heterogeneity
EMT: epithelial to mesenchymal transition
MTORC1: mechanistic target of rapamycin (mTOR) complex 1
PI3K: Phosphoinositide 3-kinase
AKT: Protein kinase B
TGF: Transforming growth-factor
TNFα: tumor necrosis factor α
NFKB: Nuclear Factor kappa-light-chain-enhancer of activated B cells

## Declarations

### Ethics approval and consent to participate

Patient-derived samples were obtained from patients consented under an Institutional Review Board (IRB)-approved protocol IRB11-00478 at Nationwide Children’s Hospital (Human Subject Assurance Number 00002860).

All animal studies were approved by Nationwide Children’s Hospital Institutional Animal Care and Use Committee (IACUC protocols AR15-00022 and AR14-00045).

### Consent for publication

Not applicable

### Availability of data and materials

The datasets supporting the conclusions of this article are available in the NCBI’s Gene Expression Omnibus (GEO) repository, [https://www.ncbi.nlm.nih.gov/geo/query/acc.cgi?acc=GSE179681].

### Code availability

The bioinformatics analyses were performed using open-source software, including SEURAT version 4.0.2^43^, clusterProfiler version 3.14.3^51^, and Cell Ranger version 3.0.2. All code is deposited in https://github.com/kidcancerlab/OSHetero2021.

## Acknowledgements

We would like to thank our funding sources. This work was generously supported by funding provided by NIH/NCI (K08CA201638, RDR), St Baldrick’s Foundation Scholar Award (RDR), Hyundai Hope on Wheels Young Investigator Award (RDR), CancerFree KIDS Foundation (RDR), Steps for Sarcoma Foundation (RDR), Sarcoma Foundation of America (RDR), a Pelotonia Fellowship (SR), a Nationwide Children’s Director’s Strategic Development Fund, and an NIH CTSA Grant UL1TR002733.

## Authors’ contributions

SR and RDR conceived and designed the study. SR and RDR developed the methodology. MC and ACG prepared libraries for next-generation sequencing. SR, EF, CT, MW, MVC and AO performed computation analysis. CAM performed immuno-fluorescence staining. TAV supported image acquisition and analysis of immunofluorescence experiments. SR and RDR interpreted all the consolidated data and wrote the manuscript. SR, EF, CAM and RDR edited the manuscript. SR, EF and RDR revised the manuscript. All authors read and approved the final manuscript.

## Additional Files

### Additional File 1

File format: .pdf

Title of the data: Supplemental Figures

Description of the data: Supplementary figures supporting the study can be found in this document.

### Additional File 2

File format: .docx

Title of the data: Supplementary Tables

Description of the data: Supplementary tables supporting the study can be found in this document.

