## Supplemental figures for "Osteosarcoma tumors maintain intra-tumoral transcriptional heterogeneity during bone and lung colonization"

Figure S1

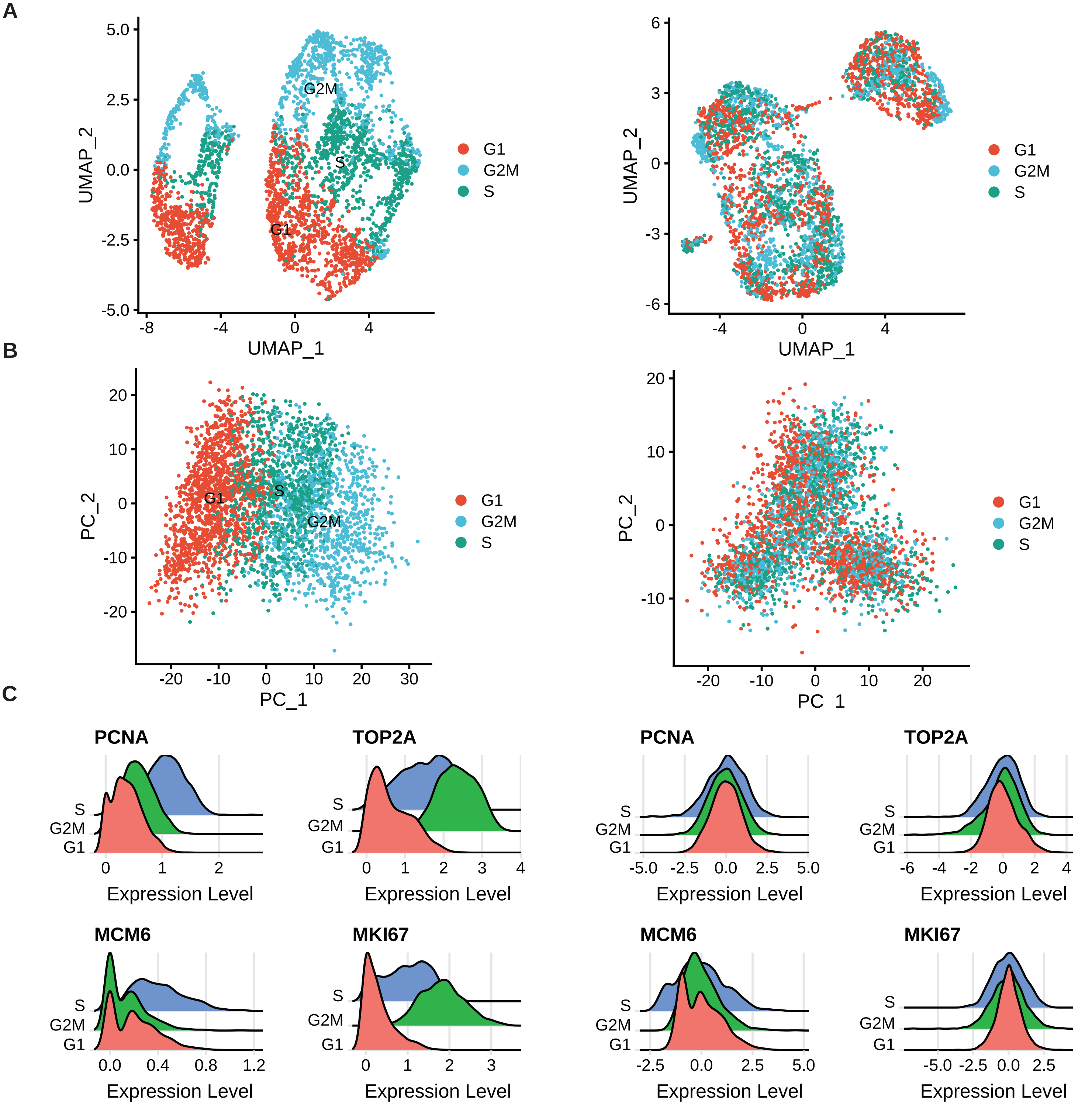


**Figure S1. Majority of the variance in our osteosarcoma datasets is dominated by cell cycle associated genes.** OS-17 cells were scored based on expression of G2/M and S phase markers, described in Tirosh et al. 2016[49]. A) UMAP analysis and B) PCA on cell cycle genes before cell cycle regression reveals that cells separate entirely by phase, confirmed using C) Ridge plots visualizing the distribution of cell cycle markers. After regressing out cell cycle scores, UMAP analysis and PCA on cell cycle genes show that cells no longer separate by phase. This is confirmed using Ridge plots of cell cycle markers. These plots are representative of all datasets.

### Figure S2

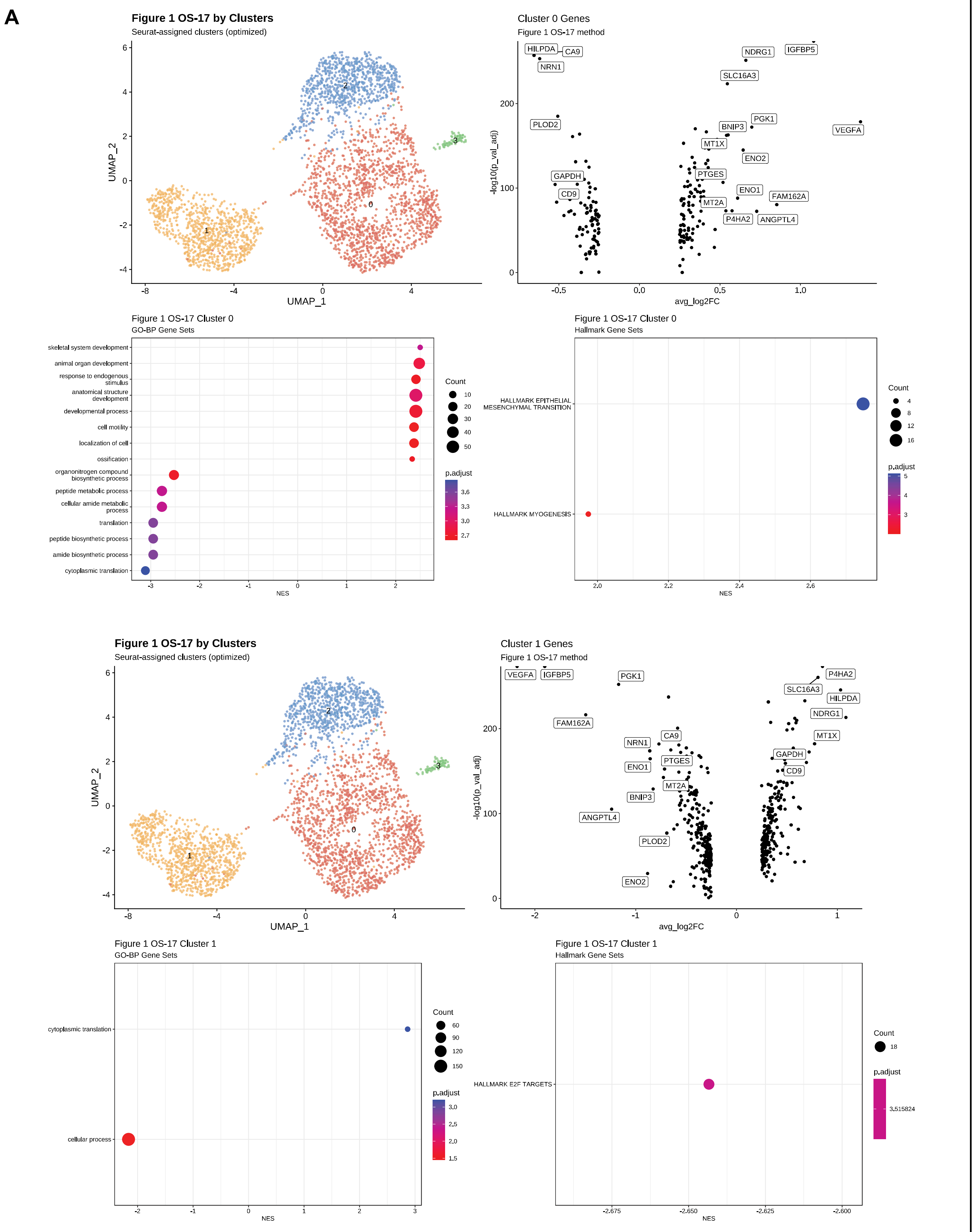

A
Figure 1 OS-17 by Clusters
Seurat-assigned clusters (optimized)
UMAP_1
UMAP_2
Cluster 0 Genes
Figure 1 OS-17 method
avg_log2FC
-log10(p_val_adj)
HILPDA
CA9
NRN1
NDRG1
IGFBP5
SLC16A3
PLOD2
PGK1
BNIP3
VEGFA
MT1X
ENO2
GAPDH
PTGES
CD9
ENO1
FAM162A
MT2A
P4HA2
ANGPTL4
Figure 1 OS-17 Cluster 0
GO-BP Gene Sets
skeletal system development
animal organ development
response to endogenous stimulus
anatomical structure development
developmental process
cell motility
localization of cell
ossification
organonitrogen compound biosynthetic process
peptide metabolic process
cellular amide metabolic process
translation
peptide biosynthetic process
amide biosynthetic process
cytoplasmic translation
NES
Count
p.adjust
Figure 1 OS-17 Cluster 0
Hallmark Gene Sets
HALLMARK EPITHELIAL MESENCHYMAL TRANSITION
HALLMARK MYOGENESIS
Figure 1 OS-17 by Clusters
Seurat-assigned clusters (optimized)
Cluster 1 Genes
Figure 1 OS-17 method
Figure 1 OS-17 Cluster 1
GO-BP Gene Sets
cytoplasmic translation
cellular process
Figure 1 OS-17 Cluster 1
Hallmark Gene Sets
HALLMARK E2F TARGETS
3,515824

### Figure S2 continued

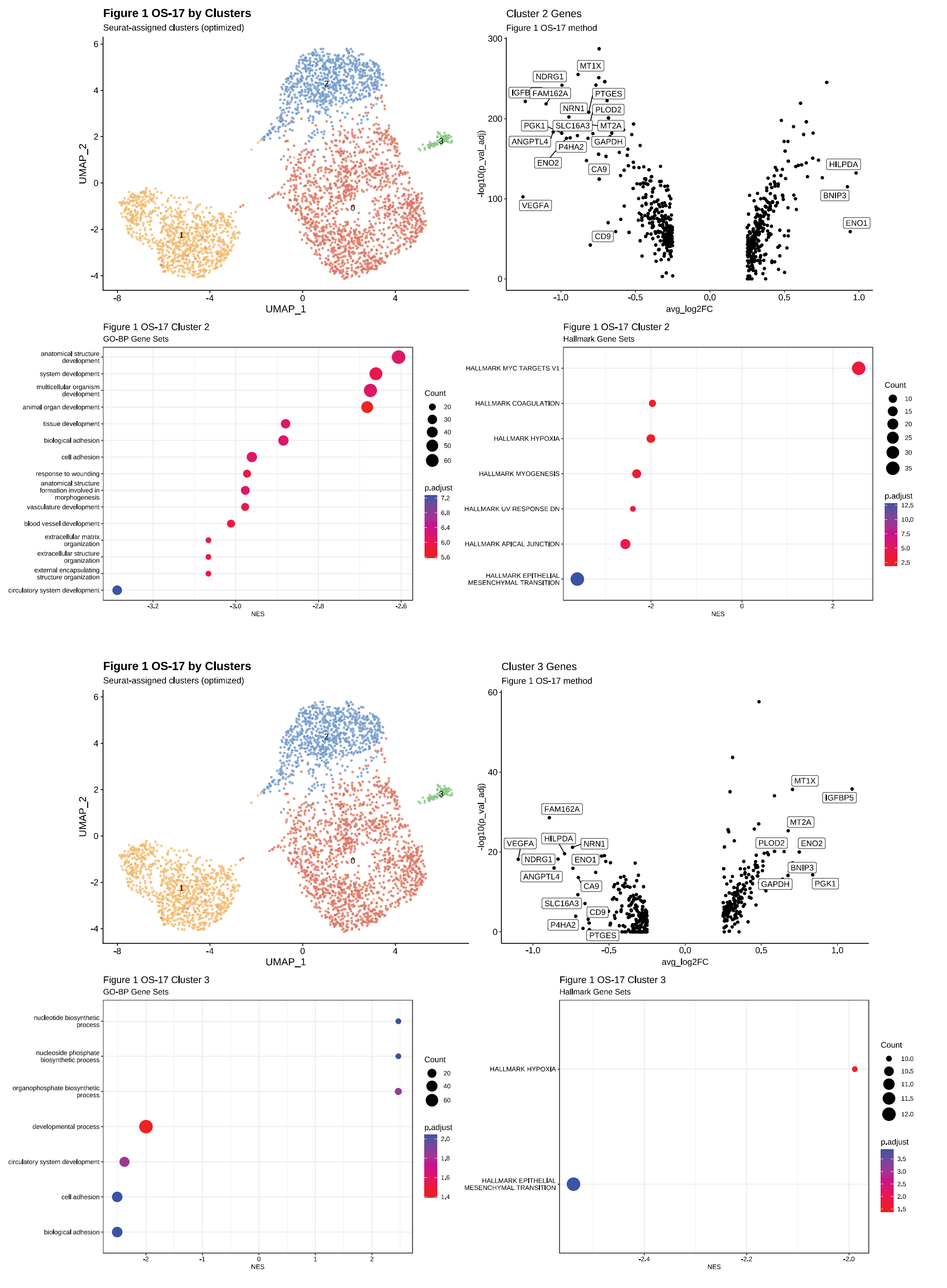

Figure S2 continued

B

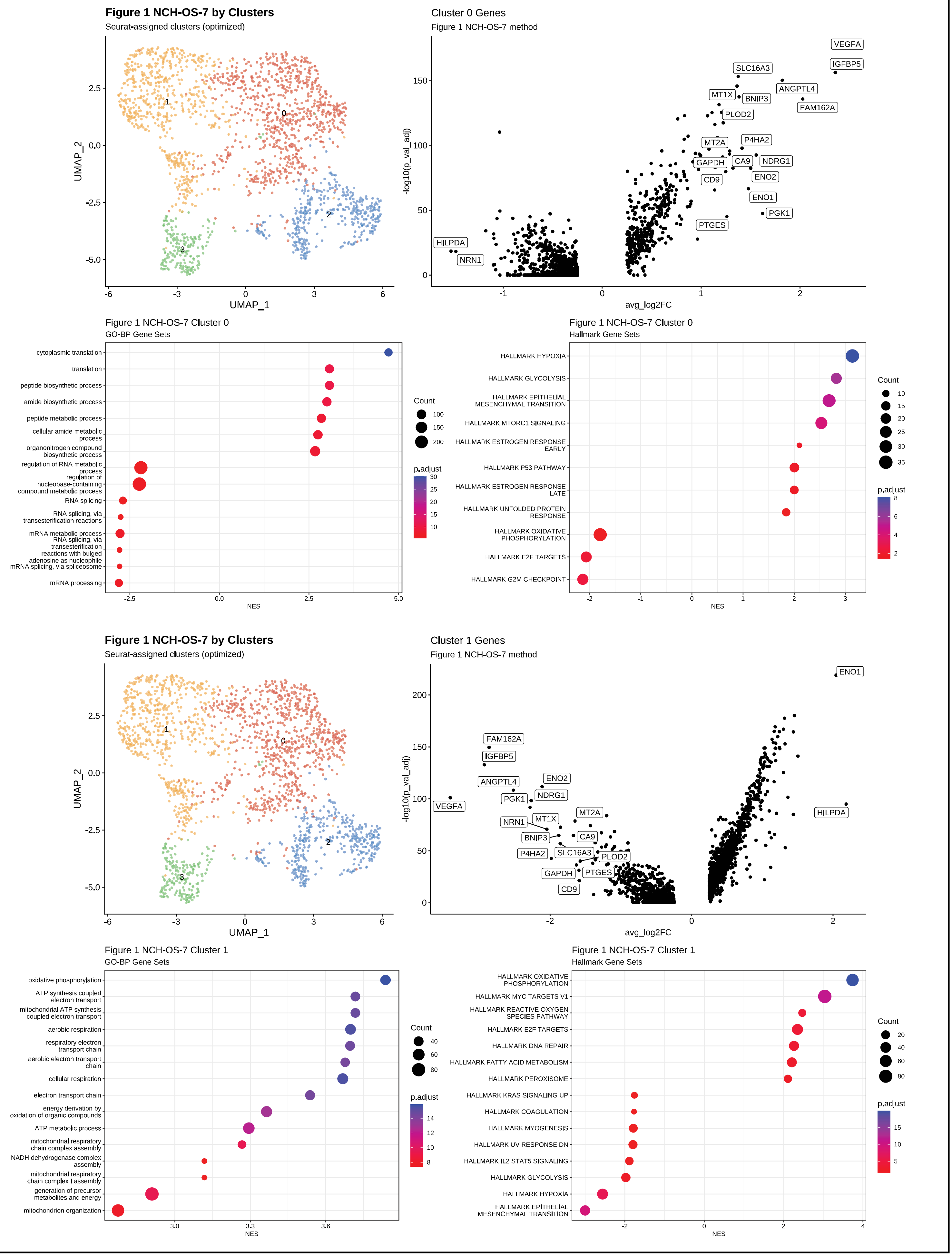

### Figure S2 continued

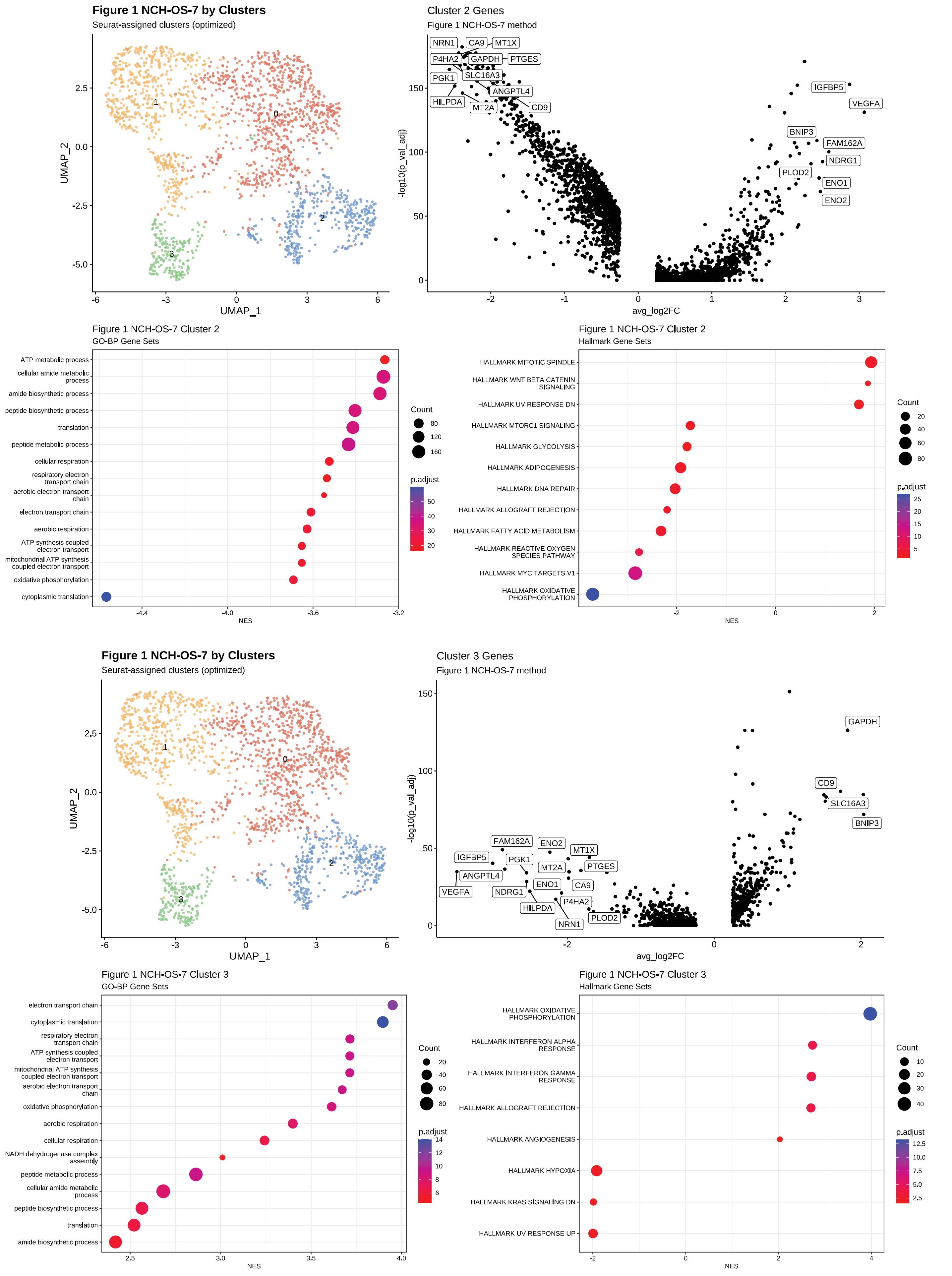

### Figure S2 continued

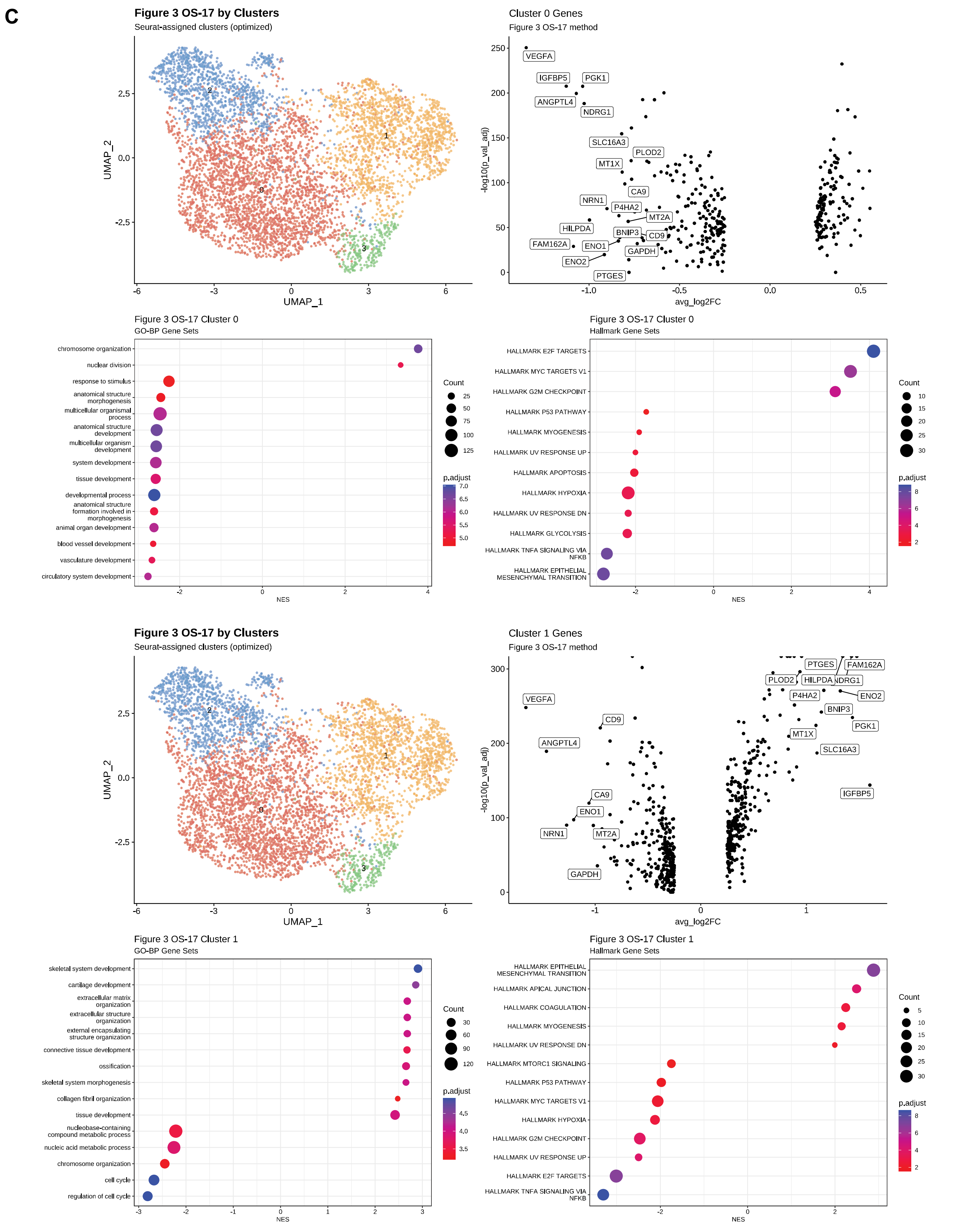

### Figure S2 continued

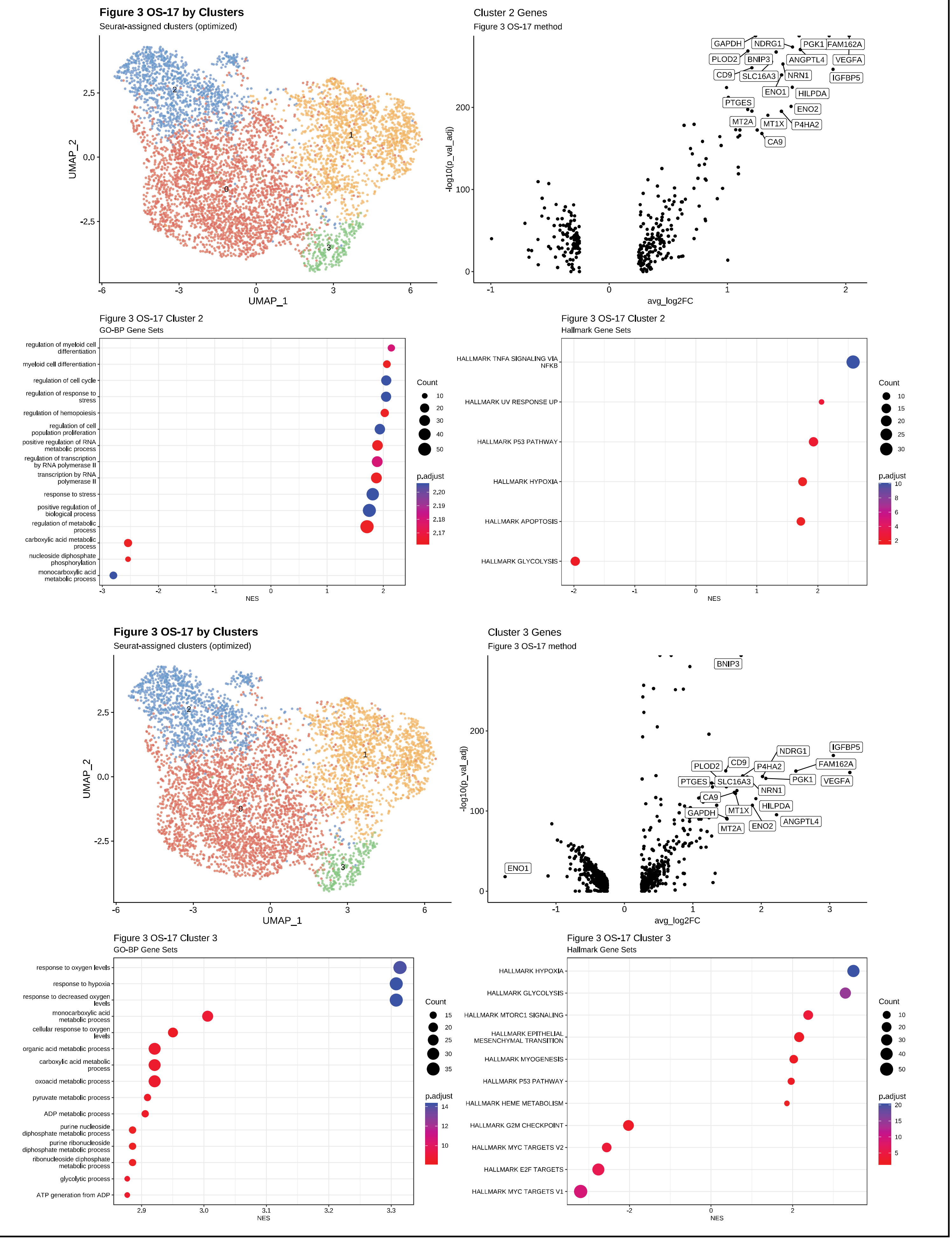

Figure S2 continued

D

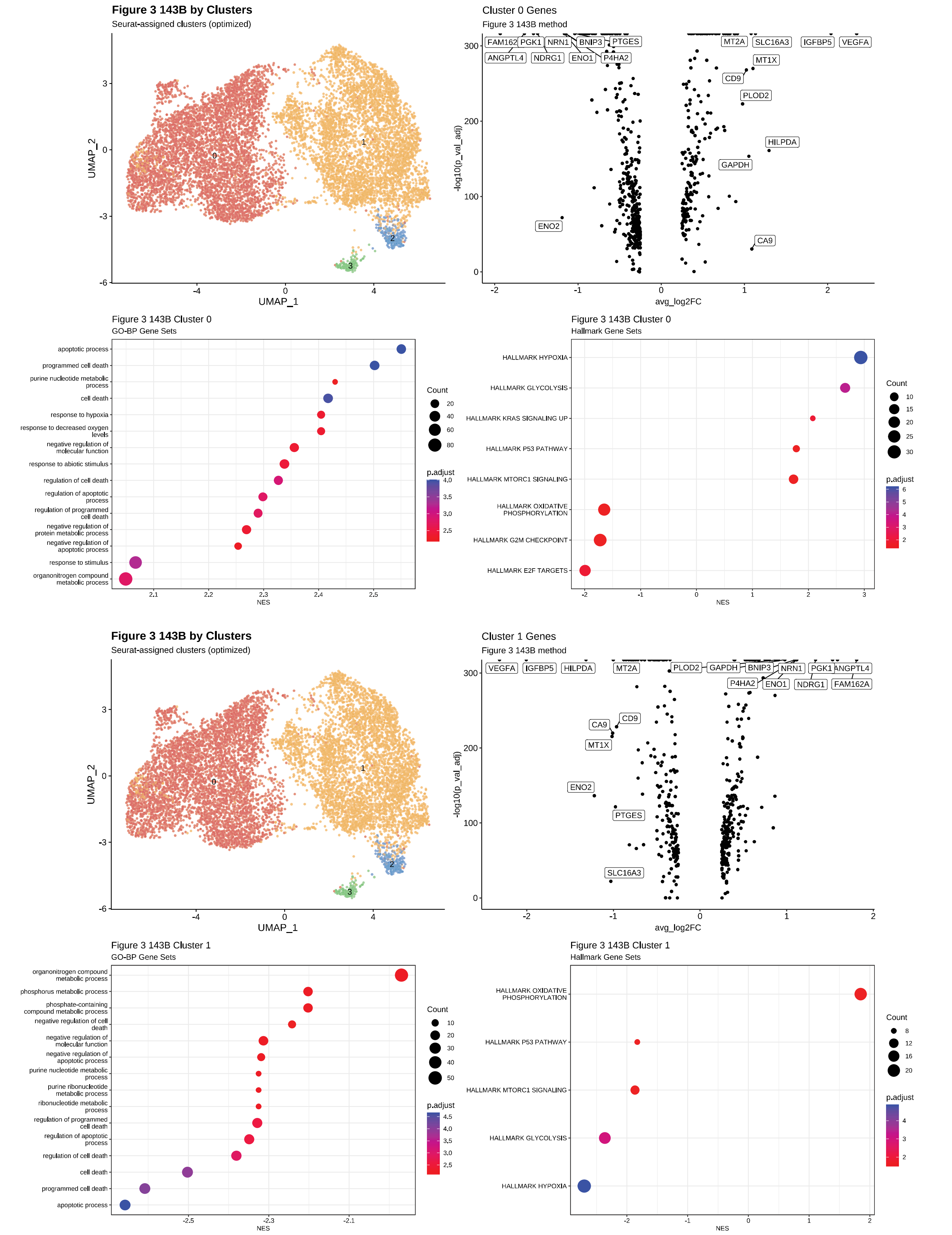

### Figure S2 continued

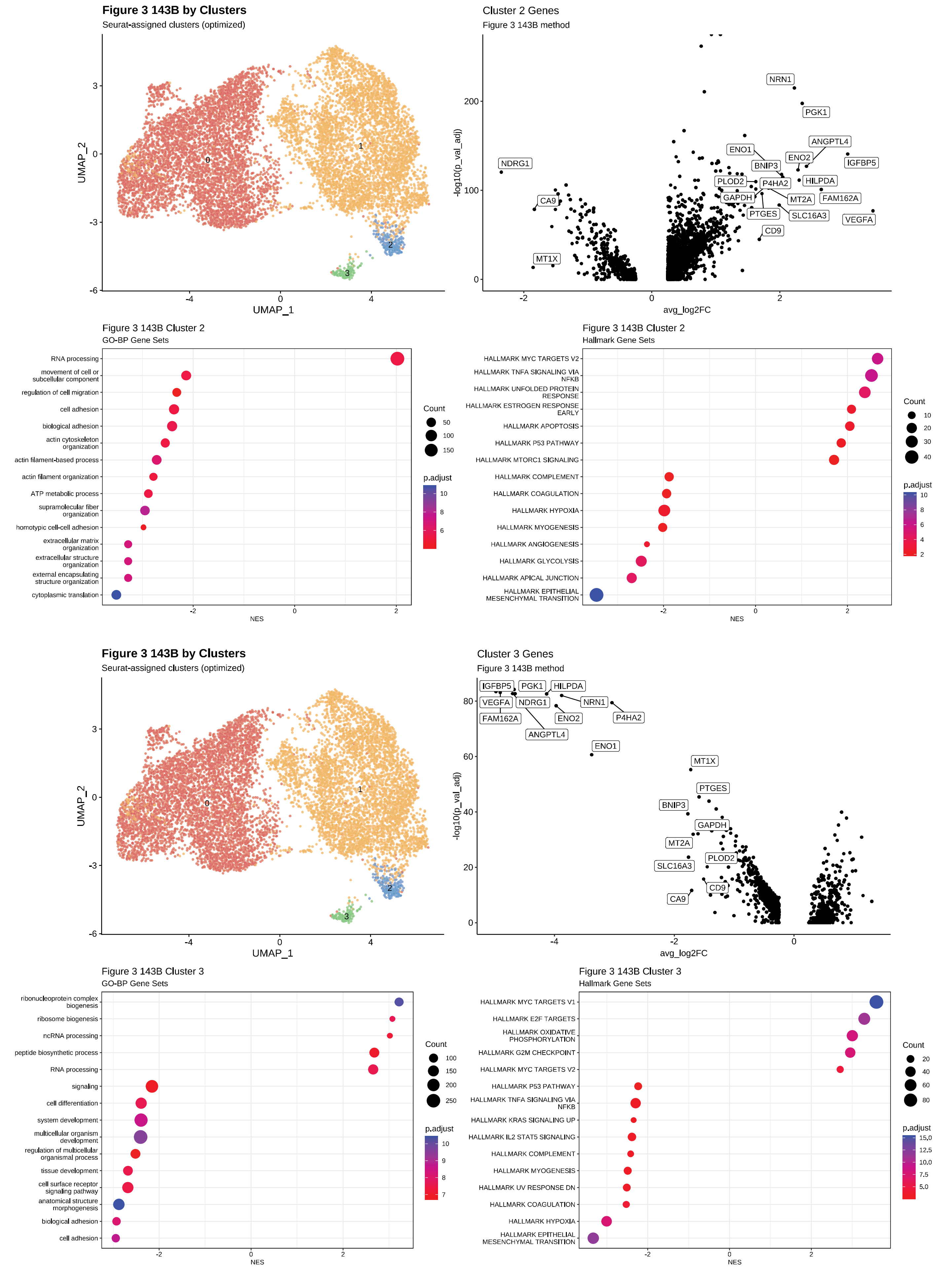

### Figure S2 continued

**E**

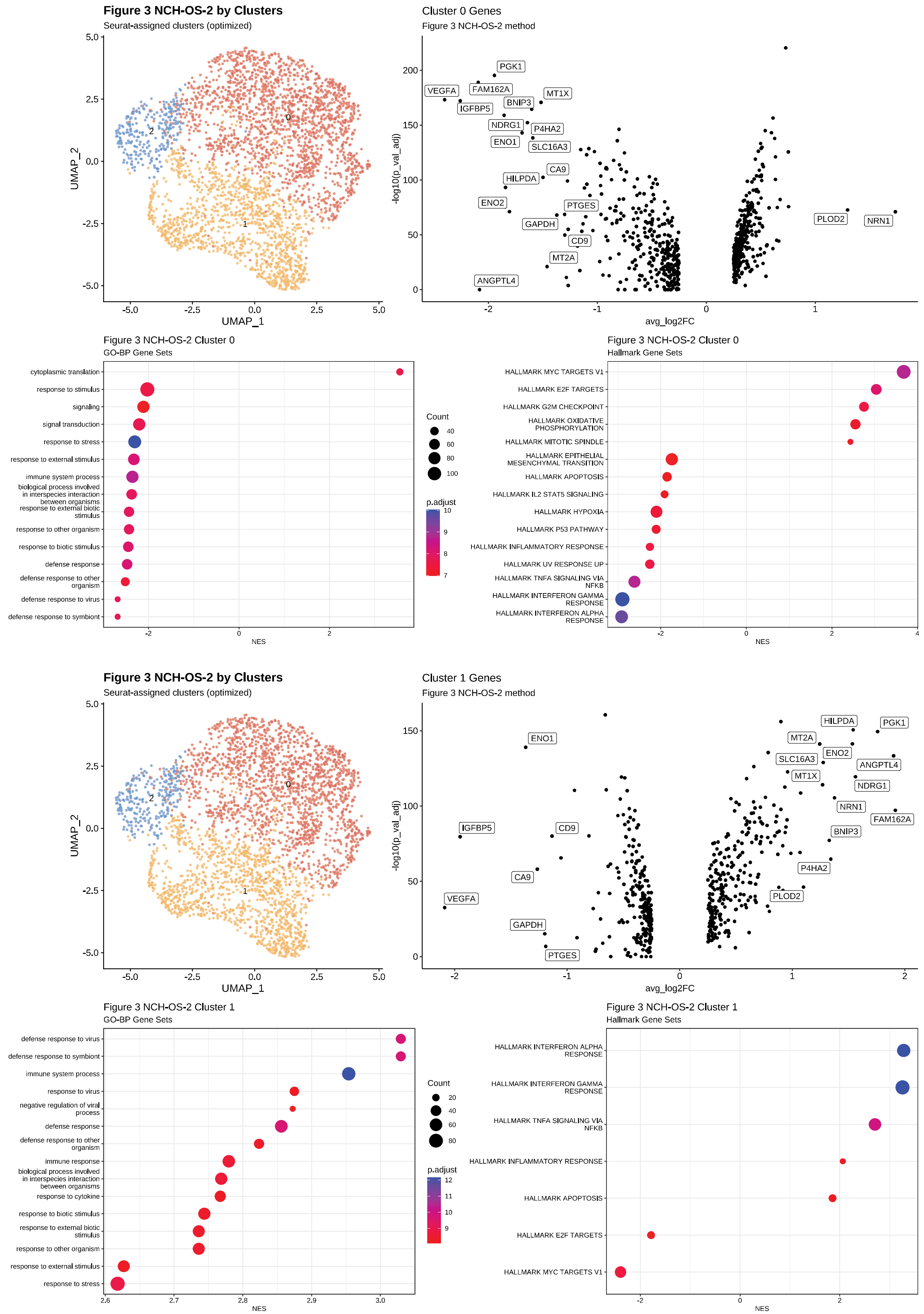

### Figure S2 continued

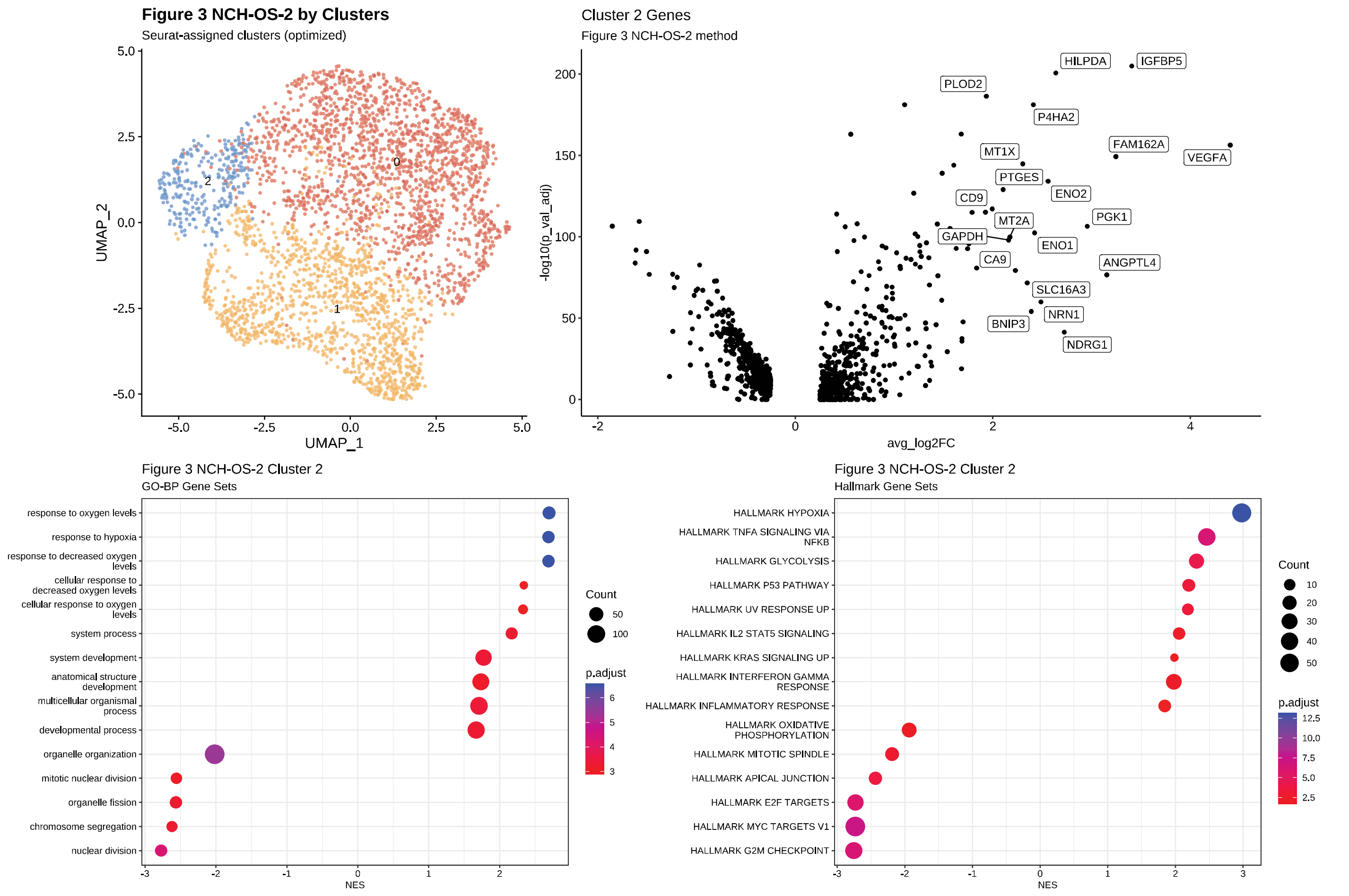


**F**

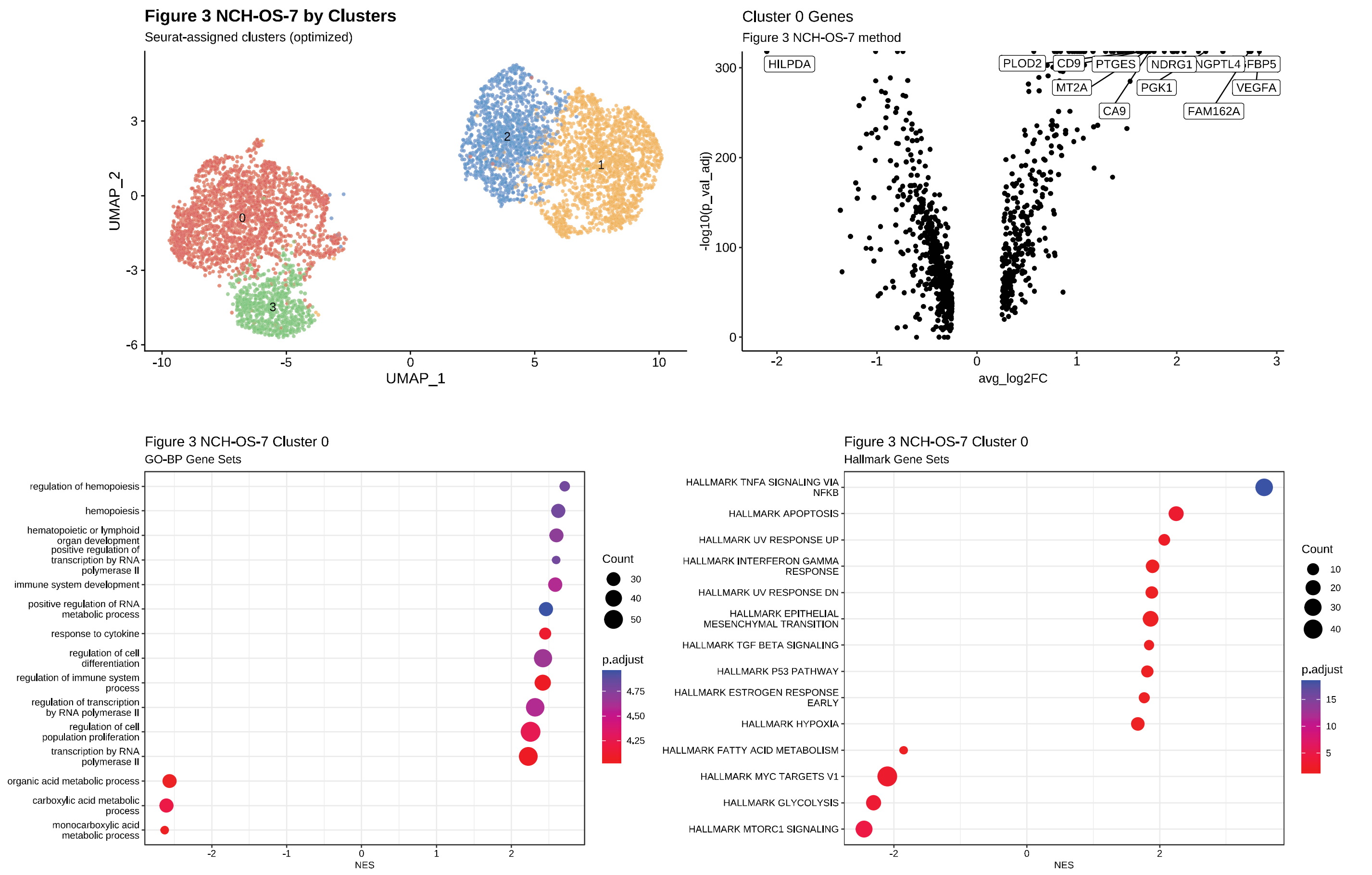

### Figure S2 continued

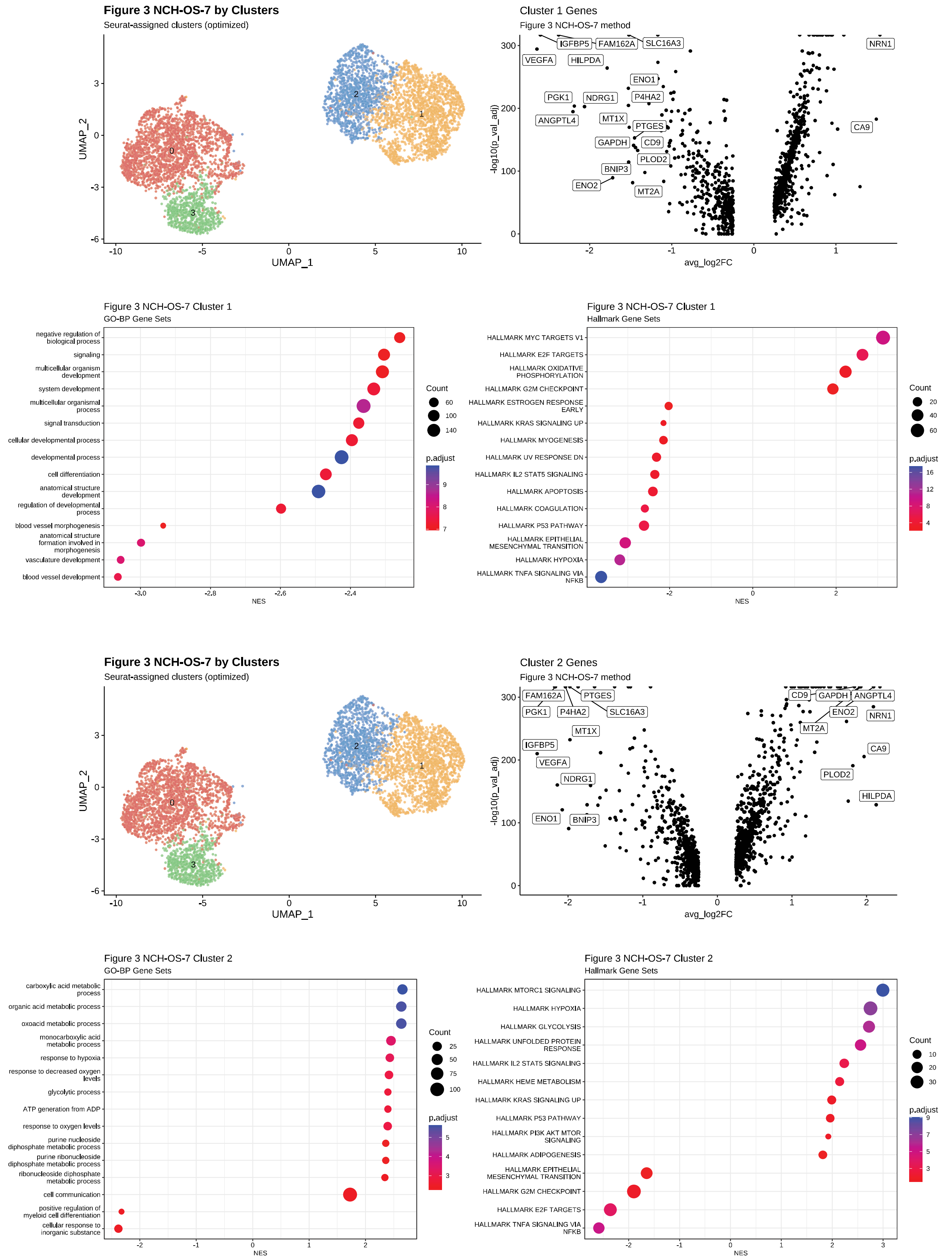

### Figure S2 continued

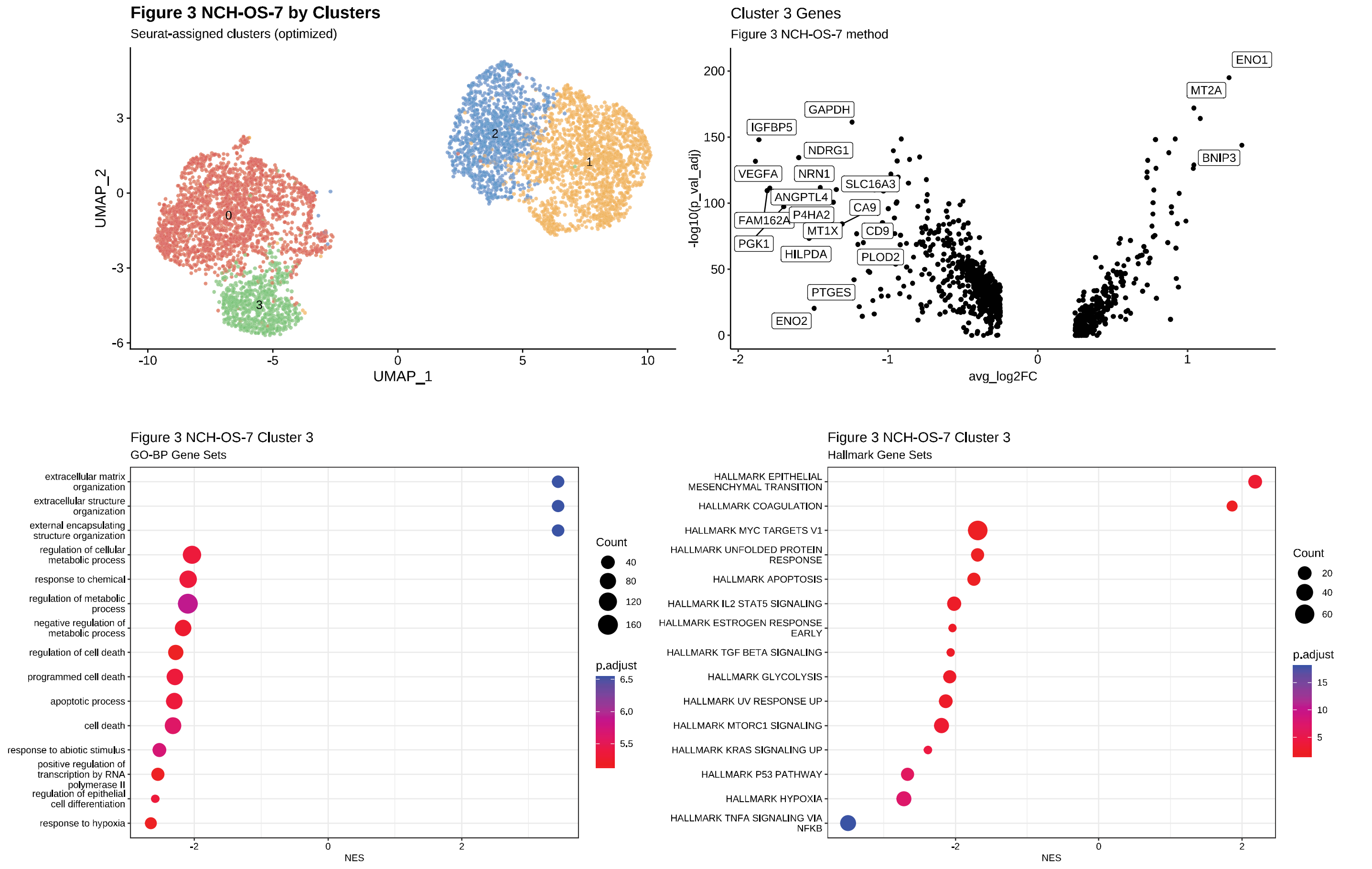


**G**

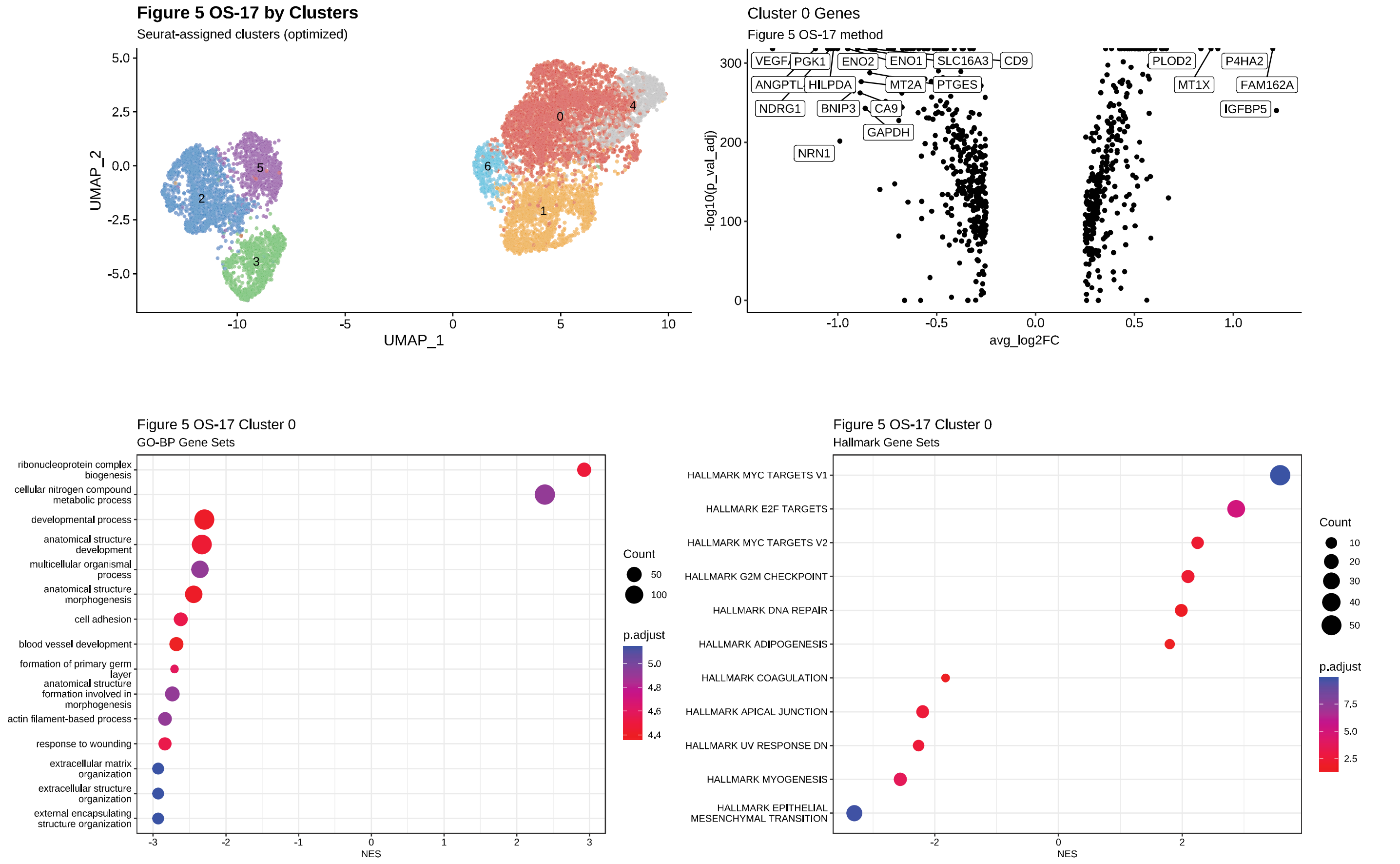

### Figure S2 continued

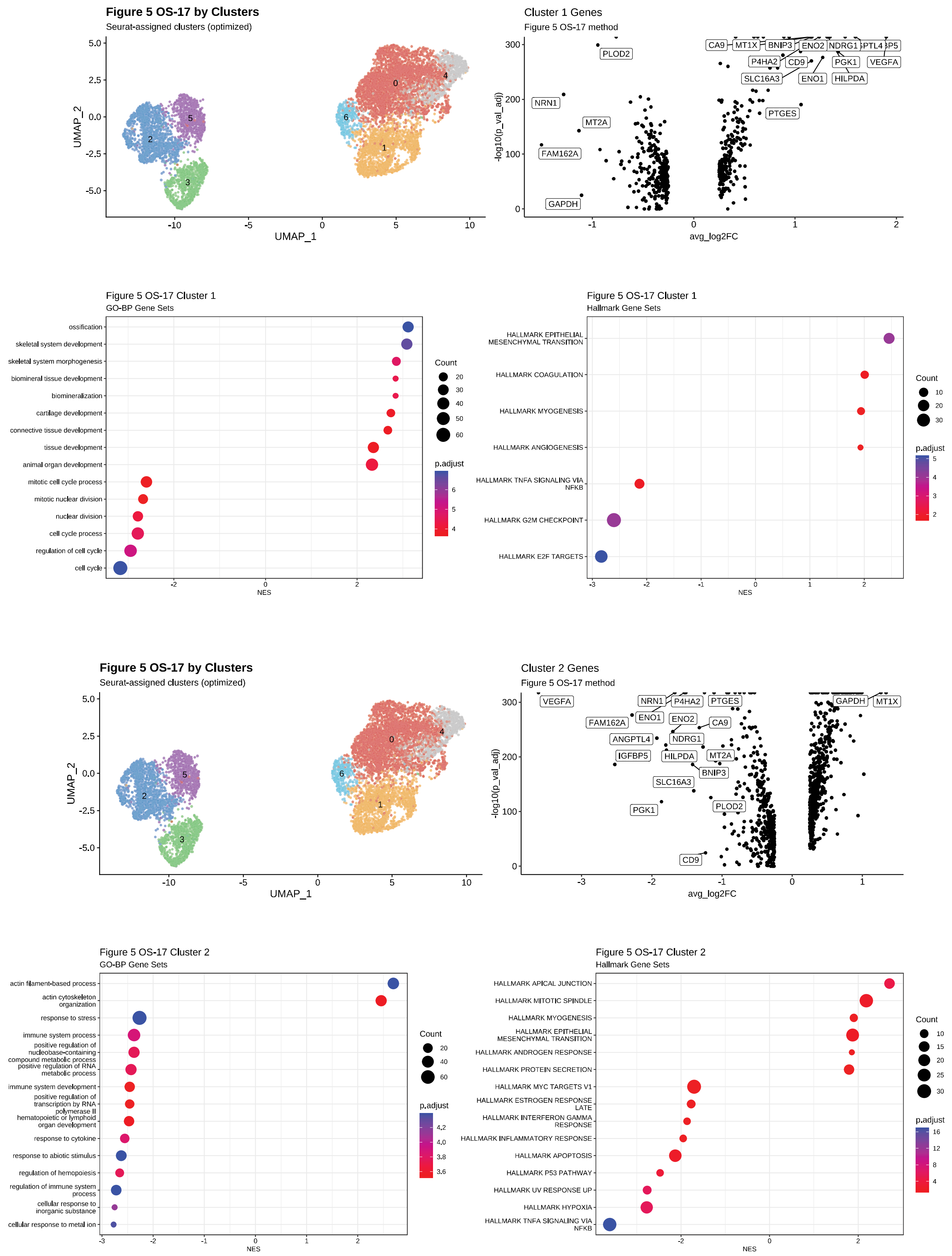

### Figure S2 continued

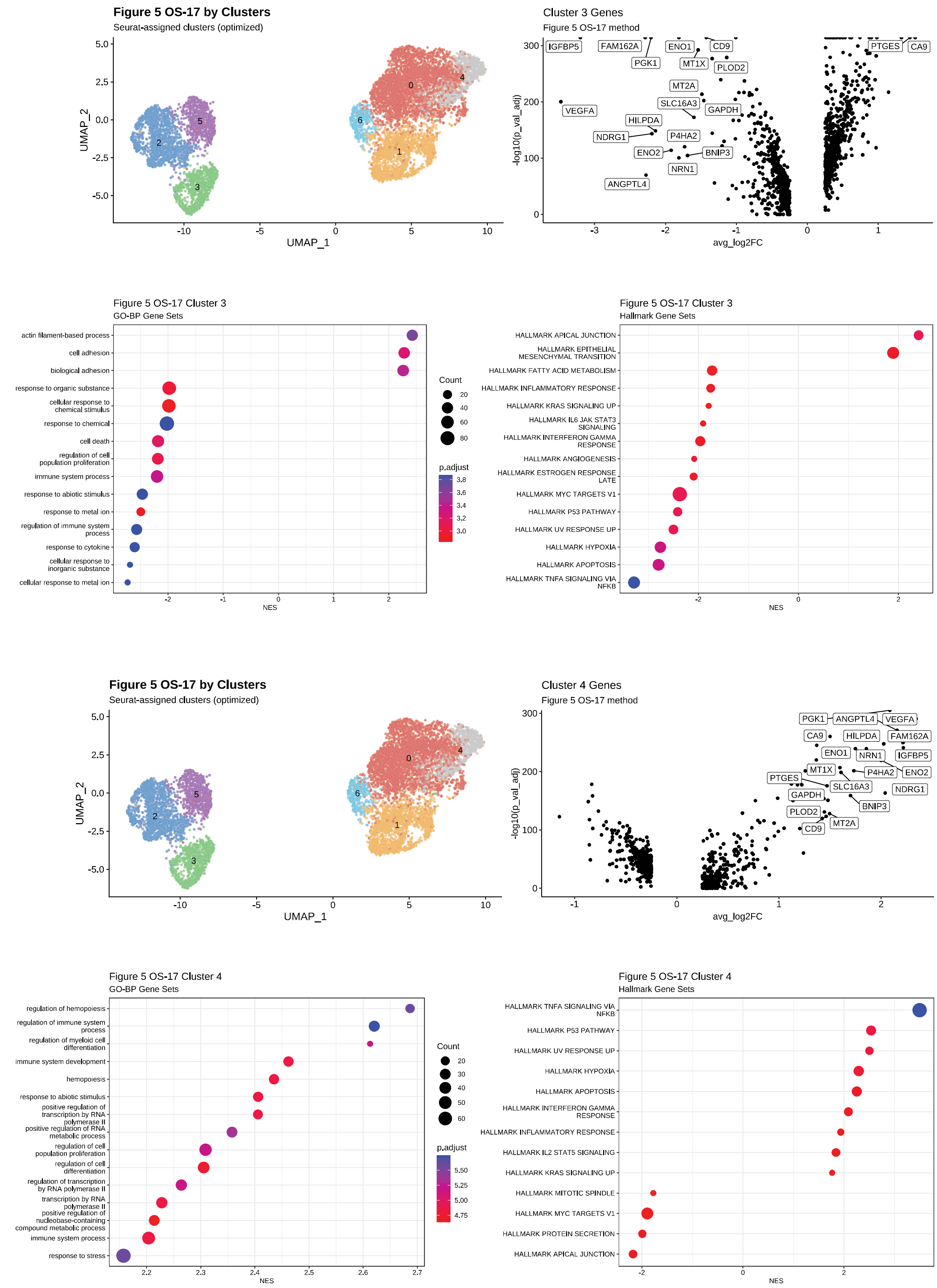

### Figure S2 continued

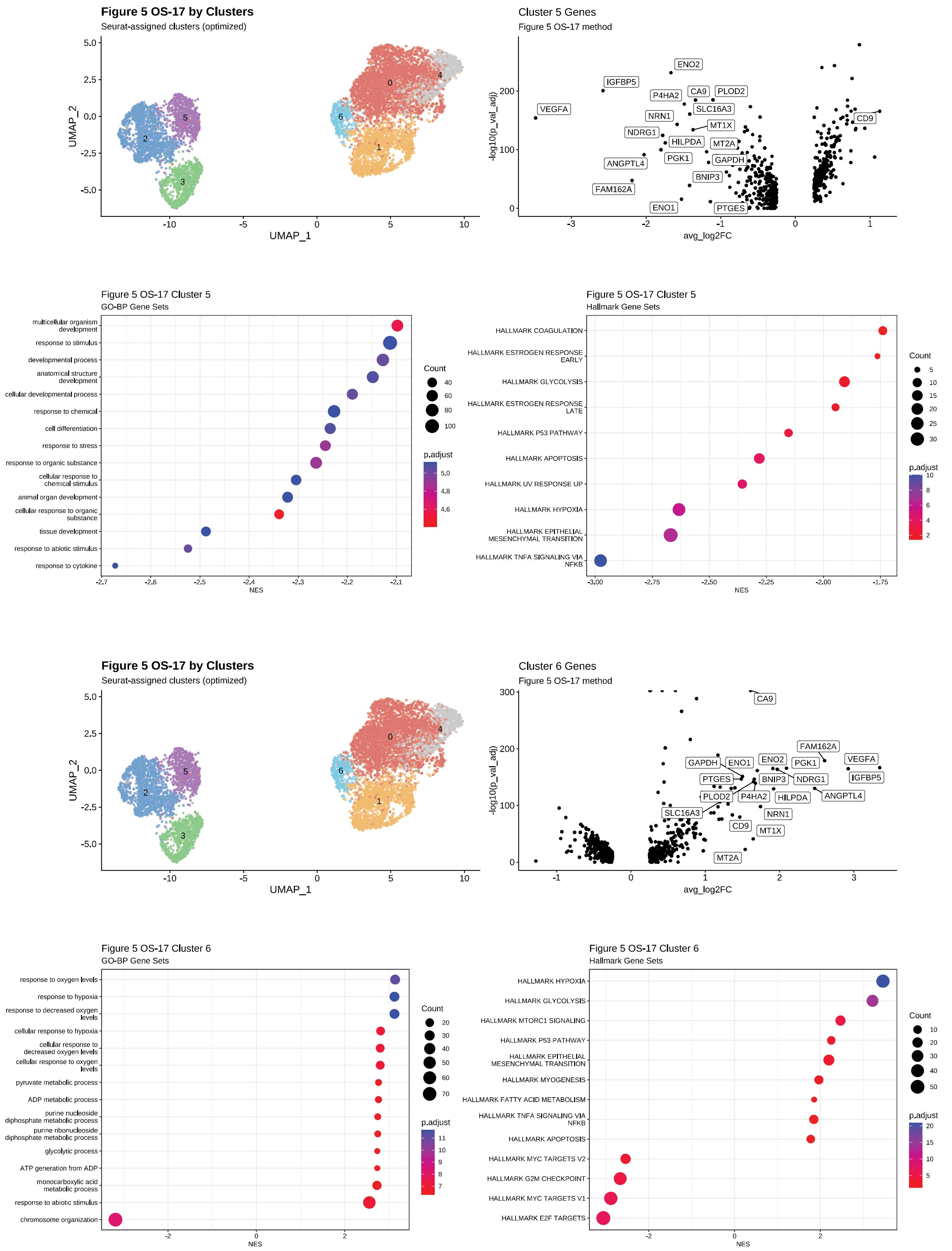

Figure S2 continued

**Figure S2. In-depth analysis of clusters utilized in analyses.** A) OS-17 cell culture and B) NCH-OS-2 flank cluster analysis (Figure 1). C-F) Merged tibia and lung cluster analysis for OS-17, 143B, NCH-OS-2, and NCH-OS-7, respectively (Figure 3). G) Lineage-tagged OS-17 cell culture, tibia, and lung cluster analysis (Figure 5). The upper left of each analysis displays all clusters for the dataset. The upper right shows a volcano plot analysis, where the top differentially expressed genes for the cluster of interest, compared to all others in the dataset, are shown. DotPlots on the bottom left depict the significantly upregulated GO BioProcesses and GO MolFunction gene lists in the cluster of interest from the differentially expressed gene list for the cluster. Bar plot on the bottom left shows upregulated msigdb Hallmark pathways from the differentially expressed gene list for the cluster. The Wilcoxon rank sum test, or “wilcox” statistical method, was used to generate differentially expressed gene lists.

### Figure S3

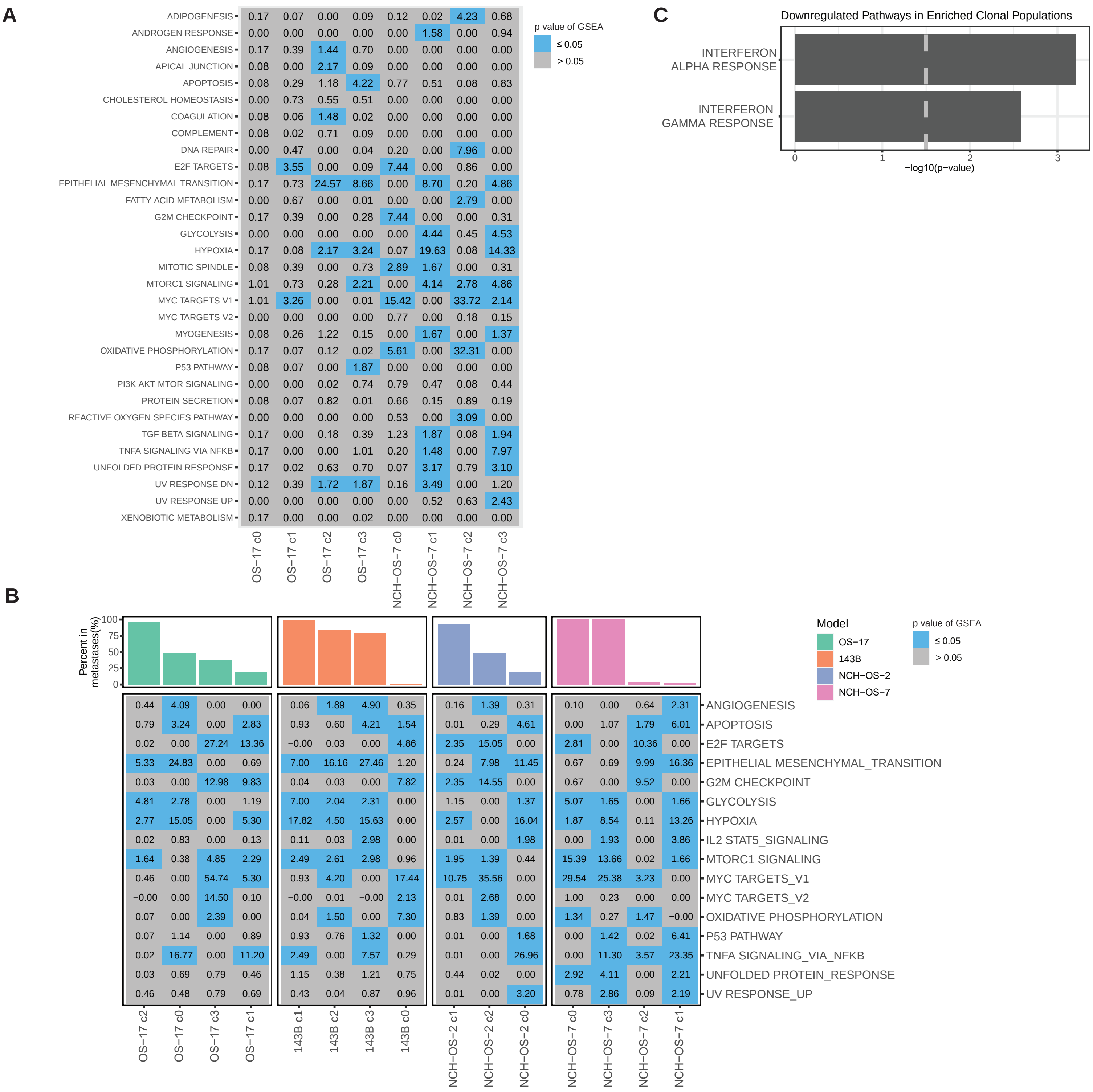


**Figure S3. Downregulated pathways across all cell line- and PDX-derived tumors.** A) Pathway enrichment analysis for hallmark gene sets associated with genes downregulated in distinct clusters identified in OS-17 and NCH-OS-7 osteosarcoma models (Figure 1C). B) Heterogeneity in downregulated hallmark pathway gene sets. Bar plots show percentage of cells identified per cluster in lung lesions. C) Pathway enrichment analysis for hallmark gene sets comparing enriched clones relative to the remaining lung-colonizing tumor cells identifies genes related to interferon alpha response and interferon gamma response to be significantly downregulated. For A-B, we used a pathway enrichment analysis for hallmark gene sets using genes differentially downregulated in each cluster relative to every other cluster within the same model. P values were adjusted for multiple comparisons. Boxes in grey identify non-significant pathway enrichments, whereas boxes in blue identify statistically significant enrichments. For C, only statistically significant (p<0.05) downregulated pathways in enriched clones relative to the remaining lung-colonizing tumor cells are shown.

Figure S4

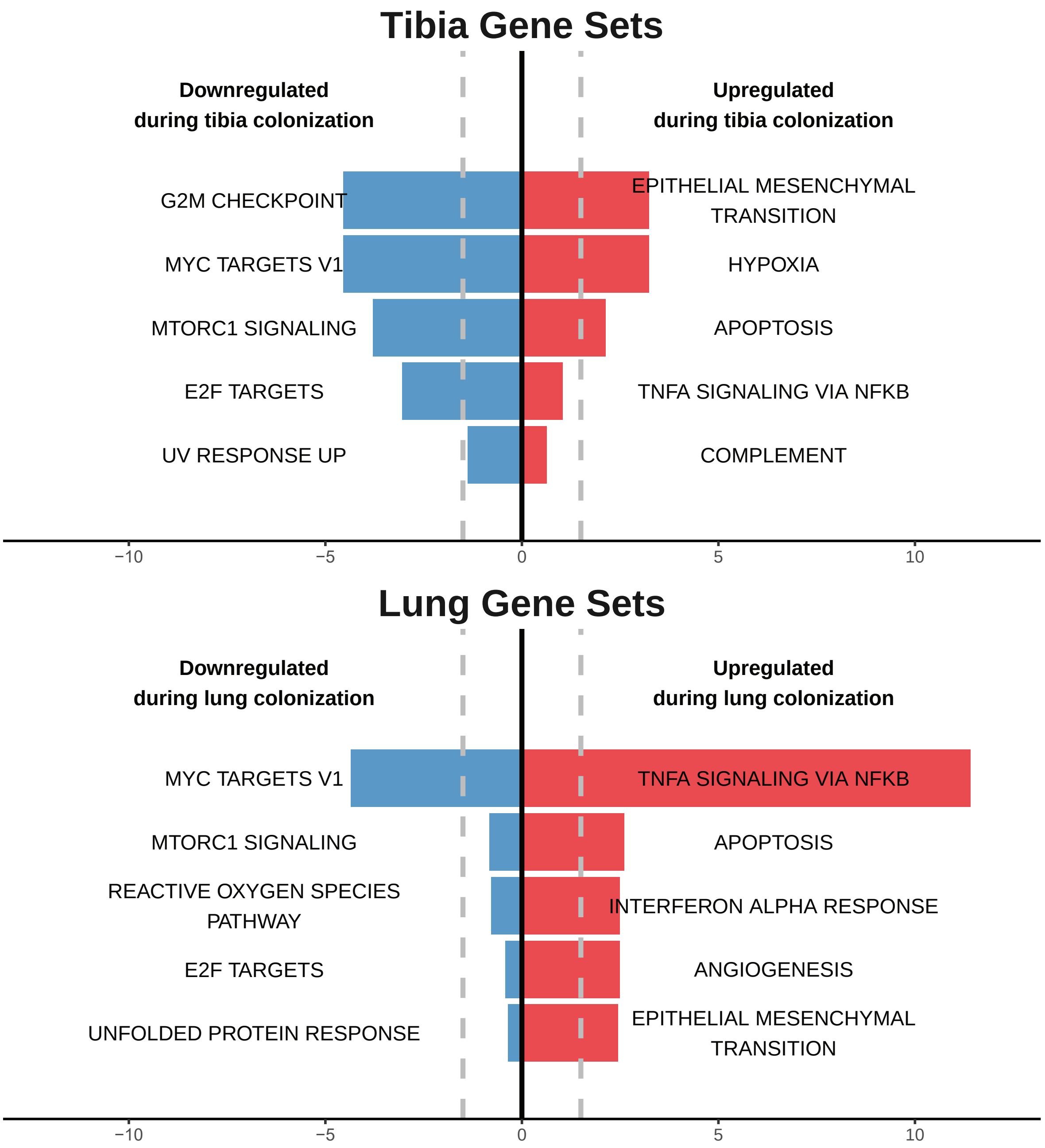


**Figure S4. Mouse osteosarcoma models share distinct transcriptional profiles with human osteosarcoma models during tibia and lung colonization.** K7M2 and F420 mouse models of osteosarcoma were added to the pathway enrichment analysis with the human models (OS-17, 143B, NCH-OS-2, NCH-OS-7) to determine conserved enriched pathways between mouse and human models. Pathway enrichment analysis with adjusted p values for hallmark gene sets associated with shared genes from the overlap of differentially expressed genes that are up- or down-regulated in tibia or lung lesions relative to corresponding starting population of cells (cell line or PDX flank tumors).

Figure S5

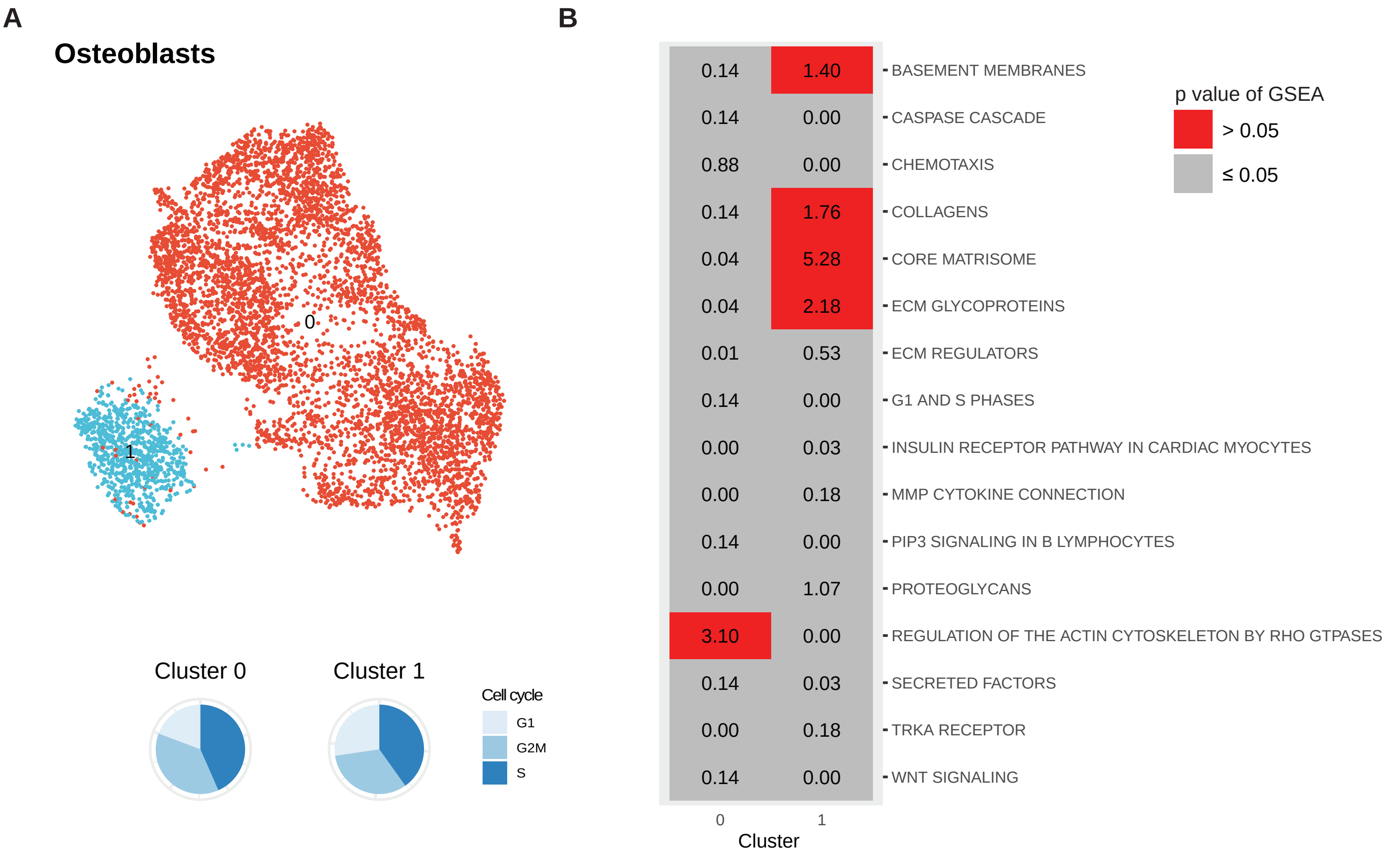


**Figure S5. Osteoblast cells demonstrate phenotypic heterogeneity.** A) UMAP analysis of normal human osteoblast cells in cell culture. B) Pathway enrichment analysis against msigdb C2 canonical pathways on genes differentially upregulated in each cluster relative remaining cells in the dataset.

Figure S6

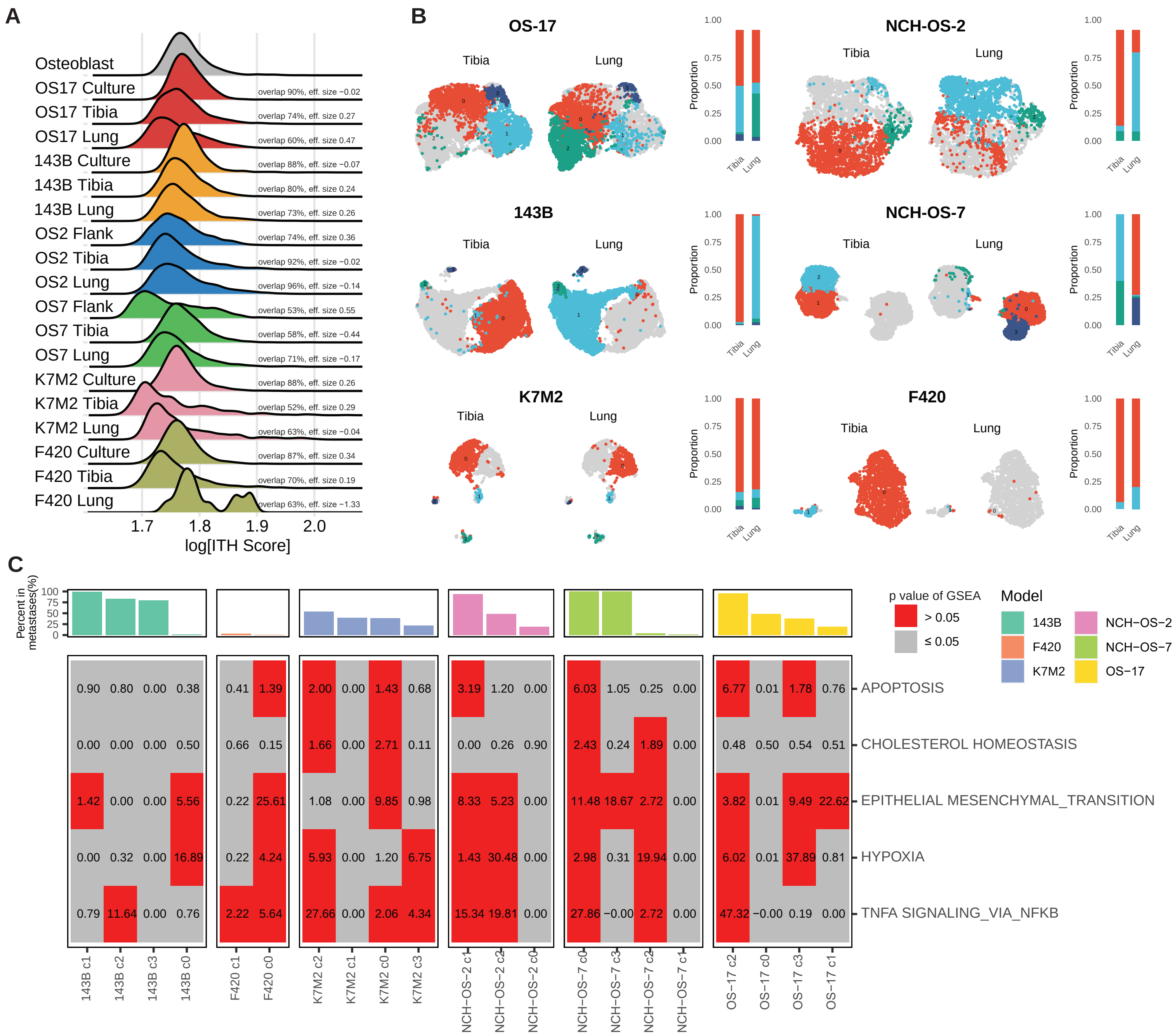


**Figure S6. Additional mouse models of osteosarcoma retain phenotypic heterogeneity despite adaptive changes in response to changing microenvironments.** A) Mouse osteosarcoma models maintained overall heterogeneity with a high degree of overlap between conditions. The ridge plot shows ITH scores, which represent the gene expression “distance” between each tumor cell within a sample and all of the other tumor cells from that same sample. The overlap statistic describes the total percentage of overlap in the observed distributions between two samples. B) UMAP analysis for merged tibia- and lung-colonizing tumor samples in each of the six models, including the mouse osteosarcoma models. Cells in grey represent remaining cells in merged sample. Cluster enrichment analysis shows distribution of cells in each cluster in the two microenvironment conditions (tibia, lung). While some cells in the tibia and lung lesions adopted shared phenotypes, others adopted distinct phenotypes. C) Glycolysis activation is not shared in mouse osteosarcoma models, but hypoxia heterogeneity is maintained. We used a pathway enrichment analysis for hallmark gene sets using genes differentially upregulated in each cluster relative to every other cluster within the same model. P values were adjusted for multiple comparisons. Boxes in grey identify non-significant pathway enrichments, whereas boxes in red identify statistically significant enrichments. Bar plot shows percentage of cells identified per cluster in lung lesions.

Figure S7

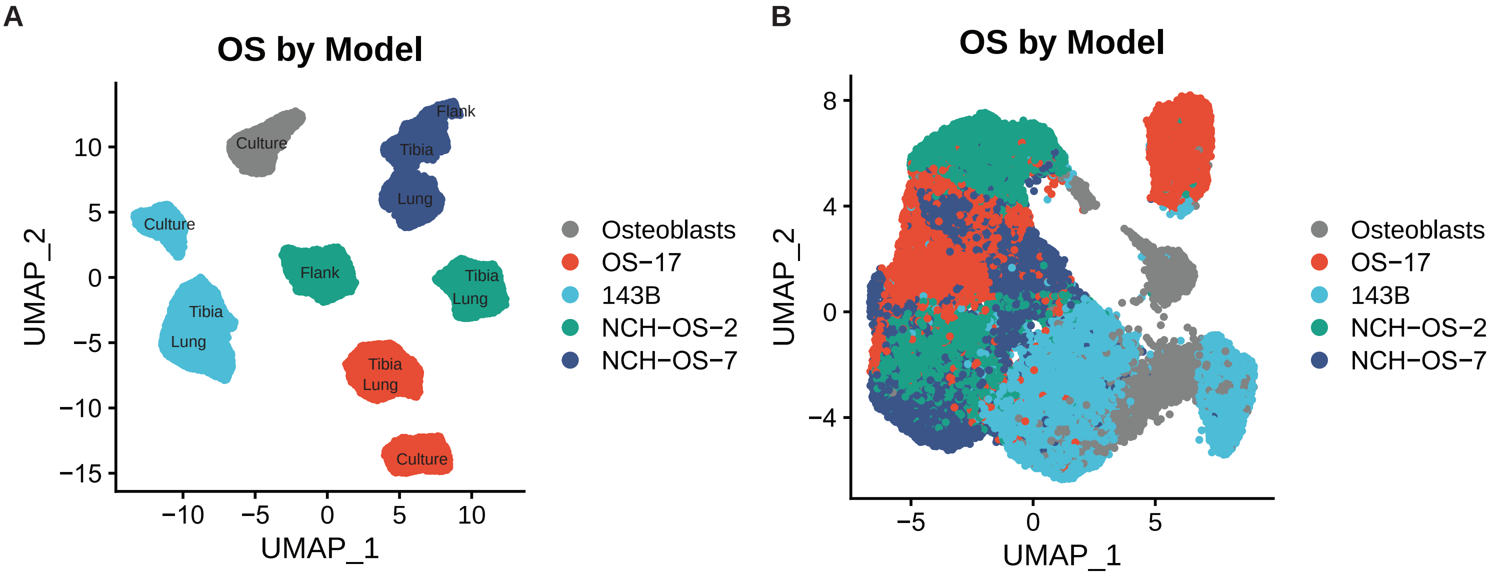


**Figure S7. High inter-tumor heterogeneity identified between models.** A) UMAP analysis of merged datasets. B) Application of Harmony forces similarity between heterogeneous populations of cells.

Figure S8

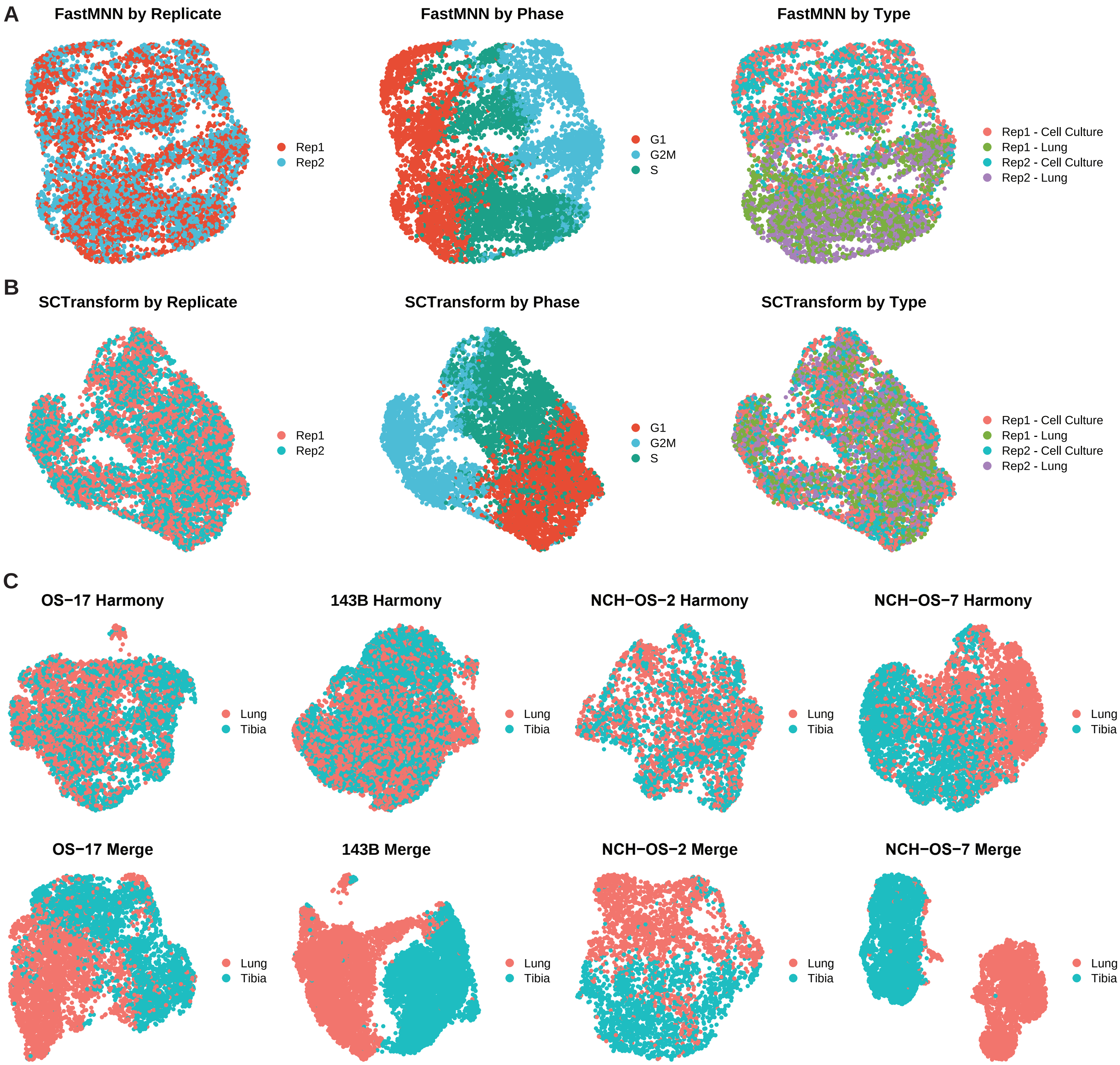


**Figure S8. Integrative analysis for within-model comparison of primary and metastatic lesions.** A, B) Application of FastMNN or SCTransform between replicate samples demonstrated complete overlay, however cells separated largely by state of cell cycle (OS-17). C) Application of Harmony or merging to Seurat objects. Application of Harmony demonstrated nearly complete overlay. Merged objects displayed some overlay while maintaining a slight distinction between samples. Cells did not cluster based on state of cell cycle (data not shown).

Figure S9

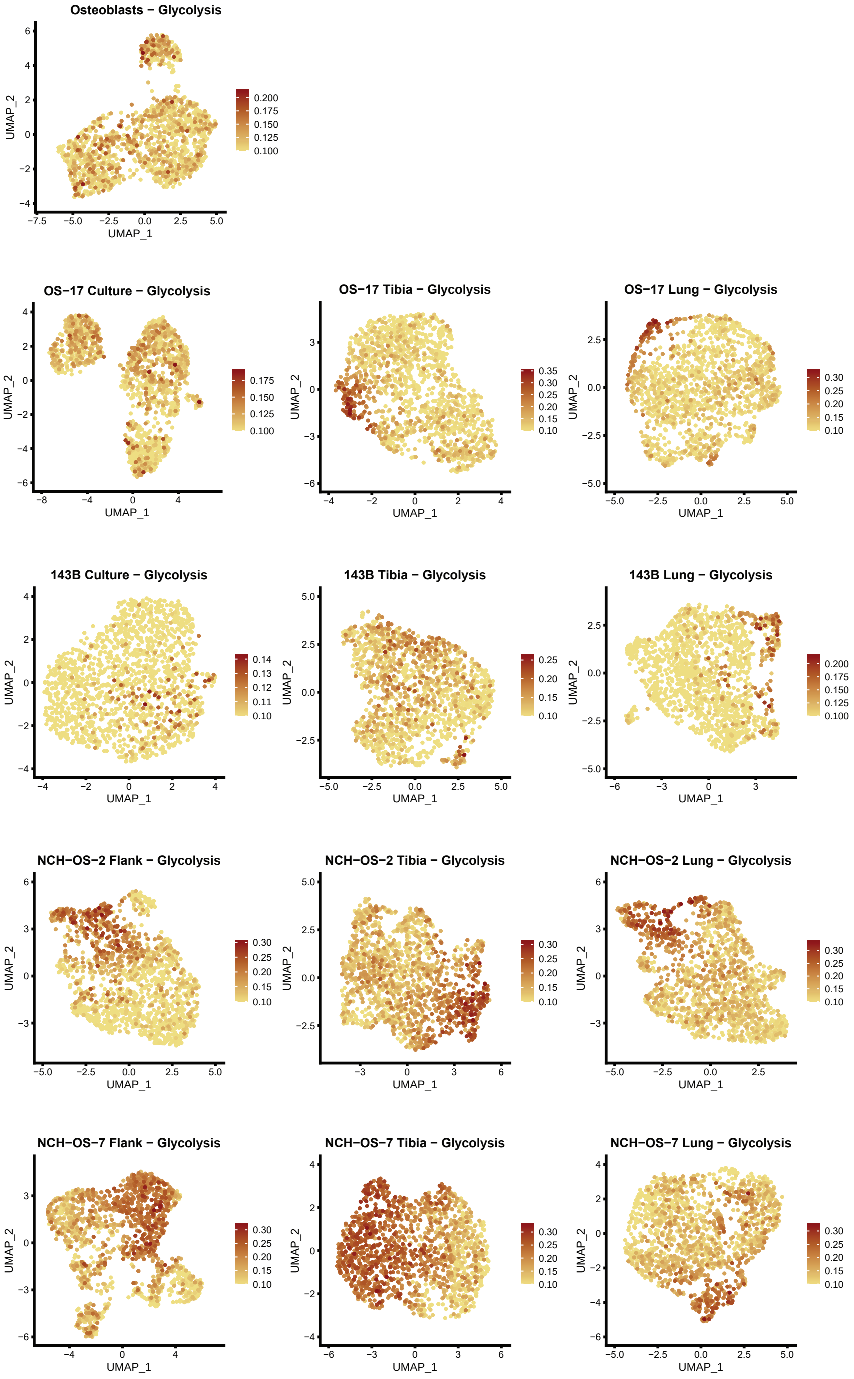


**Figure S9. Heterogeneous activation of glycolysis identified across multiple cell line and PDX tumor datasets.** FeaturePlots for glycolysis module score in each of the datasets. Module score for glycolysis was calculated using the 'AddModuleScore' function in Seurat with msigdb HALLMARK_GLYCOLYSIS genes as input features for the expression program.

Figure S10

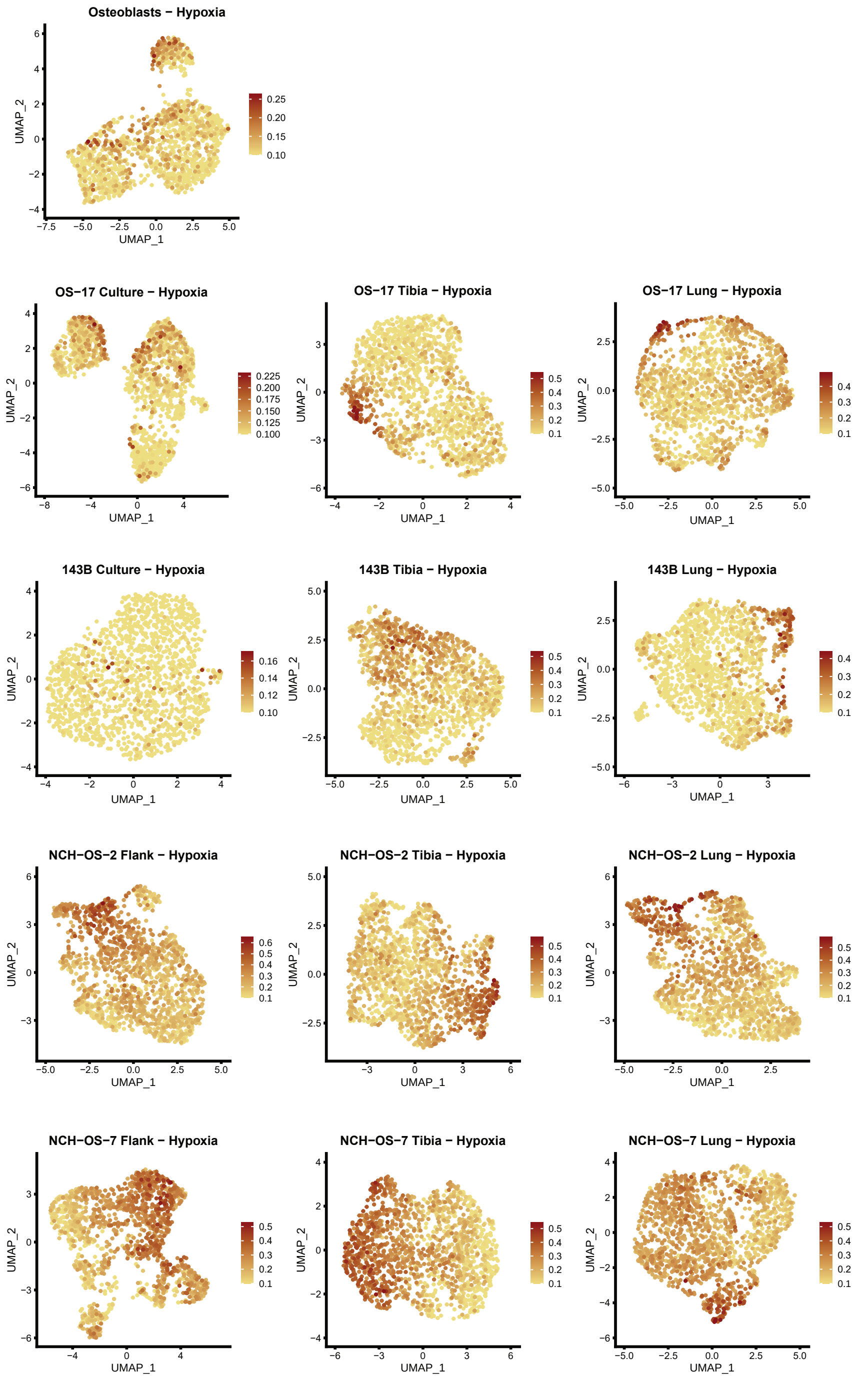


**Figure S10. Heterogeneous activation of hypoxia related genes identified across multiple cell line and PDX tumor datasets.** FeaturePlots for hypoxia module score in each of the datasets. Module score for hypoxia was calculated using the ‘AddModuleScore’ function in Seurat with msigdb HALLMARK_HYPOXIA genes as input features for the expression program.

Figure S11

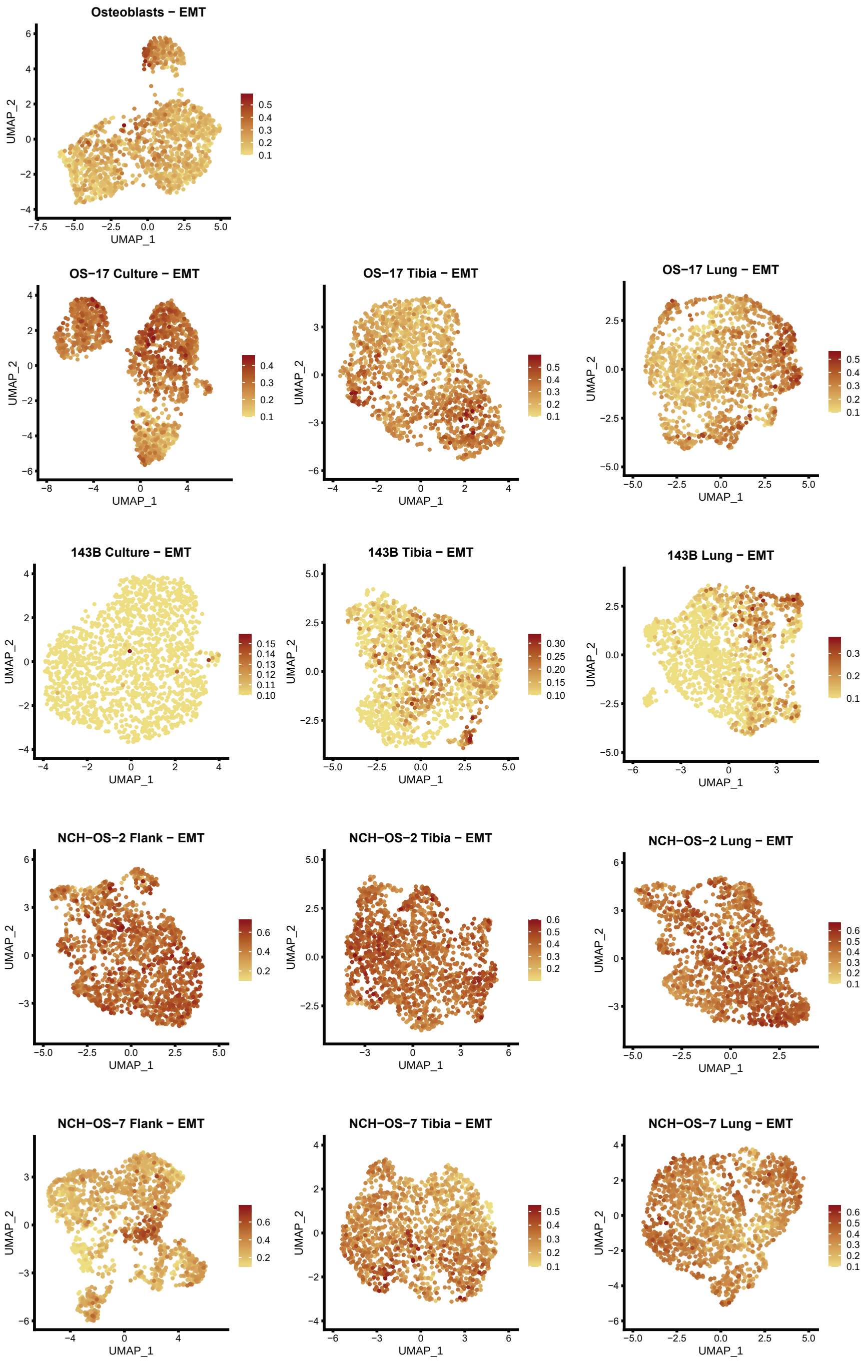


**Figure S11. Heterogeneous activation of EMT identified across multiple cell line and PDX tumor datasets.** FeaturePlots for EMT module score in each of the datasets. Module score for EMT was calculated using the 'AddModuleScore' function in Seurat with msigdb HALLMARK_EMT genes as input features for the expression program.

Figure S12

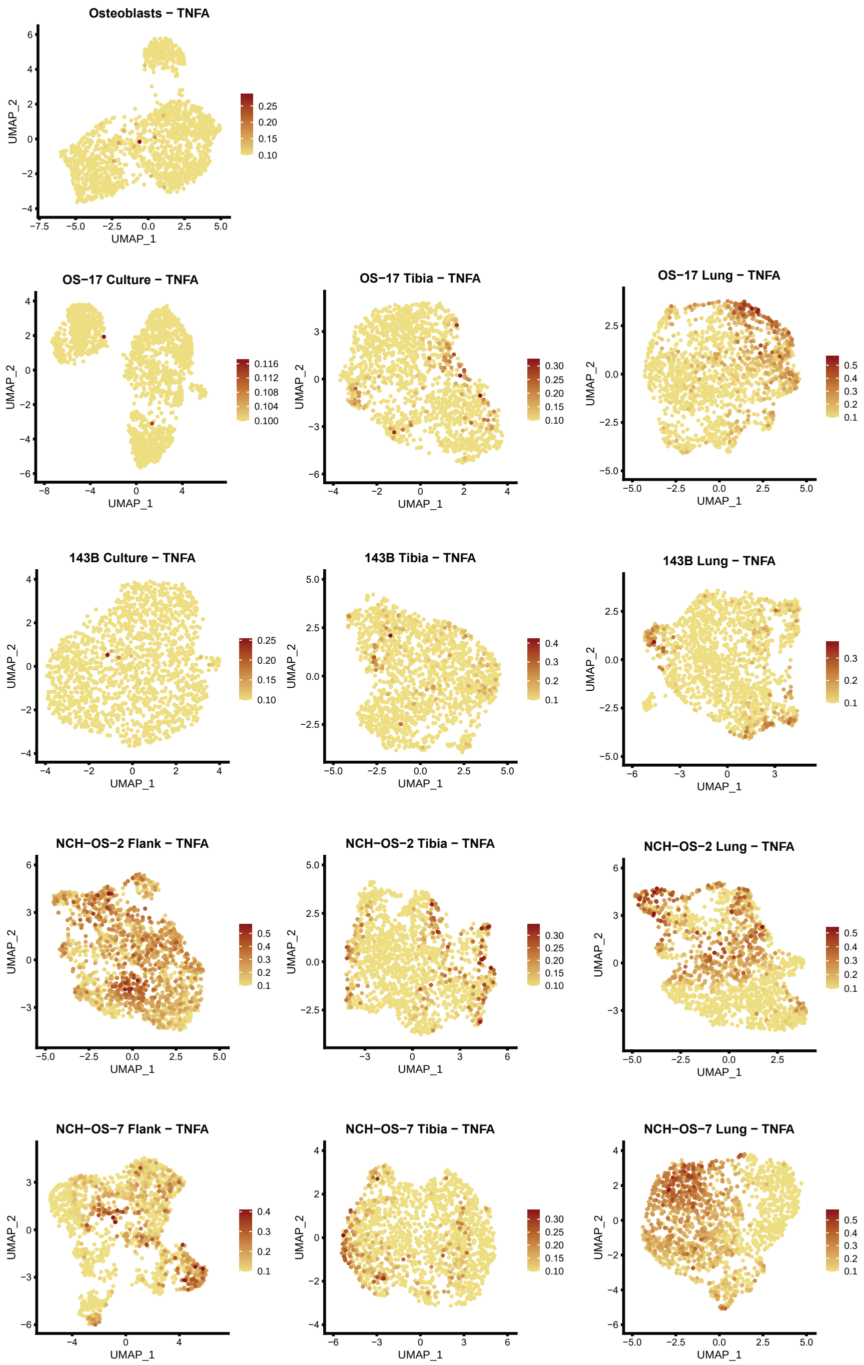


**Figure S12. Heterogeneous activation of ‘TNFα signaling via NFκB’ identified across multiple cell line and PDX tumor datasets.** FeaturePlots for TNFα signaling via NFκB module score in each of the datasets. Module score for EMT was calculated using the ‘AddModuleScore’ function in Seurat with msigdb HALL-MARK_TNFA genes as input features for the expression program.

Figure S13

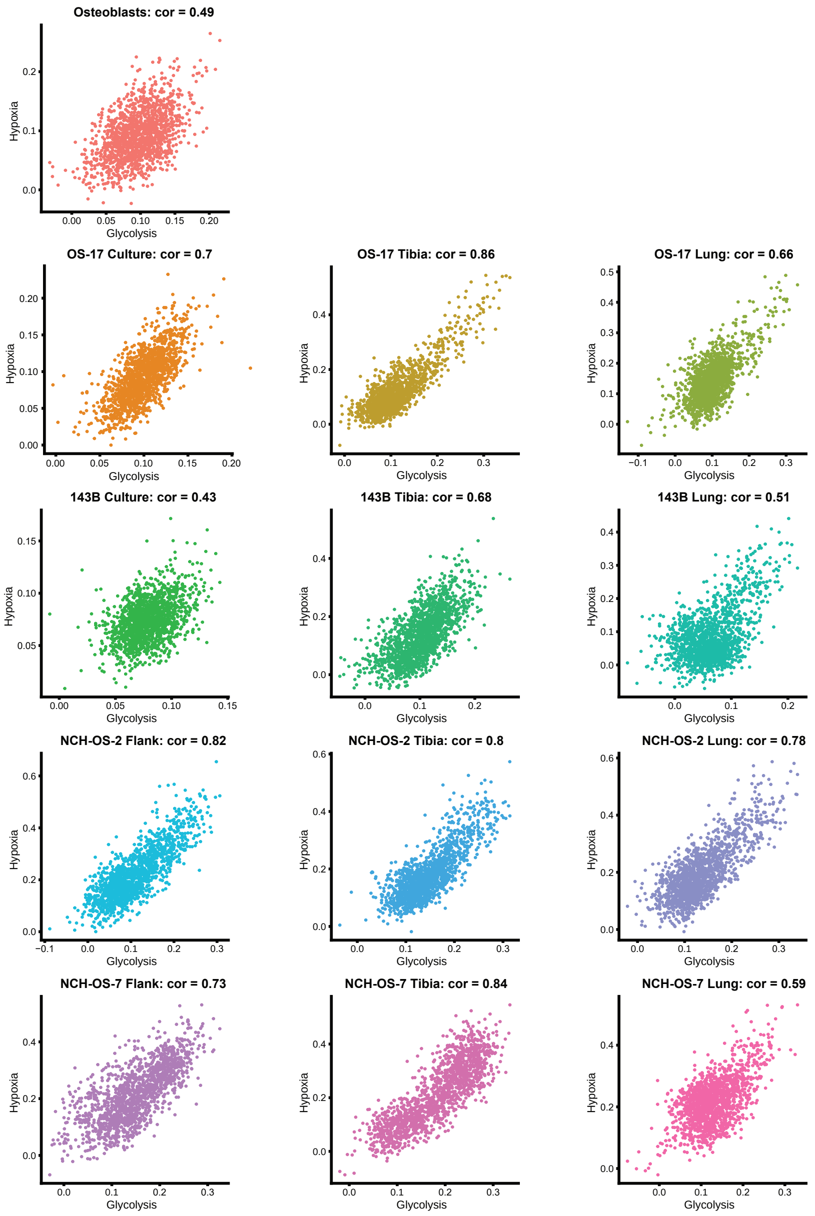


**Figure S13. Correlation analysis for multiple cell line and PDX tumor datasets reveals strong positive correlation between glycolysis and hypoxia module scores.** Scatter plots using the ‘FeatureScatter’ function in Seurat for glycolysis and hypoxia module scores in each of the datasets. Seurat-calculated correlation coefficient for each analysis is shown in the title above each plot.

Figure S14

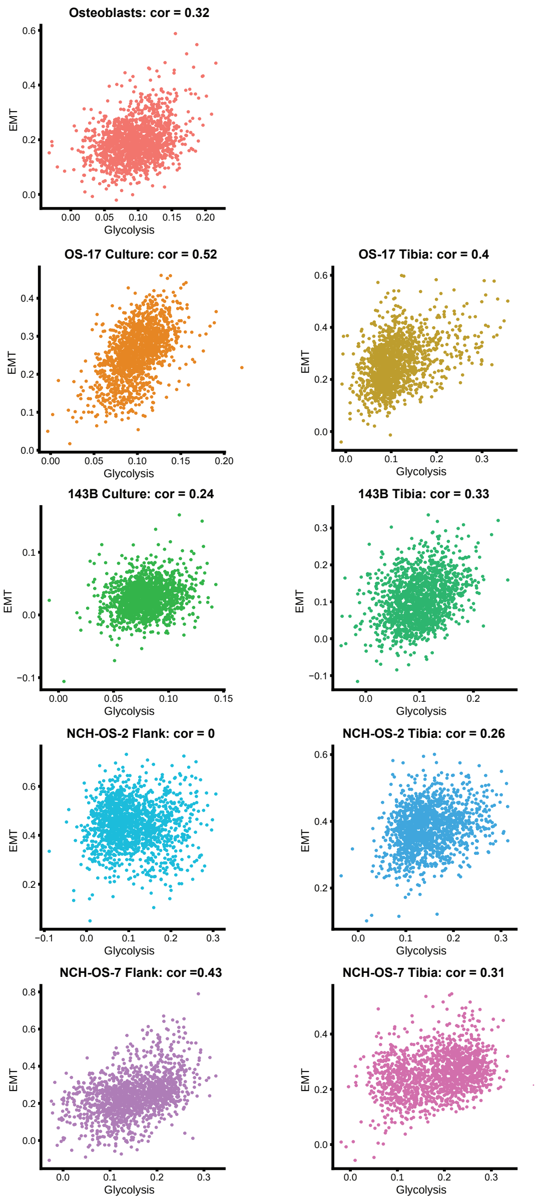


**Figure S14. Correlation analysis for multiple cell line and PDX tumor datasets reveals minimal correlation between glycolysis and EMT module scores.** Scatter plots using the 'FeatureScatter' function in Seurat for glycolysis and EMT module scores in each of the datasets. Seurat-calculated correlation coefficient for each analysis is shown in the title above each plot.

Figure S15

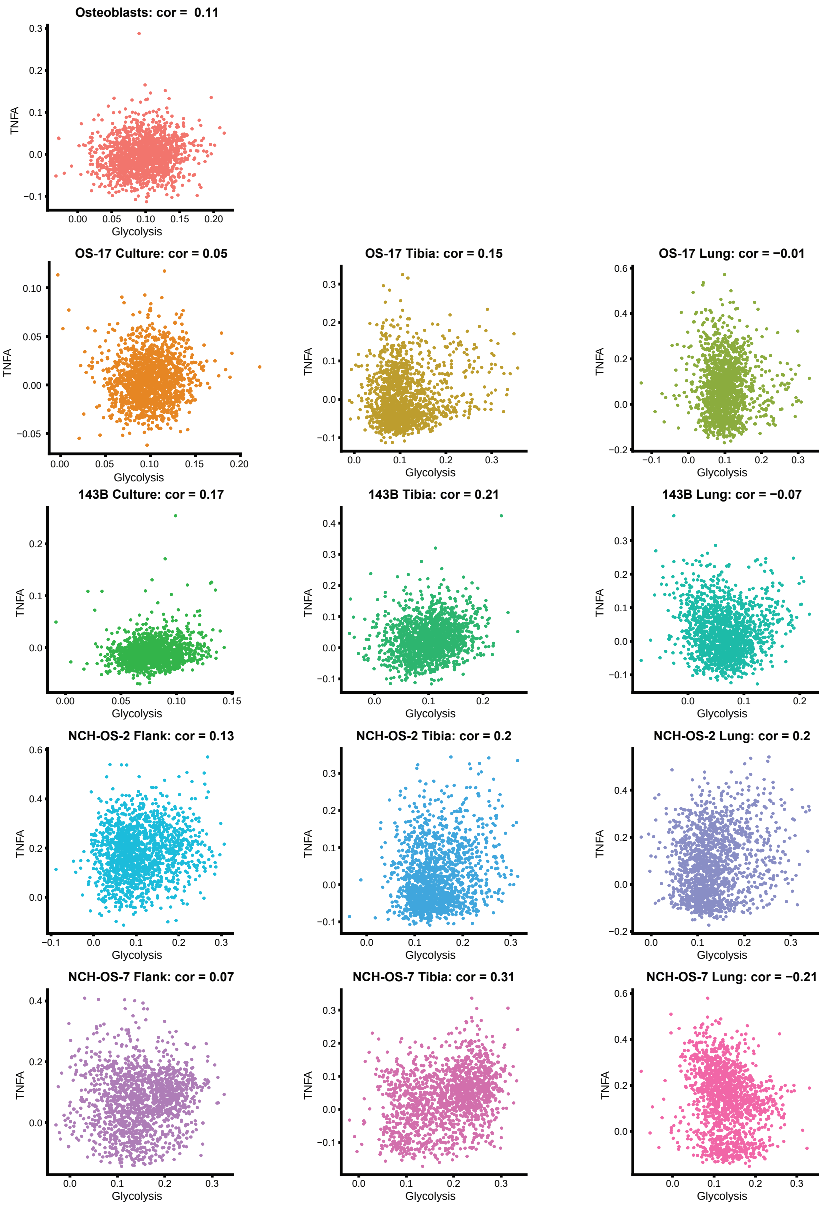


**Figure S15. Correlation analysis for multiple cell line and PDX tumor datasets reveals minimal correlation between glycolysis and 'TNFα signaling via NFκB' module scores.** Scatter plots using the 'FeatureScatter' function in Seurat for glycolysis and TNFα signaling via NFκB module scores in each of the datasets. Seurat-calculated correlation coefficient for each analysis is shown in the title above each plot.

Figure S16

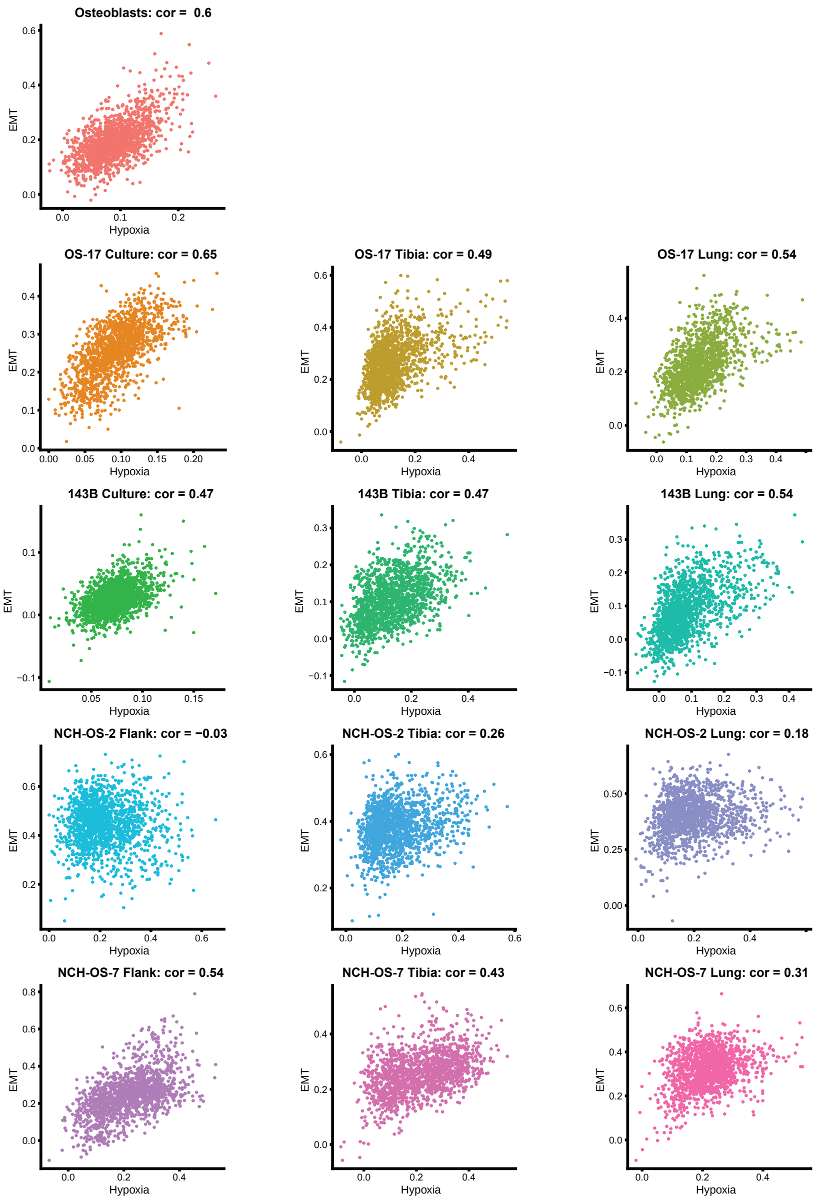


**Figure S16. Correlation analysis for multiple cell line and PDX tumor datasets reveals mixed correlation between hypoxia and EMT module scores.** Scatter plots using the 'FeatureScatter' function in Seurat for hypoxia and EMT module scores in each of the datasets. Seurat-calculated correlation coefficient for each analysis is shown in the title above each plot.

Figure S17

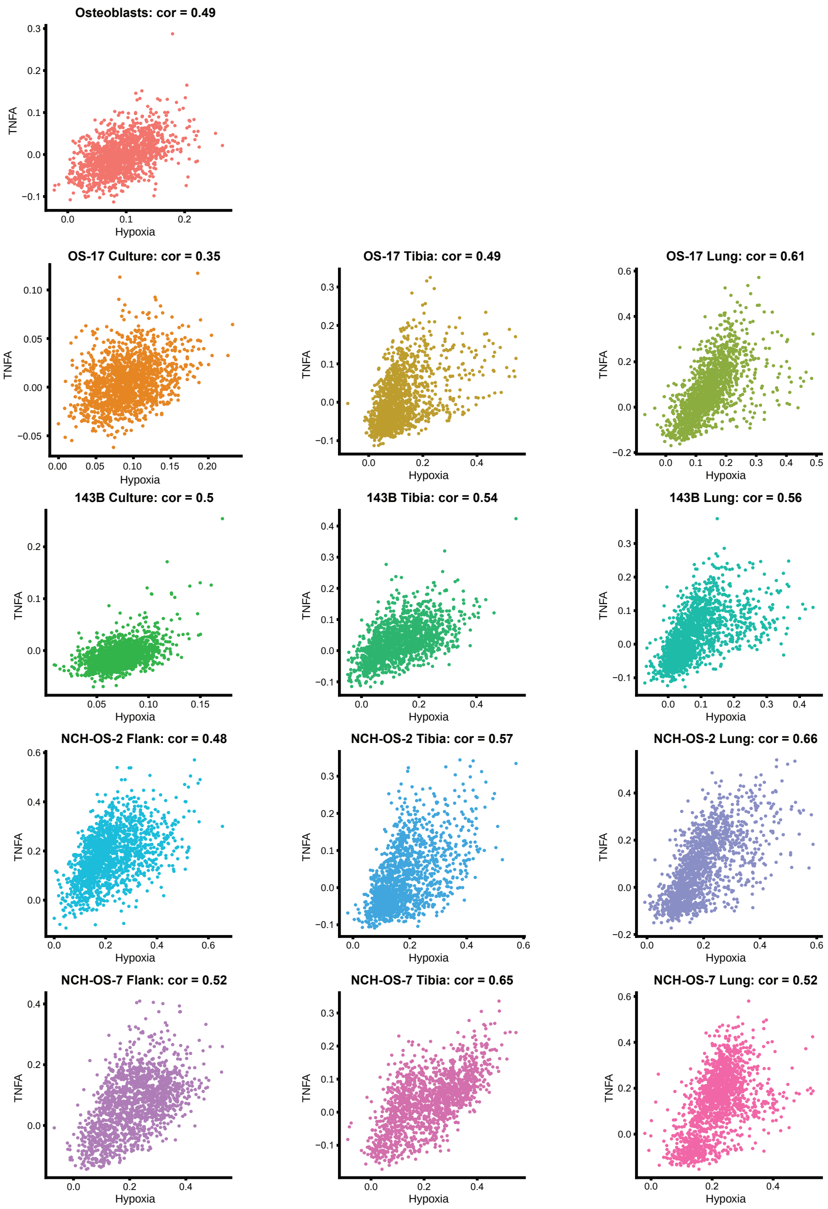


**Figure S17. Correlation analysis for multiple cell line and PDX tumor datasets reveals strong correlation between hypoxia and 'TNFα signaling via NFκB' module scores.** Scatter plots using the 'FeatureScatter' function in Seurat for hypoxia and TNFα signaling via NFκB module scores in each of the datasets. Seurat-calculated correlation coefficient for each analysis is shown in the title above each plot.

Figure S18

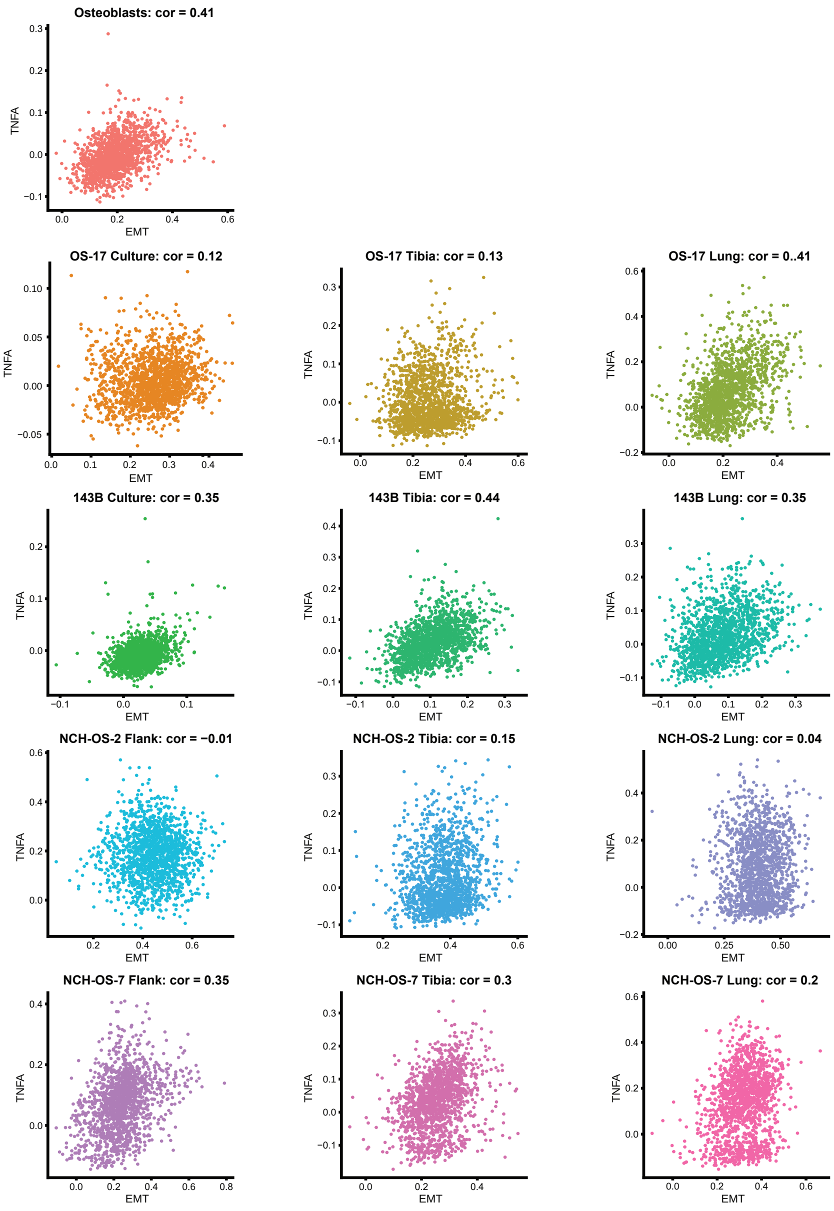


**Figure S18. Correlation analysis for multiple cell line and PDX tumor datasets reveals moderate correlation between EMT and 'TNFα signaling via NFκB' module scores.** Scatter plots using the 'FeatureScatter' function in Seurat for EMT and TNFα signaling via NFκB module scores in each of the datasets. Seurat-calculated correlation coefficient for each analysis is shown in the title above each plot.

Figure S19

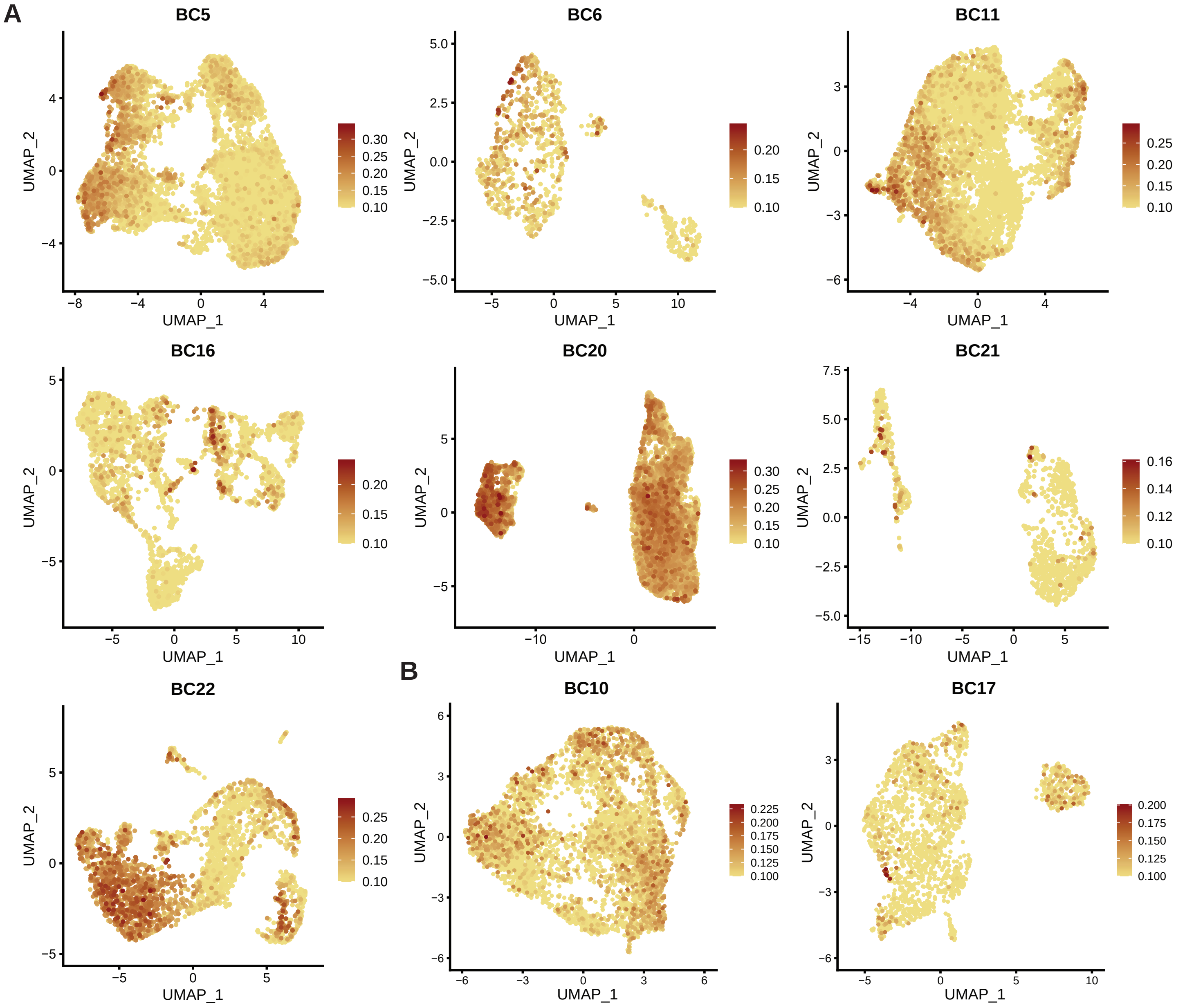


**Figure S19. Heterogeneous activation of glycolysis identified across multiple patient primary and metastatic tumor datasets.** A) Primary and B) metastatic patient tumor FeaturePlots showing glycolysis module score in each of the datasets. Module score for glycolysis was calculated using the 'AddModuleScore' function in Seurat with msigdb HALLMARK_GLYCOLYSIS genes as input features for the expression program. Patient data is from GSE152048.

Figure S20

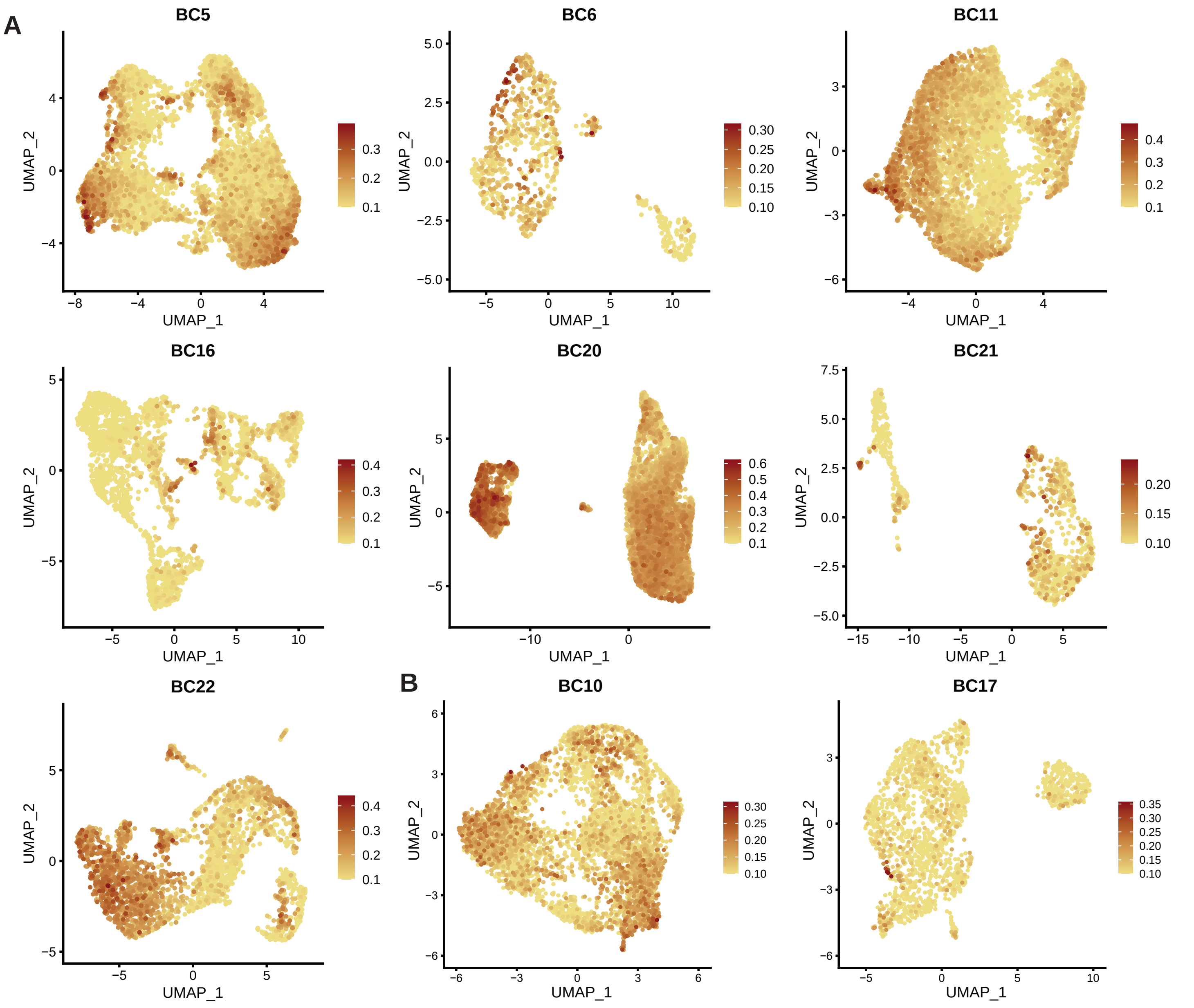


**Figure S20. Heterogeneous activation of hypoxia related genes identified across multiple patient primary and metastatic tumor datasets.** A) Primary and B) metastatic patient tumor FeaturePlots showing hypoxia module score in each of the datasets. Module score for hypoxia was calculated using the 'AddModule-Score' function in Seurat with msigdb HALLMARK_HYPOXIA genes as input features for the expression program. Patient data is from GSE152048.

Figure S21

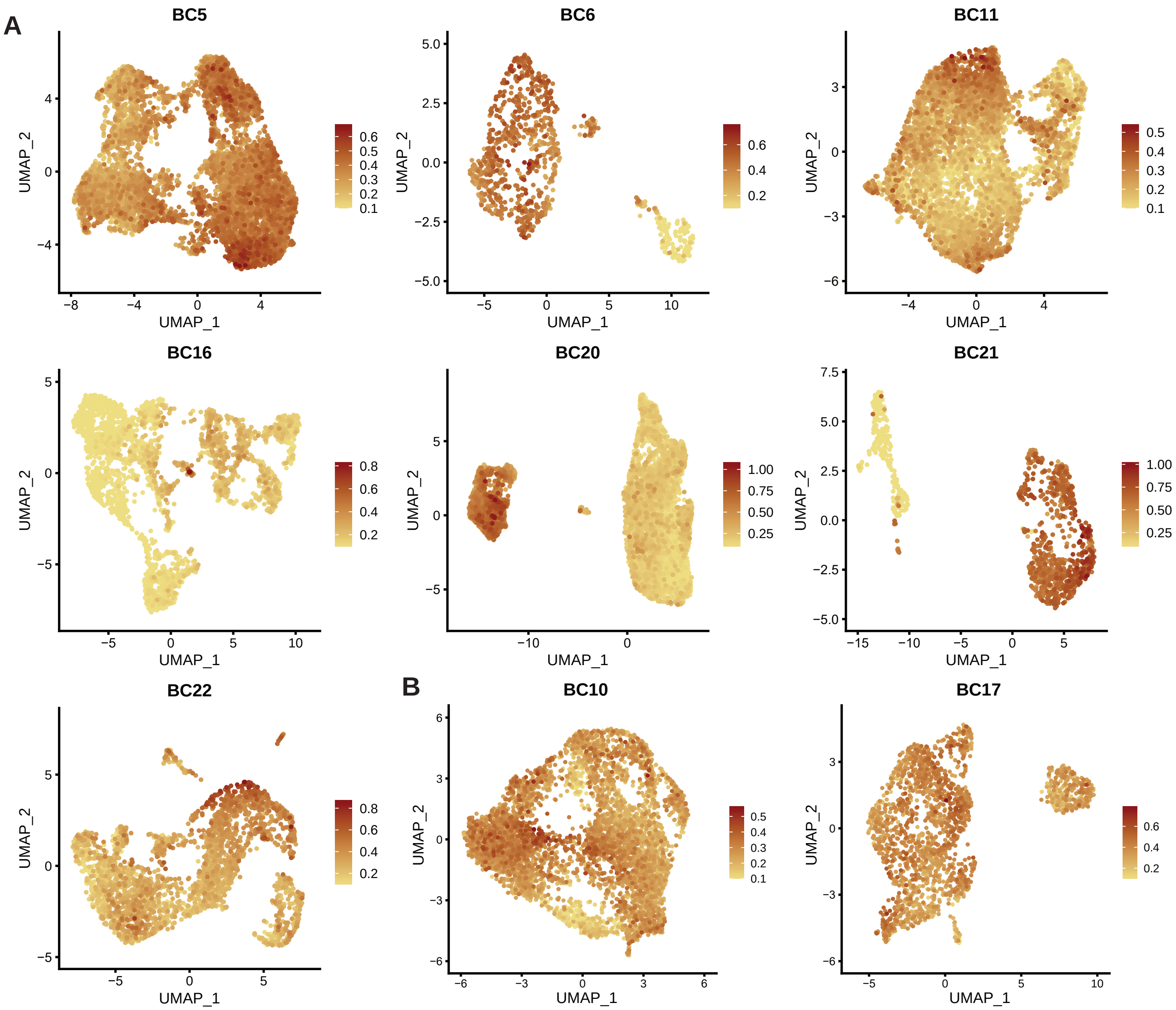


**Figure S21. Heterogeneous activation of EMT related genes identified across multiple patient primary and metastatic tumor datasets.** A) Primary and B) metastatic patient tumor FeaturePlots showing EMT module score in each of the datasets. Module score for EMT was calculated using the ‘AddModuleScore’ function in Seurat with msigdb HALLMARK_EPITHELIAL_MESENCHYMAL_TRANSITION genes as input features for the expression program. Patient data is from GSE152048.

Figure S22

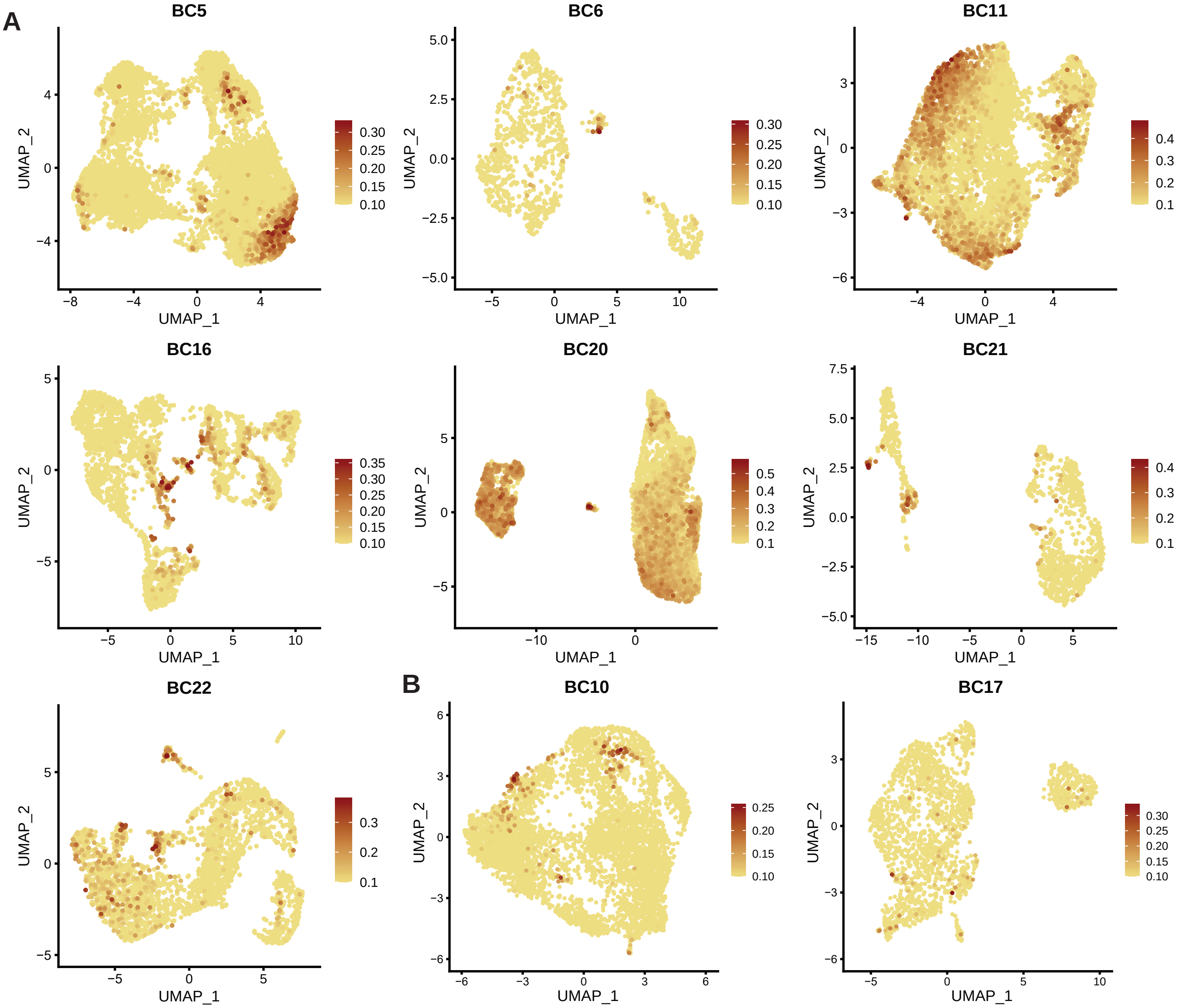


**Figure S22. Heterogeneous activation of 'TNFα signaling via NFκB' related genes identified across multiple patient primary and metastatic tumor datasets.** A) Primary and B) metastatic patient tumor FeaturePlots showing TNFα signaling via NFκB module score in each of the datasets. Module score for TNFα signaling via NFκB was calculated using the 'AddModuleScore' function in Seurat with msigdb HALLMARK_TNFA_SIGNALING_VIA_NFKB genes as input features for the expression program. Patient data is from GSE152048.

Figure S23

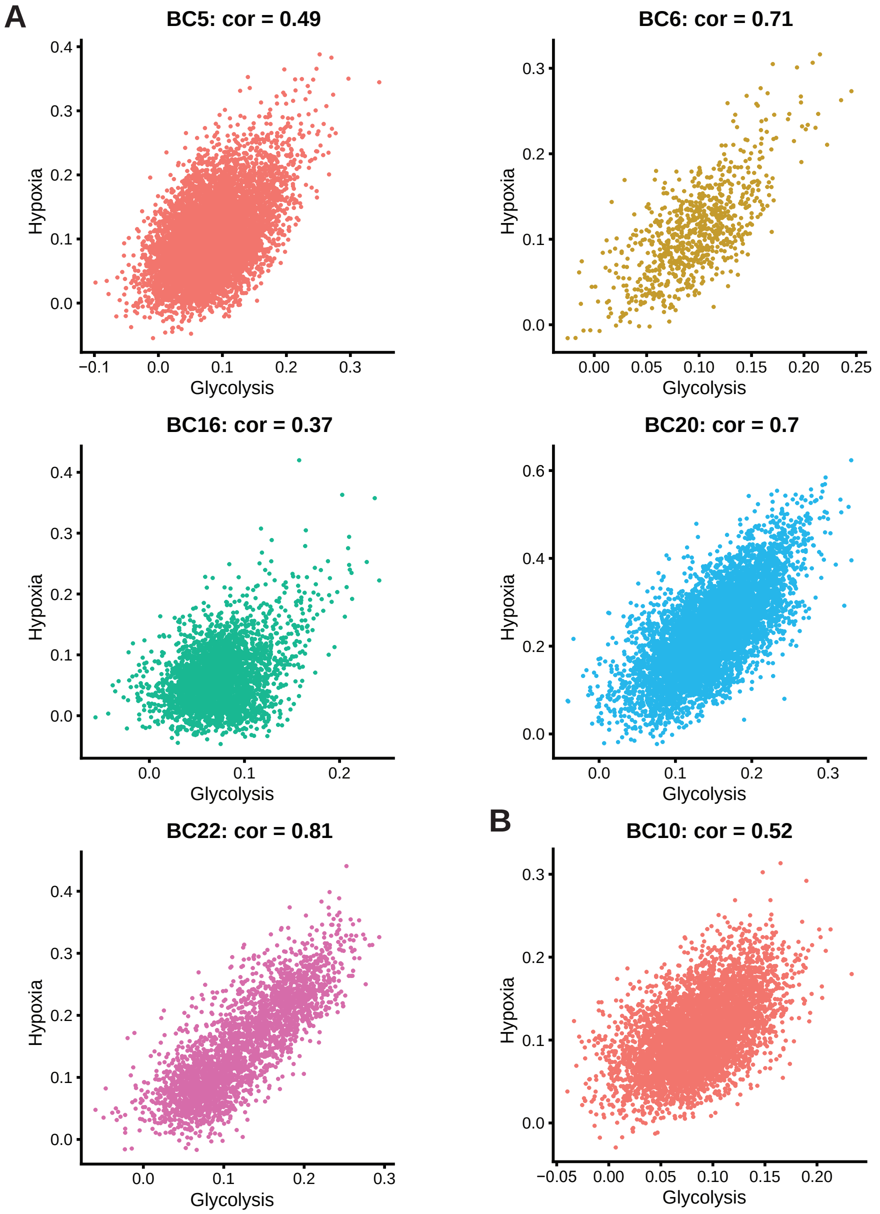


**Figure S23. Correlation analysis for multiple patient primary and metastatic tumor datasets reveals strong correlation between glycolysis and hypoxia module scores.** A) Primary and B) metastatic patient tumor scatter plots using the ‘FeatureScatter’ function in Seurat for glycolysis and hypoxia module scores in each of the datasets. Seurat-calculated correlation coefficient for each analysis is shown in the title above each plot. Patient data is from GSE152048.

Figure S24

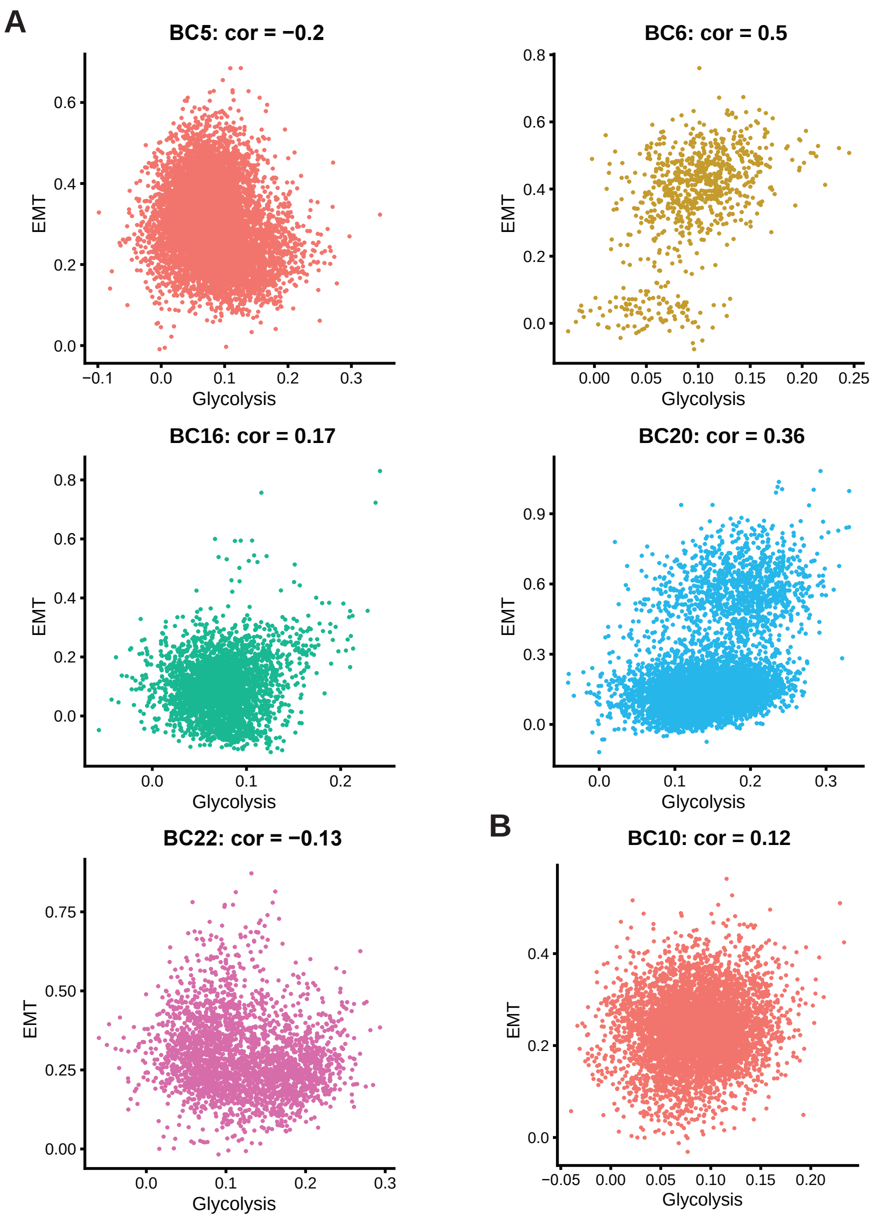


**Figure S24. Correlation analysis for multiple patient primary and metastatic tumor datasets reveals minimal correlation between glycolysis and EMT module scores.** A) Primary and B) metastatic patient tumor scatter plots using the 'FeatureScatter' function in Seurat for glycolysis and EMT module scores in each of the datasets. Seurat-calculated correlation coefficient for each analysis is shown in the title above each plot. Patient data is from GSE152048.

Figure S25

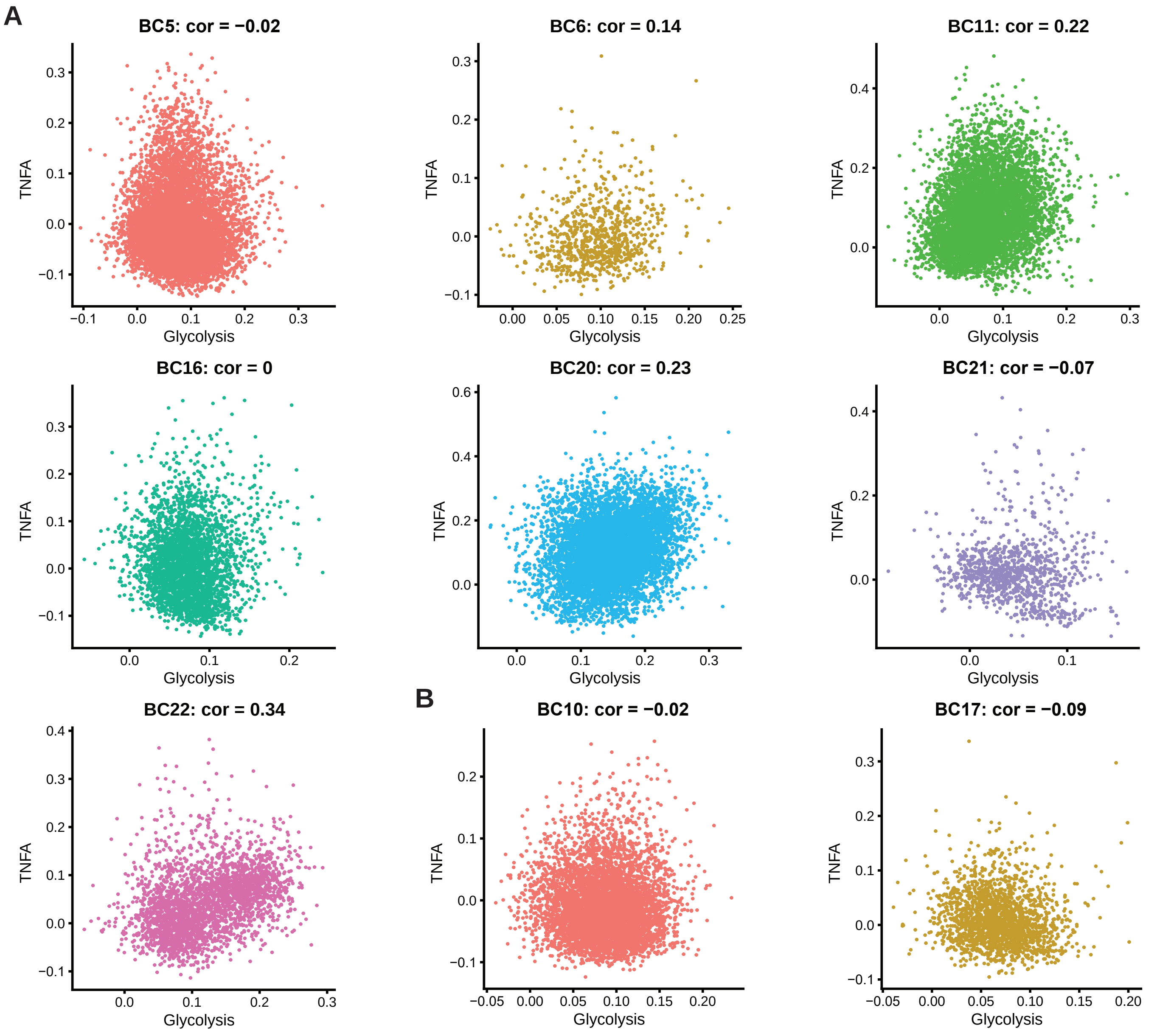


**Figure S25. Correlation analysis for multiple patient primary and metastatic tumor datasets reveals minimal correlation between glycolysis and 'TNFα signaling via NFκB' module scores.** A) Primary and B) metastatic patient tumor scatter plots using the 'FeatureScatter' function in Seurat for glycolysis and TNFα signaling via NFκB module scores in each of the datasets. Seurat-calculated correlation coefficient for each analysis is shown in the title above each plot. Patient data is from GSE152048.

Figure S26

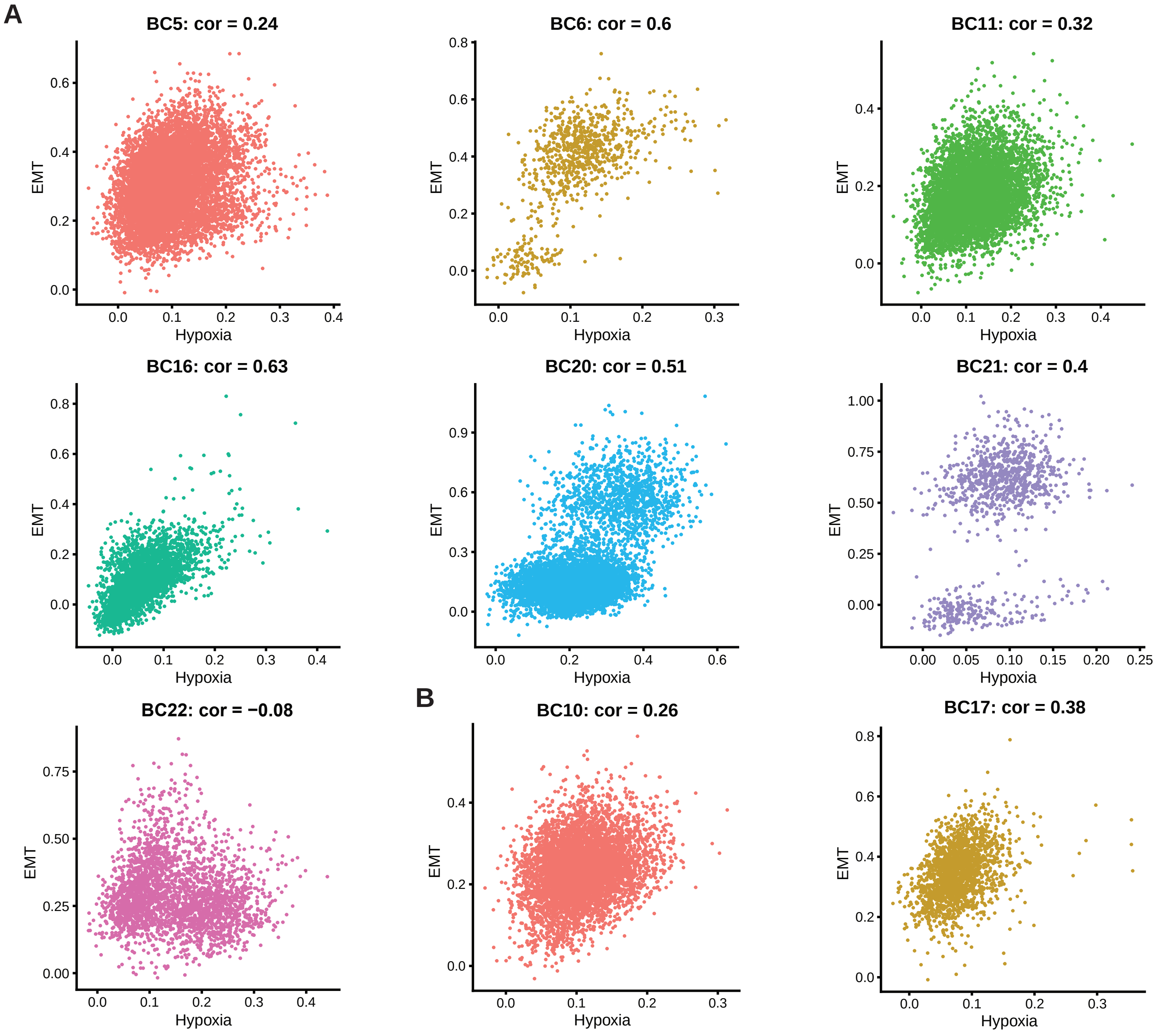


**Figure S26. Correlation analysis for multiple patient primary and metastatic tumor datasets reveals mixed correlation between hypoxia and EMT module scores.** A) Primary and B) metastatic patient tumor scatter plots using the 'FeatureScatter' function in Seurat for hypoxia and EMT module scores in each of the datasets. Seurat-calculated correlation coefficient for each analysis is shown in the title above each plot. Patient data is from GSE152048.

Figure S27

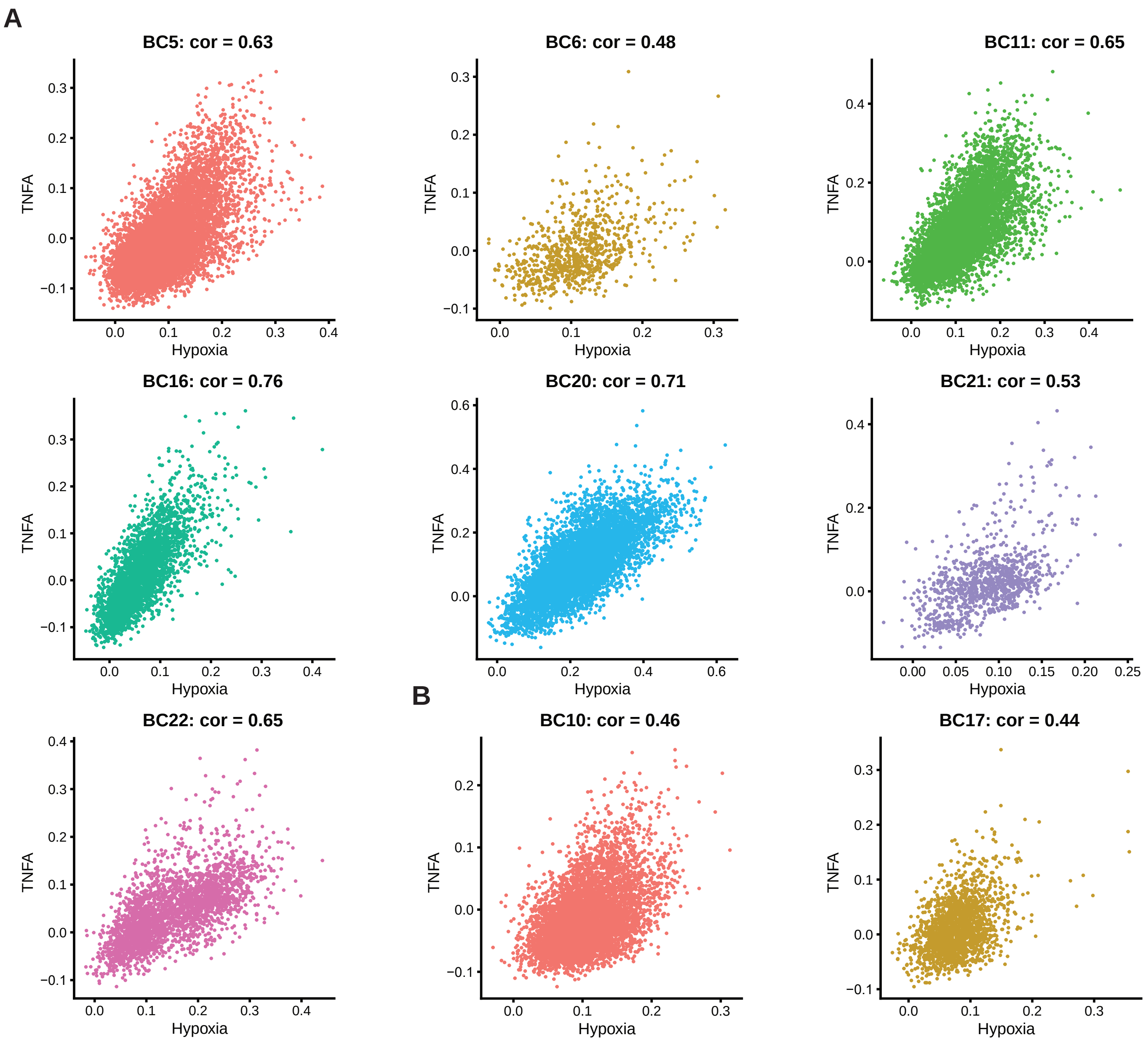


**Figure S27. Correlation analysis for multiple patient primary and metastatic tumor datasets reveals strong correlation between hypoxia and 'TNFα signaling via NFκB' module scores.** A) Primary and B) metastatic patient tumor scatter plots using the 'FeatureScatter' function in Seurat for hypoxia and TNFα signaling via NFκB module scores in each of the datasets. Seurat-calculated correlation coefficient for each analysis is shown in the title above each plot. Patient data is from GSE152048.

Figure S28

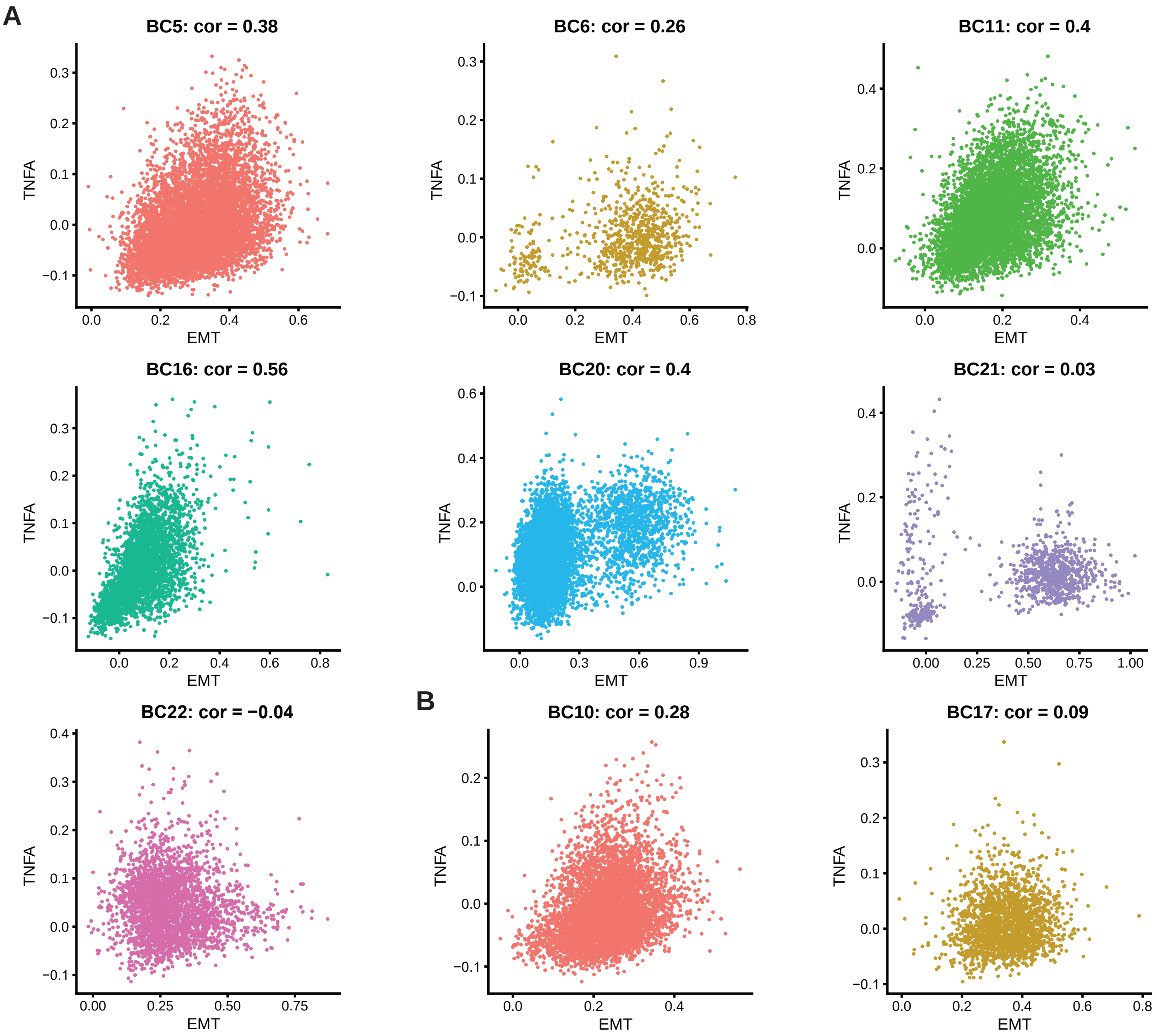


**Figure S28. Correlation analysis for multiple patient primary and metastatic tumor datasets reveals moderate correlation between EMT and 'TNFα signaling via NFκB' module scores.** A) Primary and B) metastatic patient tumor scatter plots using the 'FeatureScatter' function in Seurat for EMT and TNFα signaling via NFκB module scores in each of the datasets. Seurat-calculated correlation coefficient for each analysis is shown in the title above each plot. Patient data is from GSE152048.

Figure S30

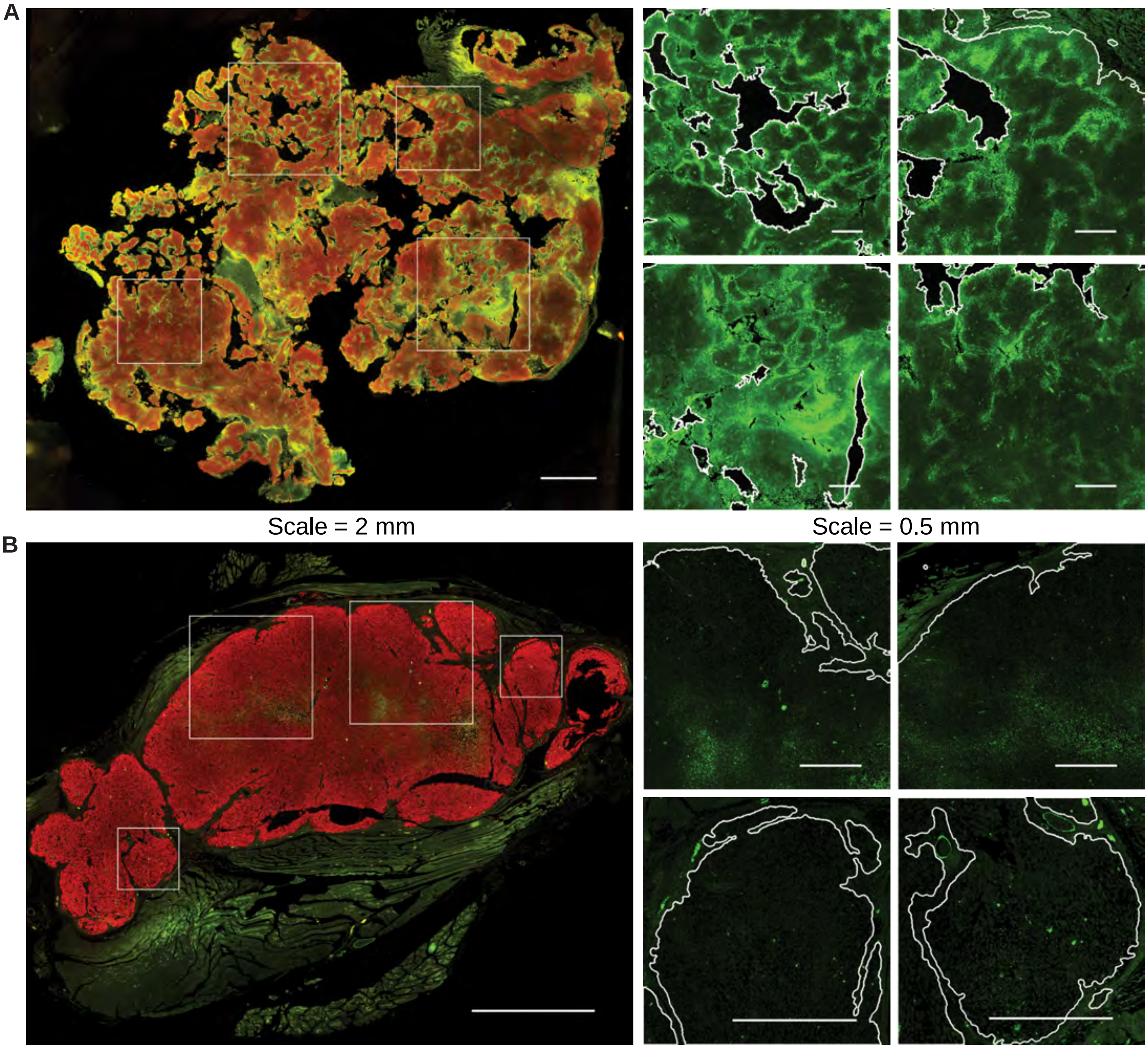


**Figure S30. Primary tumors demonstrate heterogeneity in GLUT1 staining.** A, B) Immunofluorescence staining of OS-17 and 143B primary tumors, respectively, for GLUT1 (green; a marker of glycolysis) and vimentin (red; marker to identify osteosarcoma cells). Magnified GLUT1 staining is shown for the boxed regions on the whole-section images. Lesion edges are indicated by white outlines in the magnified regions. Tumors showed a high degree of intra-tumor variation in GLUT1 staining intensity. Intensity ranged from strong to light in OS-17 and moderate to light in 143B, in which only some cells showed moderate expression of GLUT1.

Figure S29

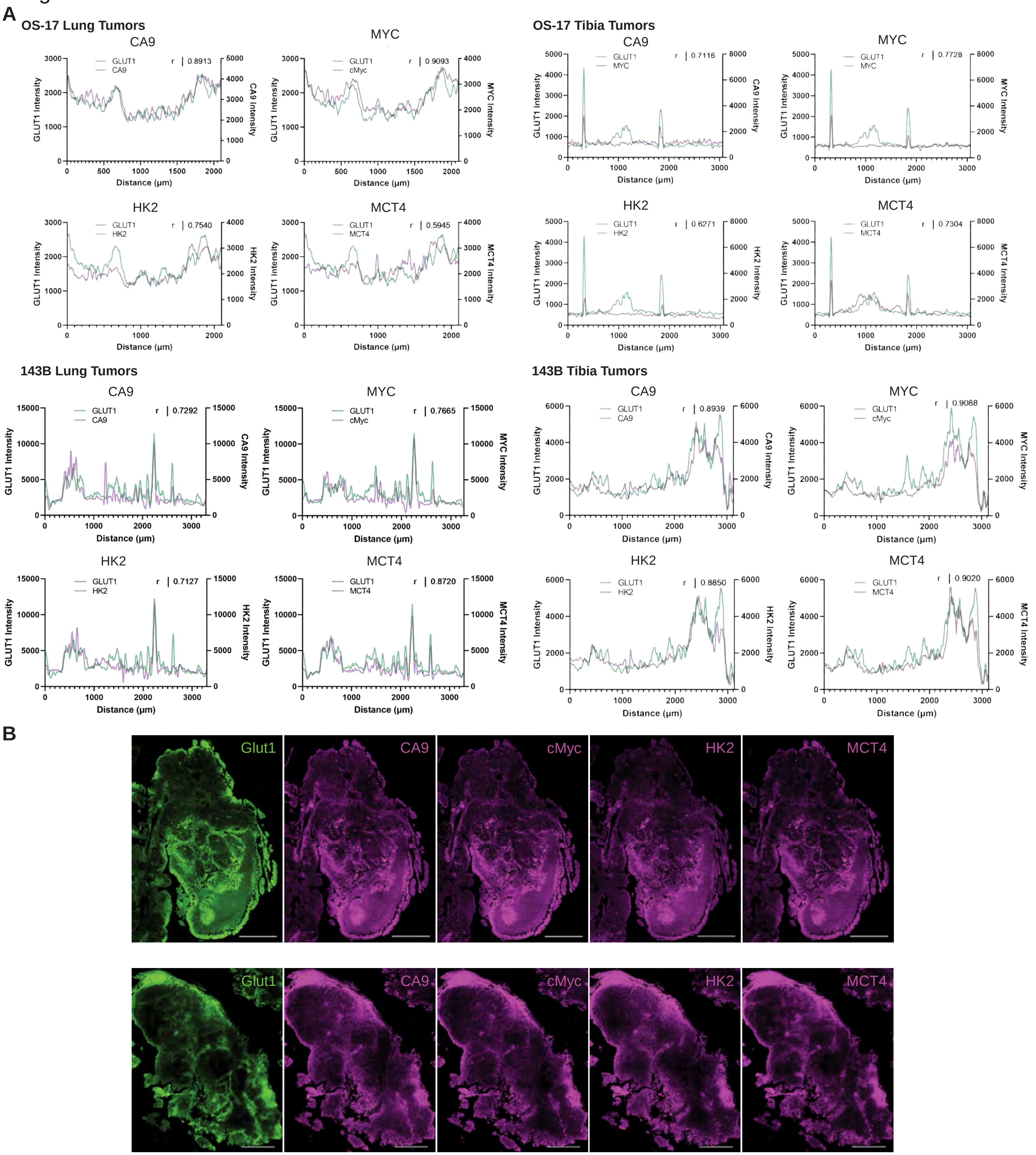


**Figure S29. High correlation between multiple markers of glycolysis activation in primary and metastatic osteosarcoma lesions.** A) Correlation of fluorescence intensity of markers of glycolysis (CA9, HK2, MCT4), an upstream regulator (cMYC) with GLUT1 along a ~3mm profile line drawn through the corresponding primary and metastatic lesions. The location and intensity of staining was highly correlated with GLUT1 for all markers. B) Fluorescent imaging of markers of glycolysis (CA9, HK2, MCT4) and an upstream regulator (cMYC) with GLUT1 along a ~1mm profile line drawn through the 143B lung lesions. The location and intensity of staining shows similarity to that of GLUT1 for all markers. Representative of n=3 mice.

Figure S31

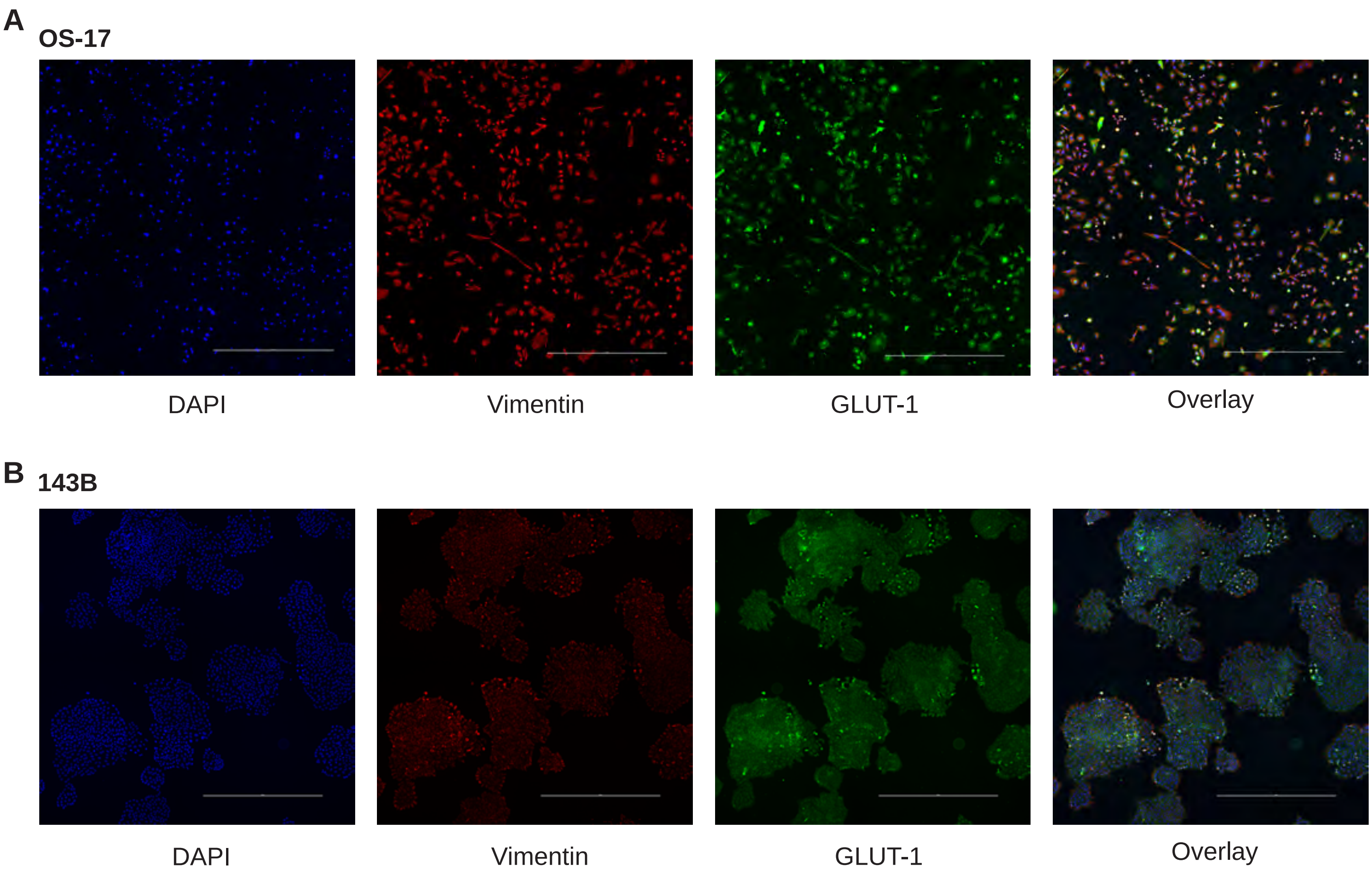


**Figure S31. Tumor cell cultures demonstrate heterogeneity in GLUT1 staining.** A, B) Immunofluorescence staining of OS-17 and 143B cell culture, respectively, for GLUT1 (green; a marker of glycolysis) and vimentin (red; marker to identify osteosarcoma cells) with scale bars of 1000 micrometers. Both cell culture models displayed strong to light expression of GLUT1.

Figure S32

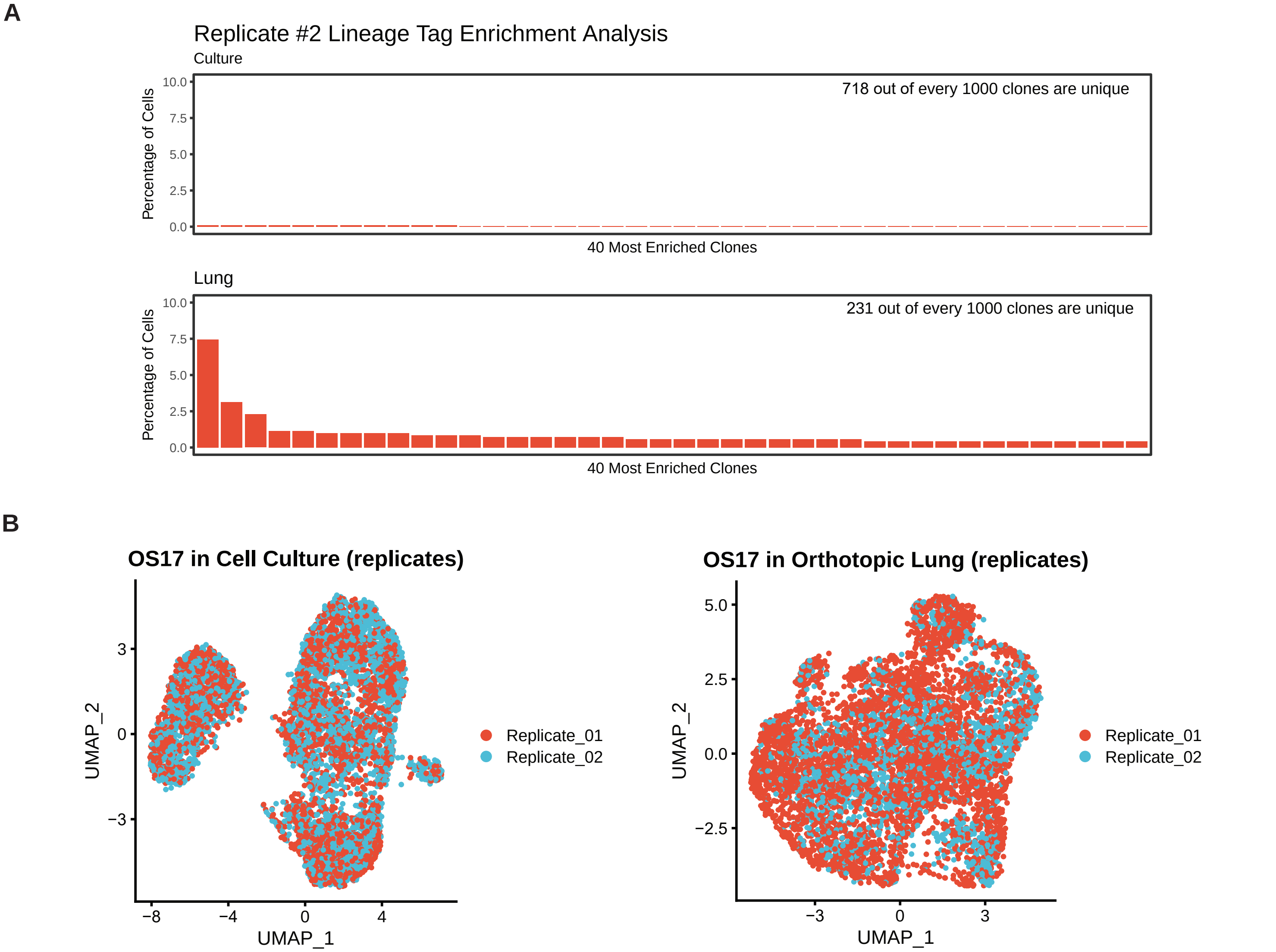


**Figure S32. Replicates demonstrate reproducible lineage tag enrichment profiles and consistent phenotypic profiles.** A) Frequency distribution of clones identified in each of the conditions generated using a biological replicate of the starting population (cells transduced with a separate batch of lentivirus library). B) Admixing of OS-17 cells from distinct, biological and technical replicates grown *in vitro* or as metastatic lesions suggests that intra-tumor transcriptional heterogeneity of osteosarcoma cells is not driven by individual tumor identity.

Figure S33

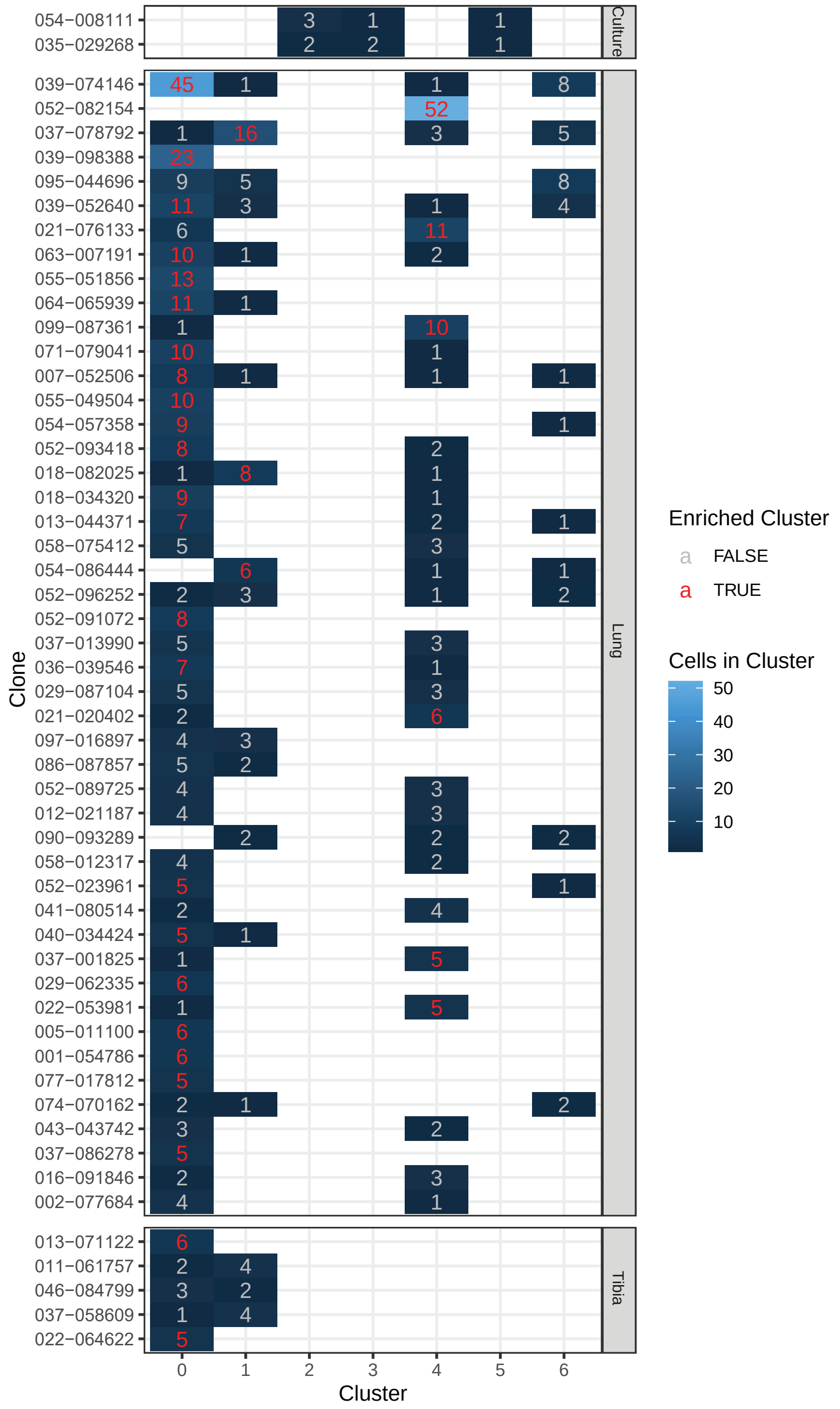


**Figure S33. Contingency table displaying the number of enriched clones per cluster in culture, tibia, and lung.** Numbers display how many cells from each clone are in a particular cluster from OS-17 tumors in the tibia or lung. Significant enrichment within one clone is indicated in red text, as calculated permutation analysis. The color of each box correlates to the total cells in each cluster.
