## Supplemental tables for "Osteosarcoma tumors maintain intra-tumoral transcriptional heterogeneity during bone and lung colonization"

Supplementary Tables

Table S1. Characteristics of Patient-Derived Tumor Xenograft Models used in this study

| **Module** | **Field** | **NCH-OS-2** | **NCH-OS-7** |
| --- | --- | --- | --- |
| **Clinical/patient** | Submitter patient ID | X0070 | X0140 |
|  | Gender | Male | Female |
|  | Age | 20-25 | 15-20 |
|  | Diagnosis | Metastatic osteosarcoma | Metastatic osteosarcoma |
|  | Consent to share data | Available to academic centers only | Yes |
|  | Ethnicity/race | Caucasian | Caucasian |
|  | Current treatment drug |  |  |
|  | Virology status | HepA(-), HepB(-), HepC(-), HIV-1(-), HIV-2(-), Mycoplasma(-) | HepA(-), HepB(-), HepC(-), HIV-1(-), HIV-2(-), Mycoplasma(-) |
| **Clinical/tumor** | Submitter tumor ID | X0070 | X0140 |
|  | Primary tumor tissue of origin | Bone | Bone |
|  | Primary, metastasis, recurrence | Metastasis | Recurrence |
|  | Specimen tumor tissue | Lung | Lung |
|  | Tumor grade; classification | High Grade | High grade |
|  | Disease stage; classification | metastatic osteosarcoma | metastatic osteosarcoma |
|  | Specific markers (diagnostic linked); platform | N/A | N/A |
|  | Is tumor from untreated patient? | Yes | No |
|  | Original tumor sample type | Surgical resection | Open biopsy |
|  | Tumor from an existing PDX model? ID? Why sub-line? | No | No |
| **Model creation** | Submitter PDX ID | NCH-OS-2 | NCH-OS-7 |
|  | Mouse strain (and source) | C.B-17/IcrHan Hsd-Prkdcscid | C.B-17/IcrHan Hsd-Prkdcscid |
|  | Strain immune system humanized? | No | No |
|  | Type of humanization | NA | NA |
|  | Tumor preparation | Solid tumor fragments and cell suspension | Solid tumor fragments and cell suspension |
|  | Injection type and site | Subcutaneous, flank; Intravenous, tail vien; Intraosseous, tibial plate | Subcutaneous, flank; Intravenous, tail vien; Intraosseous, tibial plate |
| **Model quality assurance** | Tumor characterization technology | Histology and IHC | Histology and IHC |
|  | Tumor confirmed not to be of mouse/EBV origin | 84% Human tissue | 76% Human tissue |
|  | Passage QA performed | Passage P3 | Passage P5 |
| **Model study** | Treatment, passage | No previous treatment | No previous treatment |
| **Associated metadata** | PDX model availability? | Yes, frozen tumor | Yes, frozen tumor |
|  | Governance restriction for distribution | Available to academic centers only | Available to academic centers only |

Table S2. Relative percentage of cells distributed between clusters in each model between Tibia and Lung colonization conditions

| OS-17 | Condition | Cluster | Relative percentage |
| --- | --- | --- | --- |
|  | Tibia | 0 | 50.26325 |
|  | Tibia | 1 | 43.45384 |
|  | Tibia | 2 | 1.053001 |
|  | Tibia | 3 | 5.229905 |
|  | Lung | 0 | 46.57775 |
|  | Lung | 1 | 10.17901 |
|  | Lung | 2 | 39.69814 |
|  | Lung | 3 | 3.545104 |
| 143B | Tibia | 0 | 97.89638 |
|  | Tibia | 1 | 1.149201 |
|  | Tibia | 2 | 0.447994 |
|  | Tibia | 3 | 0.506428 |
|  | Lung | 0 | 2.220491 |
|  | Lung | 1 | 92.30619 |
|  | Lung | 2 | 3.350214 |
|  | Lung | 3 | 2.123101 |
| NCH-OS-2 | Tibia | 0 | 82.28155 |
|  | Tibia | 1 | 5.339806 |
|  | Tibia | 2 | 12.37864 |
|  | Lung | 0 | 20.32767 |
|  | Lung | 1 | 71.48058 |
|  | Lung | 2 | 8.191748 |
| NCH-OS-2 | Tibia | 0 | 0 |
|  | Tibia | 1 | 59.4258 |
|  | Tibia | 2 | 40.5742 |
|  | Tibia | 3 | 0 |
|  | Lung | 0 | 71.01349 |
|  | Lung | 1 | 1.383604 |
|  | Lung | 2 | 1.176064 |
|  | Lung | 3 | 26.42684 |

Table S3. Table showing differentially upregulated genes in the top ten enriched clonal families relative to the remaining metastatic OS-17 cells

|  | P value | Average log2(fold-change) | Adjusted p value |
| --- | --- | --- | --- |
| PTPRZ1 | 3.49E-24 | 0.340175 | 7.04E-20 |
| CD9 | 9.46E-21 | 0.522326 | 1.91E-16 |
| GZMB | 3.52E-20 | 0.360582 | 7.1E-16 |
| ENO2 | 8.76E-16 | 0.516978 | 1.77E-11 |
| AQP1 | 1.84E-15 | 0.298789 | 3.71E-11 |
| CDC42EP3 | 2.58E-14 | 0.302923 | 5.2E-10 |
| P4HA2 | 1.1E-12 | 0.515958 | 2.21E-08 |
| TGFBI | 1.57E-12 | 0.522179 | 3.16E-08 |
| COL6A2 | 3.66E-12 | 0.456208 | 7.37E-08 |
| BEX1 | 9.34E-12 | 0.252685 | 1.88E-07 |
| WDR54 | 1.21E-11 | 0.264337 | 2.44E-07 |
| S100A10 | 6.43E-11 | 0.308953 | 1.3E-06 |
| MIF | 7.32E-10 | 0.370393 | 1.47E-05 |
| TPI1 | 4.65E-09 | 0.285224 | 9.38E-05 |
| ENO1 | 6.68E-09 | 0.468255 | 0.000135 |
| VEGFA | 2.77E-08 | 0.752467 | 0.000558 |
| FLNB | 2.92E-08 | 0.260708 | 0.000589 |
| COL6A1 | 1.18E-07 | 0.287235 | 0.002379 |
| PGK1 | 2.51E-07 | 0.609942 | 0.005059 |
| S100A2 | 3.03E-07 | 0.421746 | 0.006097 |
| GBE1 | 3.05E-07 | 0.318568 | 0.006149 |
| SEC61G | 4.59E-07 | 0.258519 | 0.009259 |
| PFKP | 5.71E-07 | 0.331277 | 0.011517 |
| FAM162A | 5.76E-07 | 0.682216 | 0.011619 |
| BEND5 | 6.43E-07 | 0.251597 | 0.012955 |
| LGALS3 | 1.98E-06 | 0.342243 | 0.997 |

Table S4. Lineage tag distribution statistics for two biological replicates

| Replicate 1 | | |
| --- | --- | --- |
| Number of copies | Frequency | Percentage |
| 1 | 2283 | 69.4 |
| 2 | 659 | 20.0 |
| 3 | 217 | 6.6 |
| 4 | 76 | 2.3 |
| 5 | 32 | 1.0 |
| 6 | 10 | 0.3 |
| 7 | 9 | 0.3 |
| 11 | 1 | 0.0 |
| 13 | 1 | 0.0 |
| 14 | 1 | 0.0 |
| Replicate 2 | | |
| Number of copies | Frequency | Percentage |
| 1 | 482 | 84.0 |
| 2 | 69 | 12.0 |
| 3 | 15 | 2.6 |
| 4 | 6 | 1.0 |
| 5 | 2 | 0.3 |

Table S5. Lineage tag summary statistics for two biological replicates

| Replicate 1 | |
| --- | --- |
| Summary Statistics | |
| Min. | 1 |
| 1stQu. | 1 |
| Median | 1 |
| Mean | 1.483 |
| 3rdQu. | 2 |
| Max. | 14 |
| Replicate 2 | |
| Summary Statistics | |
| Min. | 1 |
| 1stQu. | 1 |
| Median | 1 |
| Mean | 1.218 |
| 3rdQu. | 1 |
| Max. | 5 |
